# Extrachromosomal DNA Gives Cancer a New Evolutionary Pathway

**DOI:** 10.1101/2025.04.26.650733

**Authors:** Yue Wang, Oliver Cope, Jingting Chen, Aarav Mehta, Dalia Fleifel, Christina G. Ford, Poorya Behnamie, Molly Murray, Santiago Haase, Saygin Gulec, Logan Slade, Tim Elston, Philip M. Spanheimer, Caroline A Tomblin, Alison M Rojas, Tia Tate, Jeremy E. Purvis, Jeremy Wang, Joseph M Dahl, Samuel C. Wolff, Jeanette Gowen Cook, Elizabeth Brunk

**Affiliations:** Department of Genetics, University of North Carolina at Chapel Hill, Chapel Hill, NC 27516; Department of Biochemistry and Biophysics, University of North Carolina at Chapel Hill, Chapel Hill, NC 27516; Department of Chemistry, University of North Carolina at Chapel Hill, Chapel Hill, NC 27516; Integrative Program for Biological and Genome Sciences (IBGS), University of North Carolina at Chapel Hill, Chapel Hill, NC 27516; Department of Pharmacology, University of North Carolina at Chapel Hill, Chapel Hill, NC 27516; Computational Medicine Program, University of North Carolina at Chapel Hill, Chapel Hill, NC 27516; Lineberger Comprehensive Cancer Center, University of North Carolina at Chapel Hill, Chapel Hill, NC 27516; Department of Surgery, University of North Carolina at Chapel Hill, Chapel Hill, NC 27516; Bioskyrb Genomics, Inc. 2810 Meridian Pkwy Suite 110, Durham, NC 27713

**Keywords:** Extrachromosomal DNA, ecDNA, single cell, single-cell multi-omics sequencing, population heterogeneity, evolution

## Abstract

During tumor progression, it has been assumed that individual cells that have acquired advantageous mutations overtake the population. Cancers driven by extrachromosomal DNA (ecDNA) do not follow this paradigm. Instead, these tumors have a spectrum of oncogene copy numbers across cells, and graded ecDNA variation may function as a form of bet-hedging that equips tumors with a broad range of phenotypes. Using imaging, single-cell multiomics, and multiplexed proteomics, we systematically characterized ecDNA levels across thousands of single cells. Higher ecDNA dosage produces proportional changes in transcript abundance, chromatin accessibility, protein levels, cell-cycle progression, and proliferation. Genes amplified on ecDNA exhibit distinct transcriptional scaling regimes that shift when the same genes are reintegrated into chromosomal homogeneous staining regions. When we experimentally disrupted the continuum of ecDNA dosage by sorting cells into low- and high-copy number states, the population rapidly recovered its original, continuous distribution. Our time-course data, live-cell imaging, and stochastic models collectively show that restoring this spectrum is an active, deterministic process rather than the passive outcome of random segregation. Together, these findings position ecDNA-mediated expression as a distinct evolutionary mechanism that endows tumors with rapid, population-level adaptability. These findings offer insight into why ecDNA-driven cancers are among the most aggressive and treatment-resistant.

## Background

Gene amplification is common in cancer, and it arises through fundamentally different architectural modes (***Fig. 1a***). Some tumors amplify oncogenes within chromosomal homogeneous staining regions (HSRs)^1^. Others amplify oncogenes on extrachromosomal DNA (ecDNA), small circular fragments that replicate and segregate independently of chromosomes^2,3^. These architectures are believed to contribute to tumor heterogeneity^4–8^ and reshape tumor behavior^9–14^. A key distinction between HSR- and ecDNA-bearing cells lies in how amplified oncogene copies are distributed across the cell population (***Fig. 1b***). HSRs follow Mendelian inheritance: equal chromosomal segregation produces a limited number of discrete copy-number states. In contrast, ecDNAs segregate unevenly, dispersing amplified oncogene copies across a broad and continuous range^15–17^. At the extremes, cells within the same population can differ by more than 1000 oncogene copies^18,19^. Thus, HSR and ecDNA tumors may harbor comparable population average copy numbers, yet the architecture of amplification generates fundamentally different population structures. What selective advantage does distributed amplification confer?

**Figure 1:**
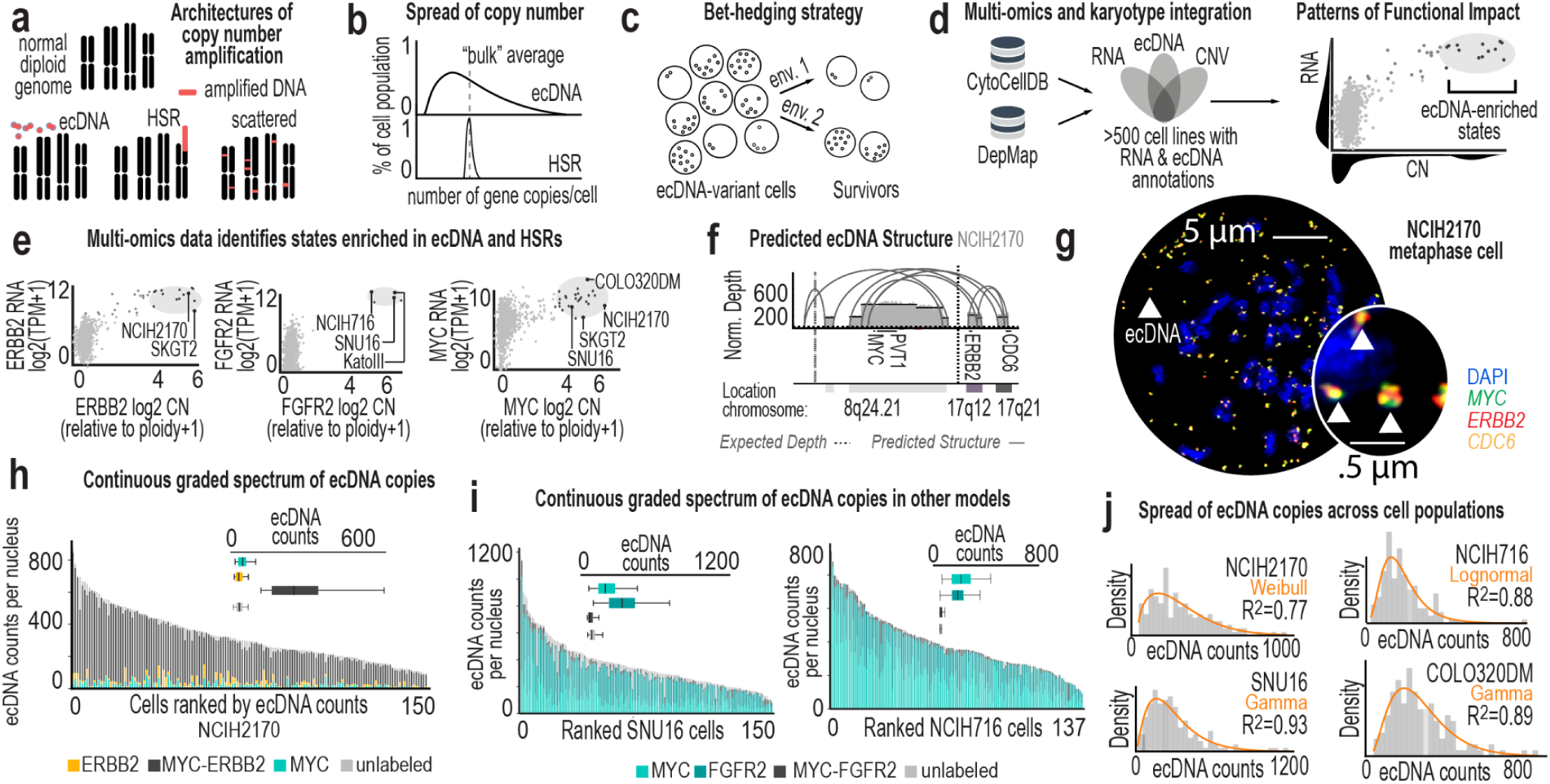
Functional genomics of ecDNA. **a**, Different architectures of gene amplification, including ecDNA, HSRs, and scattered. **b**, Schematic of number of gene copies per cell amplified on ecDNA (top) versus HSRs (bottom) across a population of cells. **c**, Schematic of exploitation of ecDNA as a “bet-hedging” strategy to increase the odds of population survival in different stress conditions or environments (env. 1 vs env. 2). **d**, Data on cell lines are available in databases such as Dependency Map and Cancer Cell Line Encyclopedia. CytoCellDB provides karyotype information on many of these cell lines. **e**, Examples of copy number vs. RNA scatter plots for three key oncogenes (*ERBB2*, left, *FGFR2*, center, *MYC*, right) In indicated cell lines. **f**, Amplicon structure of ecDNA in NCI-H2170 cells predicted using AmpliconArchitect. **g**, Metaphase FISH image of NCI-H2170 cells with triple labeling of MYC, ERBB2, and CDC6. Fluorescence channels were acquired separately and merged to generate the composite image. Probe details are provided in **Supplementary Table 3**. **h**, Continuous, skewed distribution of ecDNA counts across 150 metaphase NCI-H2170 cells. Inset shows population composition of four distinct ecDNA genetic species in NCI-H2170 cells. The dominant ecDNA species co-amplifies *ERBB2* and *MYC*. **i**, Continuous, skewed distribution of ecDNA counts across metaphase SNU16 (left) and NCI-H716 (right) cell lines. Insets indicate the compositions of ecDNA species for each cell line. **j**, Density plots of ecDNA counts (dosage) per cell for NCI-H2170 (upper left, N=150), NCI-H716 (upper right, N=150), SNU16 (lower left, N=137), and COLO320DM (lower right, N=120) cell lines.

Distributing oncogene dosage across cells may function as a bet-hedging^20^ mechanism that improves survival in fluctuating or stressful environments (***Fig. 1c***). This possibility provides an alternative to the prevailing model of cancer evolution as clonal selection^21^, where rare advantageous mutations rise to dominance resulting in one or a few major cell states^22^. ecDNA-bearing cells do not resolve into discrete clonal peaks. Instead, they maintain a persistent continuum of oncogene dosage across cells. We propose that this pattern reflects a distinct evolutionary mode, which we refer to as distributive evolution. In this mode, multiple heritable states are sustained simultaneously, creating a standing reservoir of population-level phenotypic diversity. Yet, for these heritable states to have selective value, the observed graded differences in copy number must produce graded functional phenotypes across cells in the population. Furthermore, if this continuous heterogeneity influences population-level survival, we would expect that it should be restored after disruption. Recovery of this continuous spectrum of ecDNA dosage states would reveal how populations preserve their adaptive capacity despite tradeoffs at the single cell level.

Addressing these questions requires simultaneous measurement of ecDNA dosage and functional phenotype within the same single cell as well as perturbing ecDNA distributions without introducing unrelated stress. Existing approaches rarely quantify DNA, RNA, and protein simultaneously (although DNA/RNA^23–25^ and DNA/protein^26^ quantification methods are emerging), and there are no standard tools to directly isolate or manipulate live cells based on their ecDNA states. Here, we integrate single-cell multiomics, quantitative imaging, and fluorescence-assisted cell sorting (FACS) to overcome these barriers. We map ecDNA dosage to genome-wide transcriptional and proteomic landscapes within individual cells and generate single cell whole-genome and transcriptome multiomics sequencing profiles across six models (three ecDNA- and three HSR-bearing cell lines) to directly compare regulatory consequence of genomic architecture. We isolate cells from the extremes of the ecDNA continuum and culture them independently, creating a controlled perturbation of copy-number distributions. Our results indicate that graded ecDNA dosage establishes a continuously distributed, persistent set of heritable states that enables rapid adaptation in dynamic environments. In this framework, ecDNA supports distributed evolution, a mode fundamentally distinct from classical clonal selection and potentially central to the aggressiveness and therapeutic resistance of ecDNA-driven cancers.

### Model systems for correlation of phenotypes with ecDNA copy number

Studying genotype-phenotype relationships at single-cell resolution remains challenging, particularly in ecDNA-bearing cells. Cell lines^5,27–29^, drug-selection models^30,31^, patient-derived cultures^27,32,33^ and mouse models^34^ have all been used to study ecDNA biology. However, many existing systems present challenges for functional studies because complex amplification architectures can confound interpretation (e.g. single cells can harbor both ecDNA and HSRs or multiple distinct ecDNA species)^35,36^. To identify tractable model systems, we developed CytoCellDB^19^, a curated karyotype resource of 500+ cancer cell lines within the Cancer Dependency Map (DepMap)^37^. Integrating CytoCellDB with DepMap multiomics data enabled systematic comparison of bulk copy number variation, RNA expression, and gene essentiality patterns across models (***Fig. 1d-e***).

Among these, NCI-H2170 emerged as uniquely suited for functional and evolutionary analysis. Four key growth-promoting genes (*ERBB2*, *MYC*, *PVT1*, and *CDC6*) are co-localized on a single ecDNA amplicon (***Fig. 1f-g*, *Extended Data Fig. 1a***), and none is amplified on HSRs. ecDNA copy number spans a broad and continuous range and is typically right-skewed, with most cells harboring ∼300 copies and a minority exceeding 1,000 (***Fig. 1h***). A single dominant ecDNA species accounts for the majority of amplification (***Fig. 1h* *inset***), minimizing structural complexity. This combination of ecDNA-exclusive amplification, wide dosage variability, and architectural simplicity makes NCI-H2170 a powerful system to isolate the effects of ecDNA.

We therefore selected NCI-H2170 as our primary model. Other well-studied lines, including SNU16, COLO320DM, and NCI-H716, also display broad, continuous, and skewed ecDNA dosage distributions (***Fig. 1i-j***), but harbor two dominant ecDNA amplicons, substantially increasing functional complexity despite coordinated inheritance^38^ (***Extended Data Fig. 1b***). We next defined the ultrastructure and sequence of ecDNA in NCI-H2170 using scanning electron microscopy and Correlative Light Electron Microscopy^4^ and Oxford Nanopore adaptive sampling long-read sequencing^39^ with *de novo* assembly (***Fig. 2a-b*, *Extended Data Fig. 1c-d, Supplementary Table 5***). The dominant ecDNA multimerizes fragments from 8q24, 17q12, and 17q21.2, spanning approximately three megabases. Each ecDNA carries *MYC* and *PVT1* at a 2:1 ratio relative to *ERBB2* and *CDC6* (***Fig. 2c-d***). Cells at the extremes of the dosage continuum differ strikingly in total DNA burden: ecDNA constitutes up to 30% of total DNA in ecDNA^high^ and ∼4% in ecDNA^low^ cells (***Fig. 2e***). Thus, cells within the same population differ not only in oncogene dosage but also in the amount of DNA they must replicate and regulate, imposing distinct demands on replication and chromatin organization that may underlie graded molecular and cellular states.

**Figure 2:**
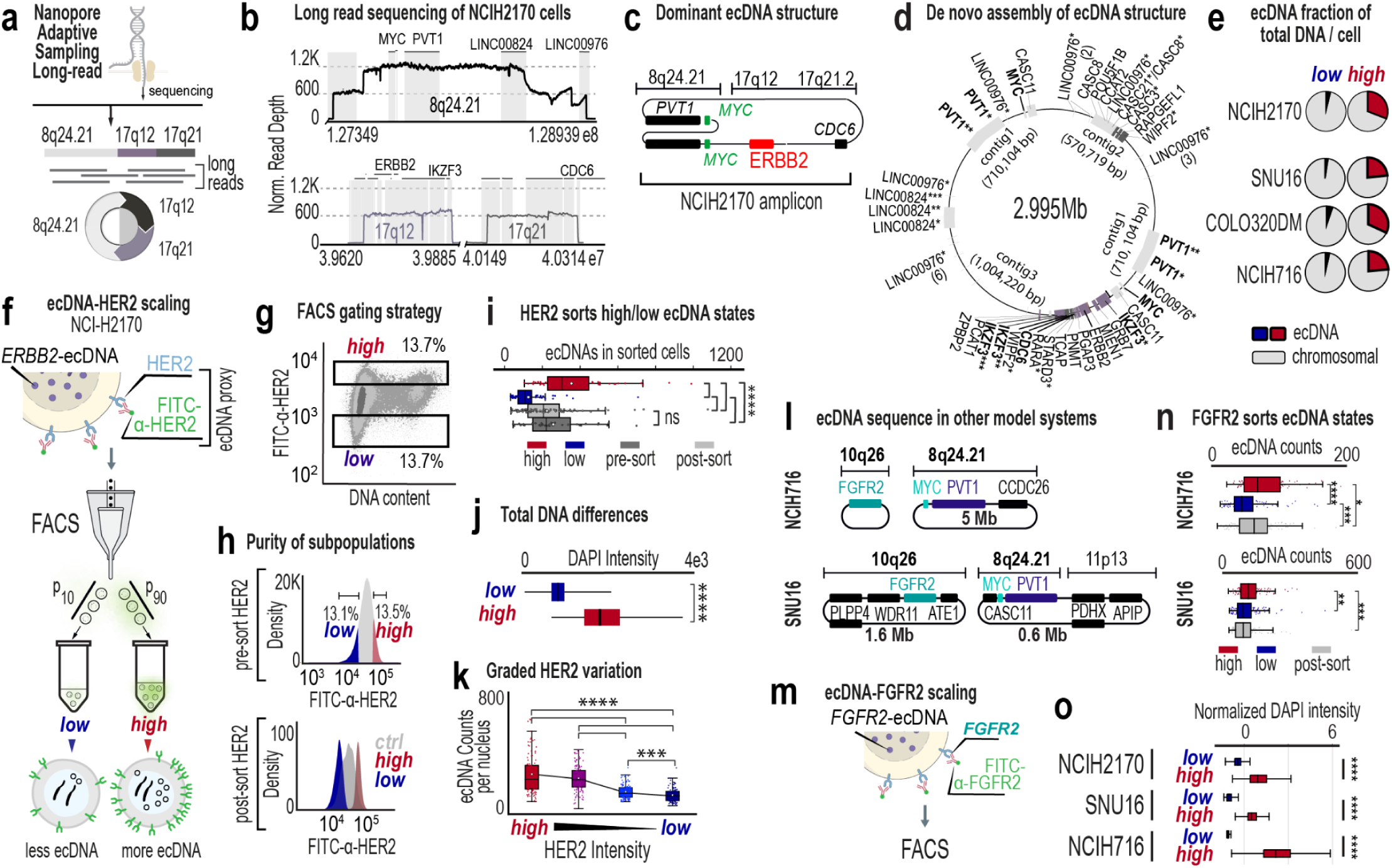
Selecting subpopulations of live cells with defined ecDNA states. **a**, Schematic of Oxford Nanopore adaptive sampling, long-read sequencing for sequencing of NCI-H2170 ecDNA chromosome 8 and 17 fragments. **b**, Long-read sequencing coverage of chromosome 8 (top) and chromosome 17 (bottom), indicating genes within these genomic windows that are highly amplified (N= ∼3 million cells). **c**, Schematic of dominant ecDNA amplicon in NCI-H2170 cells, based on long-read sequencing results. **d**, *De novo* structural assembly of ecDNA amplicon structure from long-read sequencing data.pen Asterisks denote gene fragments. **e**, Estimated total genome composition differences between ecDNA^high^ and ecDNA^low^ cells across models (e.g., fraction of ecDNA in total DNA, Supplementary Table 4). **f**, Schema of FACS approach for isolation of cells with extreme differences in HER2 fluorescence intensity and quantification of differences in ecDNA within these subpopulations. **g**, FACS gating strategy to isolate the top 13.7% and bottom 13.7% of cells (N= cells, 3 biological replicates) based on HER2 fluorescence intensity. **h**, Flow cytometry confirmation of HER2 subpopulation purity. Comparison of pre-sort (top) and post-sort (bottom) cell populations. **i**, Counted ecDNAs in sorted metaphase NCI-H2170 cells (N= 442 cells, 3 biological replicates) using DNA metaphase FISH. Comparisons between ecDNA^high^, ecDNA^low^ and control subpopulations show significant differences in ecDNA counts. **j**, Total DNA in ecDNA^high^ and ecDNA^low^ NCI-H2170 cells (N = 820 and 920 cells, two biological replicates) as quantified based on DAPI intensity across two biological replicates. **k**, ecDNA levels versus HER2 intensity in four populations of live NCI-H2170 cells (N = 453 cells). **l**, Schematics of two dominant ecDNA structures in NCI-H716 cells (top) and SNU16 (bottom) cells. **m**, Schematic of FACS approach to quantify differences in ecDNA within isolated subpopulations of cells with extreme differences in FGFR2 fluorescence intensity. **n**, Counted ecDNAs in sorted metaphase NCI-H716 cells (left, N= 286 cells) and SNU16 cells (right, N= 359 cells) using DNA FISH. **o**, Total DNA differences, measured by DAPI fluorescence intensity between ecDNA^high^ and ecDNA^low^ across model systems (two replicates, N= 364, 193 cells and N = 1,088, 749 cells for SNU16 and NCIH716, respectively).

### Isolation of live cells with defined ecDNA states

Although ecDNA dosage can be quantified via fluorescence in fixed cells, linking dosage to functional phenotypes in living cells has remained a major challenge. A direct strategy would be to isolate cells with similar loads from extreme ends of the dosage distribution, generating isogenic subpopulations that differ primarily in ecDNA dosage. However, live sorting by ecDNA level has been challenging, preventing real-time analysis of defined ecDNA states.

We exploited a key feature of NCI-H2170 cells: *ERBB2* is amplified exclusively on ecDNA. Its protein product, HER2, is an extracellular membrane receptor, enabling live-cell labeling. We used fluorophore-conjugated anti-HER2 antibodies to isolate cells expressing different HER2 levels, preserving viability and avoiding genetic engineering or permeabilization (***Fig. 2f***). Viability-gated flow cytometry cleanly separated HER2^high^ (top ∼10%), HER2^low^ (bottom ∼10%) and unsorted populations (***Fig. 2g-h***, ***Extended Data Fig. 1e***). DNA metaphase FISH confirmed that HER2^high^ and HER2^low^ subpopulations differed significantly in ecDNA dosage (***Fig. 2i*, *Extended Data Fig. 1f***), validating that HER2 abundance serves as a proxy for ecDNA dosage and total DNA content (***Fig. 2j*, *Extended Data Fig. 1g***). Sorting into four graded HER2 bins further demonstrated proportional scaling between surface HER2 and ecDNA dosage (***Fig. 2k***).

We applied the same sorting strategy to sorting live cells from SNU-16 and NCI-H716, where FGFR2 is an extracellular protein encoded on ecDNA (***Fig. 2l-m***). In both models, FGFR2 protein levels tracked with FGFR2-amplified ecDNA dosage (***Fig. 2n***) and total DNA (***Fig. 2o***). These results, together with previous work^40^, establish ecDNA-encoded protein-based sorting as a direct and scalable method to isolate live ecDNA dosage-defined subpopulations, enabling functional analyses that were previously inaccessible.

### ecDNA dosage restructures cell cycle networks

Applying our isolation strategy, we examined how ecDNA dosage shapes cell cycle progression and proliferation-associated regulatory programs (***Fig. 3a***). First, EdU-based analytical flow cytometry cell cycle experiments^41^ revealed that ecDNA^high^ cells were enriched in S and G2/M phases, whereas the ecDNA^low^ cells were enriched in G1 across three models (***Fig. 3b–d***). Second, we quantified transcript levels of 24 proliferation-regulatory genes in NCI-H2170 ecDNA^high^ and low-ecDNA^low^ cells. Subpopulations displayed significant transcriptional differences that extended beyond ecDNA-encoded loci (e.g., *ERBB2*, *MYC*, and *CDC6*), to genes distributed across the genome, indicating that ecDNA dosage influences a broader regulatory program (***Fig. 3e***). Third, we used indirect iterative immunofluorescence imaging (4i^42^) to simultaneously measure 29 regulatory proteins across thousands of single cells (***Extended Data Fig. 1h***). Using HER2 in NCI-H2170 and FGFR2 in SNU-16 and NCI-H716 as ecDNA proxies, we directly quantified protein–dosage relationships within individual cells.

**Figure 3:**
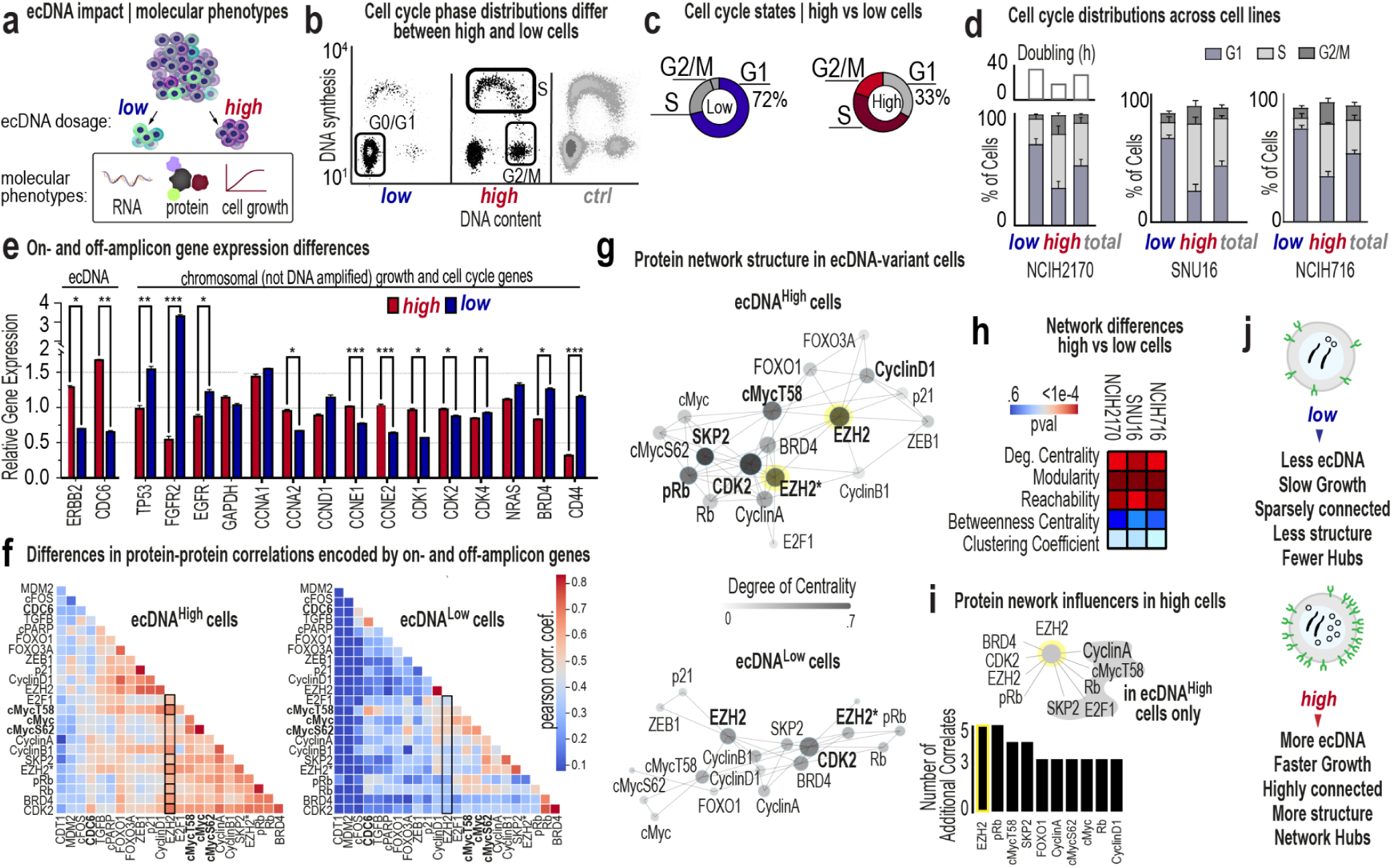
ecDNA dosage organizes a global regulome. **a**, Schema for sorting live cells into “high” and “low” ecDNA subpopulations. **b**, Illustration of EdU-based cell cycle analyses and partitioning of NCI-H2170 cells into ecDNA^high^ and ecDNA^low^ variant populations to compare their cell cycle distributions (as in Fig 2g). Representative of three independent biological replicates. **c**, Diagrams of percentages of ecDNA^high^ and ecDNA^low^ populations from NCI-H2170 cells in G1, S, and G2/M cell-cycle phases. **d**, Percentages of ecDNA^high^ and ecDNA^low^ populations from indicated cell lines in G1, S, and G2/M cell-cycle phases. Each graph is pooled from three independent biological replicates in each cell line; mean with error bars ± SEM. Doubling estimated by real time growth analysis. **e**, RNA quantification of 24 genes via qPCR in ecDNA^high^ and ecDNA^low^ populations from NCI-H2170 cells (N = 3 technical replicates). Genes encoded by the ecDNA and by chromosomes are indicated. **f**, Inter-protein correlations across 29 proteins from iterative indirect immunofluorescence imaging data in NCI-H2170 cells across two independent biological replicates (wells). **g**, Differences in protein network interconnectivity (based on protein abundance correlations) between ecDNA^high^ and ecDNA^low^ NCI-H2170 cells. **h**, Network analysis of ecDNA^high^ versus ecDNA^low^ NCI-H2170, SNU16, and NCI-H716 cells reveals significant differences in connectivity, organization, and degree centrality. **i**, Number of correlates between prominent “influencer” proteins in ecDNA^high^ NCI-H2170 cells. **j**, Schematic of how ecDNA^high^ and ecDNA^low^ cells establish growth differences through restructuring of transcript and proteomic networks.

Network analysis of 4i data revealed that ecDNA^high^ cells form significantly denser and more coordinated regulatory networks than ecDNA^low^ cells, which exhibit fragmented and weakly correlated interactions (***Fig. 3f-g***). In NCI-H2170 ecDNA^high^ cells, strong connectivity emerged among transcriptional and chromatin regulators, including both on-amplicon proteins (*HER2*, *CDC6*, *MYC*) and chromosomally-encoded factors (*EZH2*, *Cyclin D1*, *ZEB1*, *cFOS*, *E2F1*, *Rb*), consistent with integrated control of proliferation and transcription^43^. Notably, amplified cell-cycle genes displayed distinct scaling behaviors. MYC protein increased with ecDNA dosage but showed weaker scaling than other ecDNA-associated genes, indicating that MYC abundance is less tightly coupled to ecDNA level than genes such as the ecDNA-encoded CDC6 and the chromosomally encoded Cyclin D1, a regulator of G1 progression (***Extended Data Fig. 1i-j***). However, phosphorylated MYC species occupied more central positions in ecDNA^hig^ protein networks, suggesting that MYC-related regulatory state changes may be more evident in network organization than in proportional changes in total MYC abundance. Consistent with these observations, analysis of residualized protein program scores derived from the 4i panel revealed that ecDNA^high^ cells exhibit significantly elevated replication/proliferation signaling, whereas ecDNA^low^ cells are relatively enriched for checkpoint and stress-associated proteins, producing a strongly separated net cell-cycle activity score (Mann-Whitney *P* ≈ 4 × 10⁻¹¹¹, ***Extended Data Fig. 1k***).

Network centrality, reflecting the influence of a protein within the regulatory system^44,45^, differed significantly between ecDNA^high^ and ecDNA^low^ cells across models (***Fig. 3h***). In NCI-H2170 ecDNA^high^ cells, EZH2 showed markedly increased network centrality, with abundance correlated to BRD4, phosphorylated MYC, E2F1, and multiple proliferation-associated regulators (***Fig. 3i***). In contrast, EZH2 displayed minimal connectivity in ecDNA^low^ cells. Similar shifts in network connectivity were observed in SNU-16 and NCI-H716 (***Extended Data Fig. 1l***), indicating that ecDNA dosage is associated with large-scale reorganization of regulatory networks. These network changes coincide with the elevated proliferation-associated signaling programs observed in ecDNA^high^cells (***Fig. 3j***).

### A continuous ecDNA dosage axis structures the population for distributed evolution

Cells at the extreme ends of the ecDNA spectrum have been shown to occupy distinct phenotypic states^26,40^, and display dosage-dependent transcriptional differences across cells^10,32,33^, consistent with our data. Whether intermediate states form a structured molecular continuum across the population has remained unresolved.

To address this, we performed ResolveOME^25^ to simultaneously sequence whole genomes and transcriptomes from hundreds of single cells across six model systems, including three ecDNA- and three HSR-bearing models (***Fig. 4a***). For each single cell, we measured an ecDNA-encoded protein prior to sequencing, generating matched DNA, RNA, and protein data within the same single cells. Inferred ecDNA copy number closely matched DNA metaphase FISH distributions across models (***Fig. 4b*, *Extended Data Fig. 2a-k***). HER2 in NCI-H2170 and FGFR2 in SNU-16 scaled proportionally with ecDNA dosage (***Fig. 4c*, *Extended Data Fig. 2h***), whereas MYC protein in COLO320DM did not (***Extended Data Fig. 2k***), indicating gene-specific buffering.

**Figure 4:**
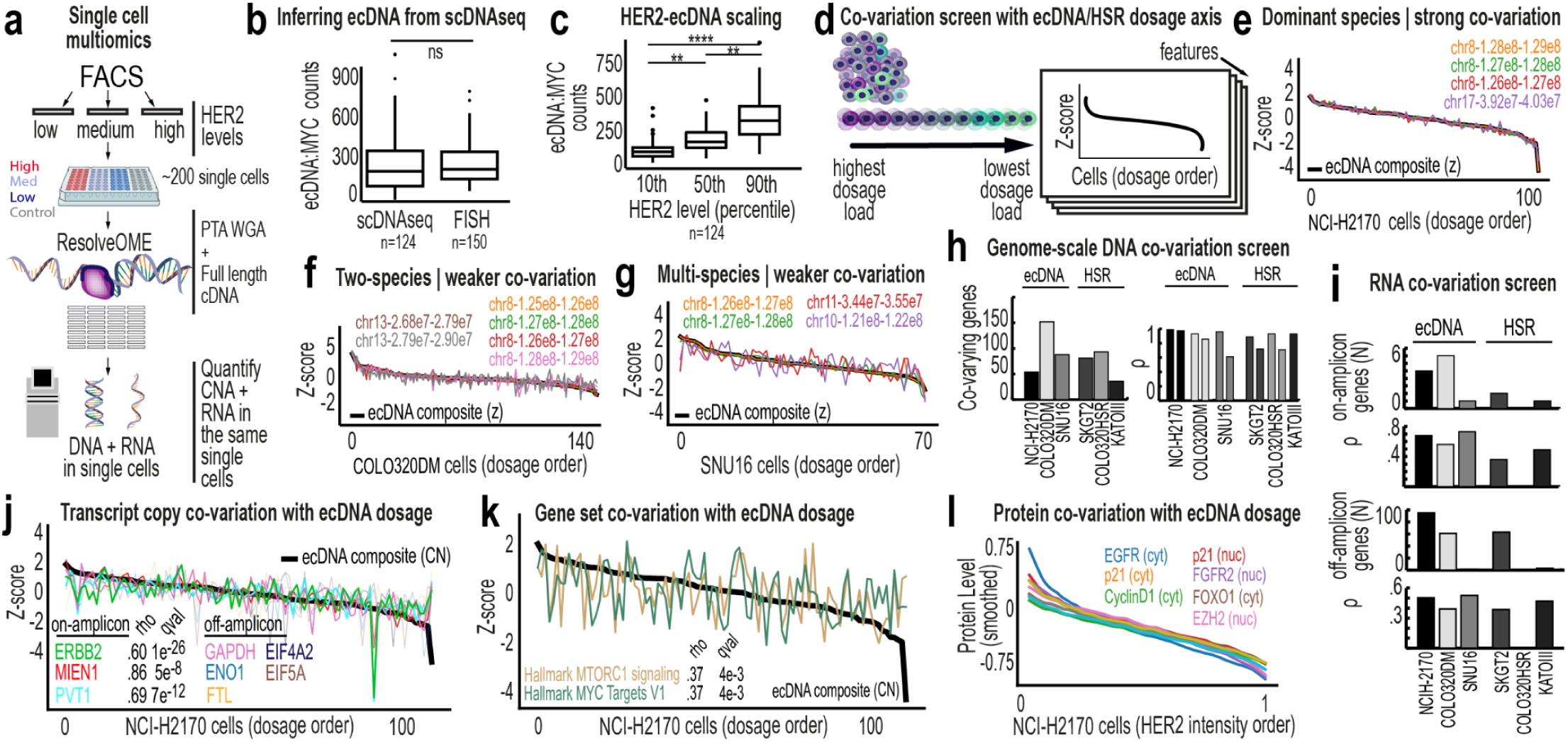
A continuous ecDNA dosage axis structures the population for distributed evolution. **a**, Overview of single-cell multiomics sequencing using the ResolveOME platform to simultaneously quantify DNA and RNA. **b**, Inferred ecDNA copy numbers obtained from single cell DNA sequencing (N=124 cells, two independent biological replicates) and quantified ecDNA counts obtained from DNA metaphase FISH experiments in NCI-H2170 cells (N=150 cells). **c**, Inferred ecDNA copy numbers in NCI-H2170 cells (N=124 cells, two independent biological replicates) sorted by HER2 protein levels (percentile, measured by fluorescence intensity via FACS). **d**, Schematic of cells ordered along a continuous ecDNA/HSR dosage axis defined by the composite copy number trajectory (black line), which represents the inferred ordering of single cells by amplification level. Each genomic bin, transcript, and protein was then evaluated individually to determine whether its abundance changed monotonically along this ordering using Spearman correlation. Values are shown as z-scores relative to the population mean. **e**, Co-variation plot showing specific genomic bins that significantly co-vary with ecDNA dosage axis in NCI-H2170 cells (N=139 cells, two independent biological replicates). **f**, Co-variation plot showing specific genomic bins that significantly co-vary with ecDNA dosage axis in COLO320DM cells (N= 175 cells, two independent biological replicates). **g**, Co-variation plot showing specific genomic bins that significantly co-vary with ecDNA dosage axis in SNU16 cells (N= 111 cells, two independent biological replicates). **h**, Comparison between ecDNA and HSR models in terms of the number of genes that significantly co-vary with ecDNA/HSR dosage trajectory (left) and the average ρ values (right) that corresponds to these co-varying features. **i**, Number of co-varying genes and their ρ values (correlation strength with dosage trajectory) for on-amplicon (top) and off-amplicon (bottom) genes. **j**, Co-variation plot showing specific RNA transcripts that significantly co-vary with ecDNA dosage axis in 97 NCI-H2170 cells. **k**, Co-variation plot of specific transcriptional hallmark gene sets that significantly co-vary with the ecDNA dosage axis in 97 NCI-H2170 cells. **l**, Co-variation plot showing specific protein levels (smoothed) that significantly co-vary with the ecDNA dosage axis in NCI-H2170 cells.

We next asked whether amplification architecture defines a coordinated genomic axis. Using an unbiased computational approach, we identified genomic bins whose copy number co-varied smoothly along the ecDNA or HSR dosage axis (***Fig. 4d***) and identified fragments with verified amplified genes (***Fig. 4e-g***). In NCI-H2170, fragments from chromosomes 8 and 17 co-varied with near-perfect synchrony (Spearman ρ ≈ 0.9), precisely reconstructing the ecDNA amplicon mapped by long-read sequencing (***Fig. 4e***). Cell lines harboring multiple ecDNA species showed weaker coordination (***Fig. 4f-g***), while HSR models exhibited limited co-variation (***Fig. 4h***, ***Extended Data Fig. 3a-b***).

We extended this framework to RNA and proteins. In ecDNA models, 50-100 transcripts co-varied along the dosage axis, far beyond those physically encoded on ecDNA (***Fig. 4i*, *Extended Data Fig. 3c***). HSR models showed substantially fewer dosage-associated transcripts (***Extended Data Fig. 3d***). In NCI-H2170, nearly 100 genes scaled smoothly with ecDNA dosage, forming a graded transcriptional program rather than discrete expression clusters (***Fig. 4j***). MYC target genes and MTORC1 signaling were among the most dosage-responsive pathways (***Fig. 4k***). Proteomic profiling revealed similar scaling. Numerous proteins not encoded on ecDNA, including EGFR, Cyclin D1, p21, FGFR2, FOXO1, and EZH2, increased proportionally with ecDNA dosage in NCI-H2170 cells (***Fig. 4l***). HER2 protein levels increased with ERBB2, GRB7, and CDC6 transcript abundance across cells. (***Extended Data Fig. 3e-f***), whereas MYC transcripts scaled more weakly (***Extended Data Fig. 3g***). Representative RNA-DISH images further illustrate cell-to-cell variability in ERBB2 transcript signal (***Extended Data Fig. 3h***).

To assess epigenetic coupling, we inferred ecDNA dosage in 10X multiOME data using an XGBoost model trained on ResolveOME RNA features (PCC > 0.9; ***Extended Data Fig. 4a–c***), because dosage distributions inferred from scWGS are more consistent with ground-truth FISH data than those inferred from scATAC (***Extended Data Fig. 4d–e***). In NCI-H2170 cells, ecDNA dosage was associated with coordinated shifts in metabolic, stress-response, and ribosomal programs (***Extended Data Fig. 4f–h***). Chromatin accessibility changes were concentrated at regulatory regions linked to ecDNA genes. Peaks on chromosome 17 correlated with multiple ecDNA-encoded transcripts, while long-range interactions on chromosome 8 modulated PVT1 and CCDC26 expression (***Extended Data Fig. 5a–b***). The most dosage-sensitive peaks are localized within IKZF3, which is fragmented and repositioned on ecDNA (***Extended Data Fig. 5c-e***). Regions from chromosomes 17 and 8 formed distinct regulatory modules (***Extended Data Fig. 5b,f–h***) with IKZF3 fragments showing the strongest correlations to chromosome 17 gene expression. These patterns are consistent with ecDNA-driven reorganization of cis-regulatory architecture (***Extended Data Fig. 5i***).

To assess whether mutational differences contribute to the observed phenotypic variation, we performed pseudobulk variant calling^46^ using ResolveOME single-cell DNA data (***Extended Data Fig. 6a-b***). NCI-H2170 ecDNA^high^ and ecDNA^low^ cells showed no significant differences in variant burden within amplified regions (***Extended Data.* *Fig. 6c-f***), and shared the same activating *ERBB2* mutation^47,48^ (***Extended Data Fig. 6g-h***). Similar analyses in COLO320DM and SNU16 cells yielded consistent results, with no detectable differences in variant burden between ecDNA^high^ and ecDNA^low^ populations (***Extended Data Fig. 6i-j***). These findings indicate that phenotypic divergence arises from variation in ecDNA dosage rather than clonal mutation. Consistent with this, ecDNA dosage defines a continuous organizing axis that propagates across genomic, transcriptional, and protein-level regulation.

### Amplification architecture influences transcriptional scaling

Previous studies have shown that ecDNA can shape chromatin accessibility and regulatory landscapes^9,49,50^. We asked whether amplification architecture also alters transcriptional yield per gene copy. Across six models, we related transcript abundance to ecDNA dosage within the same single cells and classified scaling behavior as linear, threshold, or saturating (***Fig. 5a***).

**Figure 5:**
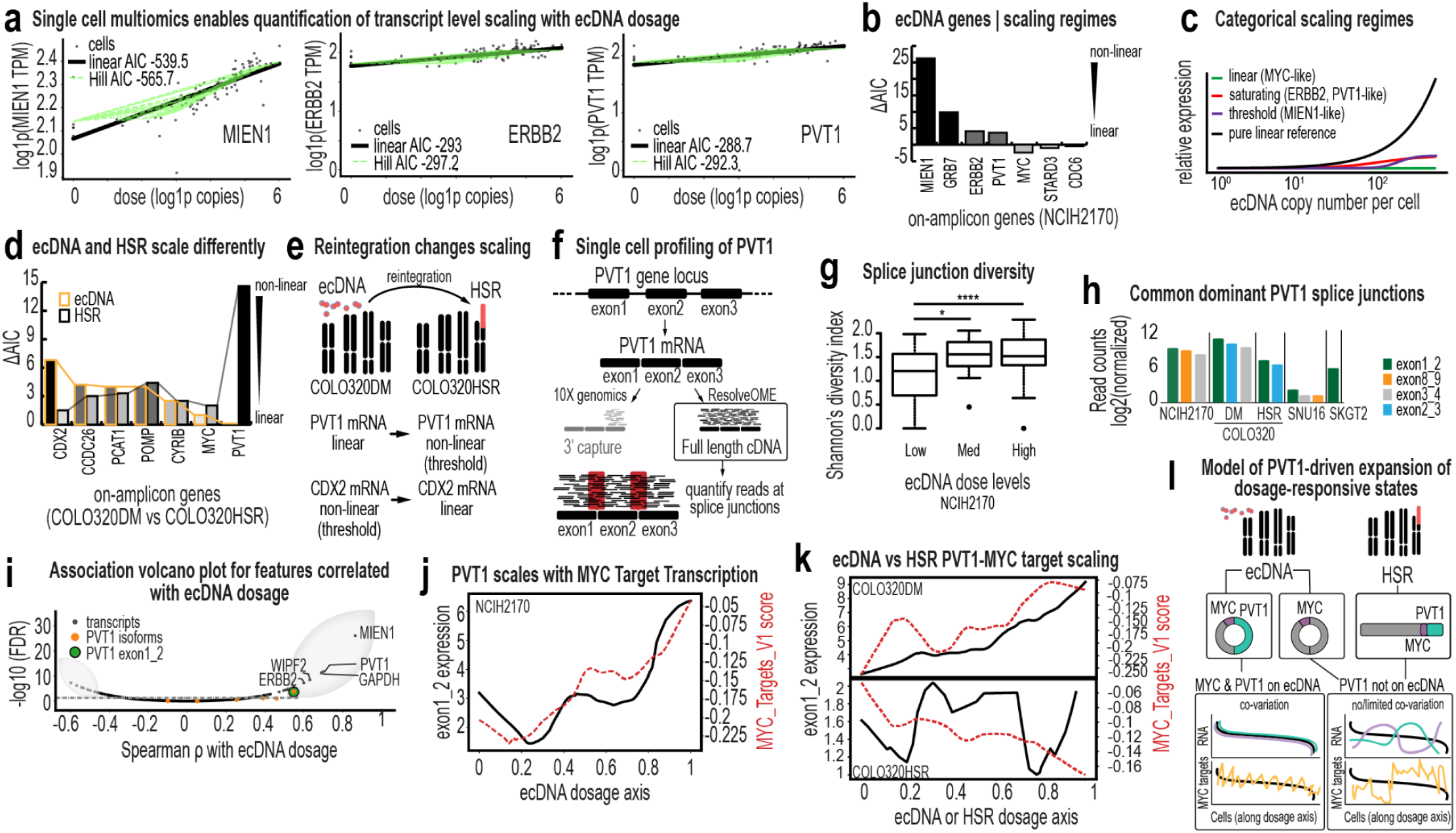
Amplification architecture influences transcription. **a**, Scatter plots showing transcriptional yield (TPM) per ecDNA dosage. Two models (linear and Hill) are shown across three different ecDNA-amplified genes in NCI-H2170 cells. AIC (Delta Akaike Information Criterion) is computed for each model, where lower AIC values indicate a better model fit. **b**, Bar plot showing ΔAIC values (measuring the relative difference between linear and Hill models per gene) for on-amplicon genes in NCI-H2170 cells. ΔAIC > 2 suggests substantial differences between models and ΔAIC > 6 indicates strong evidence of non-linear transcriptional scaling (favoring the Hill model). **c**, Cartoon schematic illustrating representative transcriptional scaling regimes identified in NCI-H2170 cells. Curves represent model classes (e.g., linear, threshold, and saturating) used to fit gene expression as a function of ecDNA dosage. Individual genes were assigned to the model providing the best fit based on ΔAIC. **d**, Bar plot showing ΔAIC values for amplified genes in COLO320DM (orange border) and COLO320HSR (black border). The scaling regimes of *PVT1* and *CDX2* are most influenced by amplification architecture. **e**, Schematic showing the change in scaling regimes of *PVT1* and *CDX2* change when ecDNA reintegrates into HSR. **f**, Schematic illustrating the quantification of splice junction reads using ResolveOME to infer *PVT1* mRNA isoform landscape in single cells. **g**, Box plots showing splice junction diversity in NCI-H2170 cells across low, medium and high ecDNA dosage groups (N=215 cells). **h**, Normalized read counts across *PVT1* splice junctions across model cell lines. **i**, Volcano-style association plot showing the strength of association (Spearman correlation, ρ, with the ecDNA dosage axis) versus statistical significance (–log10 FDR) to identify features that scale monotonically with ecDNA dosage. **j**, Continuous ecDNA dosage (x-axis) plotted against *PVT1* exon 1–2 read counts (left y-axis) and *MYC* target gene set enrichment scores (right y-axis). Both structural transcript abundance and *MYC* target gene expression increase monotonically with ecDNA dosage, demonstrating coordinated scaling of oncogene output and transcription regulatory network activity. **k**, *PVT1* exon 1–2 read counts and *MYC* target enrichment scores are plotted against the dosage axis in COLO320DM (ecDNA, top) and COLO320HSR (HSR, bottom) cells. COLO320DM cells show robust monotonic scaling of transcript abundance and downstream *MYC* target activation with increasing dosage, whereas COLO320HSR cells display reduced or altered coupling, suggesting that ecDNA amplification confers distinct regulatory dynamics. **l,** Proposed model of *PVT1*-driven expansion of dosage-responsive transcriptional states. Amplification of *PVT1* on ecDNA broadens the set of transcripts that co-vary with ecDNA dosage, increasing the fraction of genes exhibiting graded, population-wide scaling behavior.

In NCI-H2170 cells, ecDNA-encoded genes displayed heterogeneous scaling regimes (***Fig. 5b***). Some genes increased proportionally with copy number, whereas others exhibited threshold-like or saturating behavior. Thus, ecDNA does not function as a uniform high-copy template; individual genes respond differently to increased dosage, adding complexity to proposed hypertranscription models of ecDNA regulation^51^. Across all models, genes clustered into the three scaling categories (***Fig. 5c***). To determine whether scaling depends on amplification architecture, we leveraged a quasi-isogenic pair, COLO320DM and COLO320HSR, in which the same oncogenes are amplified on ecDNA or reintegrated HSRs, respectively. Strikingly, *PVT1* and *CDX2* exhibited distinct scaling regimes depending on their genomic context (***Fig. 5d–e***). On ecDNA, *PVT1* scaled linearly and *CDX2* followed a threshold-like (non-linear) scaling regime; in the HSR model, both genes displayed altered profiles.

The pronounced architectural sensitivity of *PVT1* prompted further analysis. ResolveOME enabled isoform splice junction profiling of this long noncoding RNA (***Fig. 5f*, *Extended Data Fig. 7a-e***). In NCI-H2170 cells, splice-junction diversity increased with ecDNA dosage (***Fig. 5g***) and ecDNA^high^ and ecDNA^low^ cells exhibited distinct splice-junction profiles (***Extended Data Fig. 7f***). Across ecDNA models, exon 1–2 junction reads predominated (***Fig. 5h*, *Extended Data Fig. 7e***), consistent with prior reports^52^. Both *PVT1* abundance and exon 1–2 usage scaled monotonically with ecDNA dosage (***Fig. 5i*, *Extended Data Fig. 7g-h***), a pattern not observed in HSR models (***Extended Data Fig. 7i***). PVT1 expression also tracked MYC target gene expression across two ecDNA models (NCI-H2170 and COLO320DM; ***Fig. 5j*, *Extended Data Fig. 7j***), linking architectural context to broader transcriptional programs.

In SNU16 cells, however, PVT1–dosage co-variation was attenuated (***Extended Data Fig. 8a***). ResolveOME and 10X multiOME revealed a two-state population (***Extended Data Fig. 7i***, ***Extended Data Fig. 8b-g***), and DNA FISH confirmed that 37% of cells lacked PVT1 amplification on ecDNA (***Extended Data Fig. 8h-i***). This observation is further supported by analysis of reads spanning PVT1–PDHX junctions, which were detected in only 35-48% of reads at these loci, consistent with only a subset of PVT1 copies being incorporated into ecDNA (***Extended Data Fig. 8j***). Within this subset, PVT1 did not scale with ecDNA dosage. Together, these findings indicate that transcriptional scaling and downstream co-variation depend on whether PVT1 resides on ecDNA. When amplified extrachromosomally, PVT1 reports graded copy-number states through both expression and isoform usage; when confined to HSRs or absent from ecDNA, this architecture-dependent behavior is diminished (***Fig. 5l***).

### Populations rapidly rebuild the ecDNA continuum consistent with asymmetric inheritance and division

If the continuous ecDNA spectrum contributes to population-level fitness, its disruption would be expected to result in restoration. We live-sorted NCI-H2170 cells into ecDNA^high^ and ecDNA^low^ populations and quantified ecDNA copy number by DNA metaphase FISH at 0, 48, 96, and 432 hours post-sort (***Fig. 6a*, *Extended Data Fig. 9a***). Rather than drifting toward new equilibria, both subpopulations rapidly reconstituted the same continuous dosage distribution within 3 generations (***Fig. 6b-c***). Across more than 1,650 images analyzed by AI-assisted quantification (***Extended Data Fig. 9b-i***), initial differences collapsed within 96 hours, and by nine generations sorted populations were indistinguishable from controls (***Extended Data Fig. 10a-c***). HER2 protein levels and ecDNA-encoded transcripts similarly returned to baseline (***Extended Data Fig. 10d-f***). Mixed-effects modeling confirmed a significant convergence of ecDNA copy number distributions over time (***Fig. 6d***). Longitudinal sorting and metaphase FISH in two additional ecDNA-positive models, SNU-16 and NCI-H716, demonstrated similar rapid restoration dynamics (***Extended Data Fig. 10b***).

**Figure 6:**
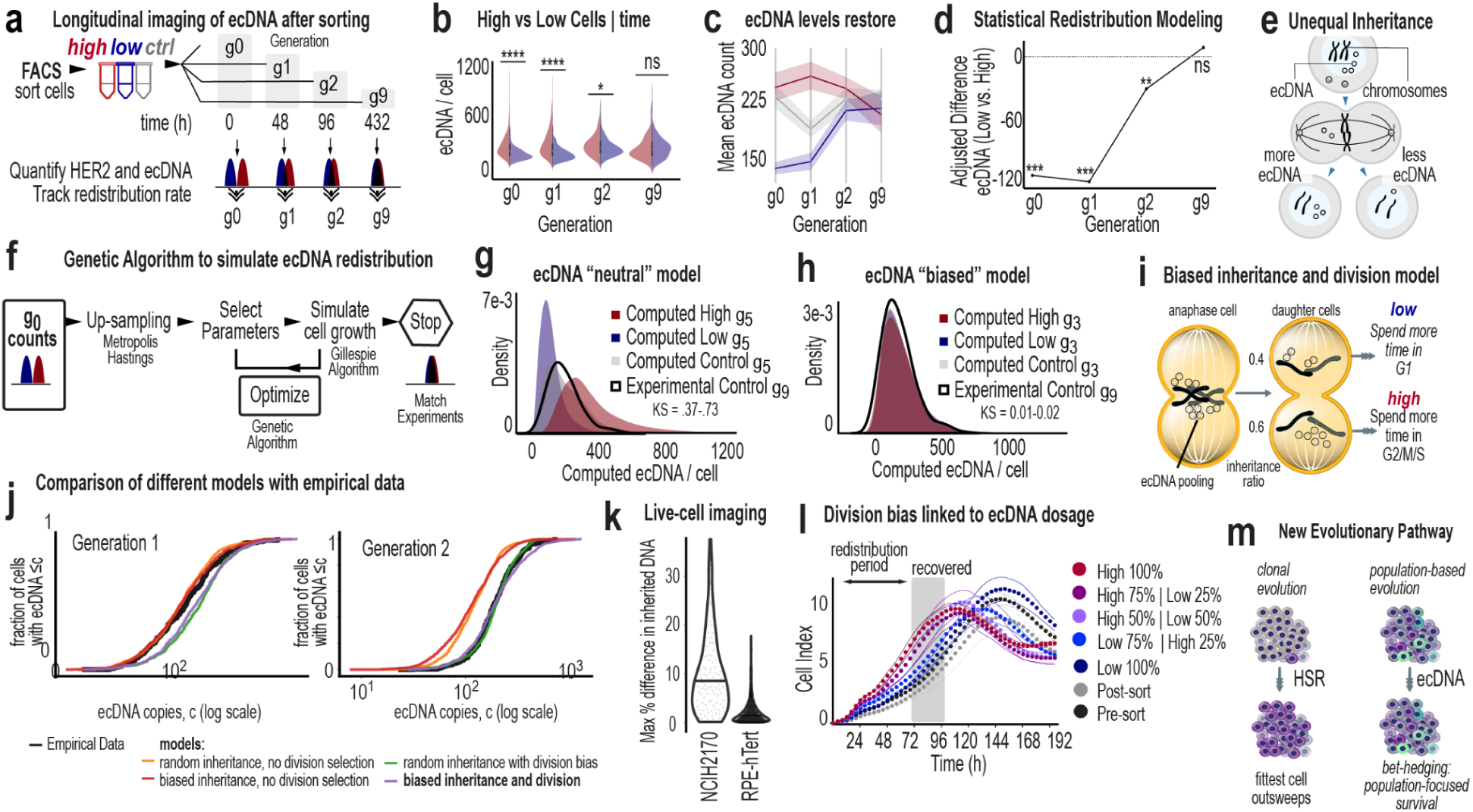
Populations rapidly restore baseline ecDNA variation levels. **a**, Schema showing the strategy to track longitudinal changes in ecDNA dosage via DNA metaphase FISH after sorting ecDNA^high^ and ecDNA^low^ cells. **b**, Split violin plot showing the changes in the distributions of ecDNA dosage over time in post-sorted ecDNA^high^ (red) and ecDNA^low^ (blue) NCI-H2170 cells (N = ∼260 cells, two biological replicates). Sorted control cells were also quantified (not shown). **c**, Changes in mean ecDNA dosage over time in ecDNA^high^ (red), ecDNA^low^ (blue) and control populations. Shaded regions indicate 95% CI. **d**, Adjusted ecDNA dosage differences between ecDNA^high^ and ecDNA^low^ populations over time, computed with mixed linear effects models. **e**, Uneven inheritance of ecDNA during mitosis. **f,** Genetic algorithm used in combination with Gillespie models to simulate ecDNA copy number variation rebound after cell sorting. **g**, Distributions of ecDNA dosage in simulated ecDNA^high^, ecDNA^low^ and controlled populations after five generations (g5) using a neutral ecDNA model, in which division and death hazards are constant across cells and do not depend on ecDNA dosage, and inheritance is unbiased binomial with p=0.5. **h**, Distributions of ecDNA dosage in simulated ecDNA^high^, ecDNA^low^ and controlled populations after three generations (g3) using a biased ecDNA model that assumes biased inheritance and division. **i**, Proposed model for biased inheritance and division and how these characteristics influence population remodeling. **j**, Computational models that assess the importance of biased inheritance and division on ecDNA variation recovery. **k,** Percent difference in inherited DNA between sister cells in an ecDNA-bearing cell line (NCI-H2170, 96 sister pairs) and a non-ecDNA bearing cell line (RPE-hTert, 123 sister pairs), representative of two independent experiments. **l,** Kinetic growth curves of sorted ecDNA^high^, ecDNA^low^, and mixtures of ecDNA^high^-ecDNA^low^ NCI-H2170 cells. Curves represent mean trajectory across 6 technical replicate wells and dashed lines represent variability (mean ± standard deviation) across three technical replicates from one independent experiment (biological n = 1). Controls considered both pre-sort and post-sort NCI-H2170 cells. **m,** Model of proposed distributive evolution pathway enabling a bet-hedging strategy.

We next asked whether random inheritance alone could explain this rapid recovery (***Fig. 6e***). We developed a stochastic model incorporating ecDNA-dependent division rate, death rate, and segregation bias (***Fig. 6f***). Parameter optimization using a genetic algorithm identified solutions that reproduced the observed dynamics (***Fig. 6g***, ***Extended Data Fig. 10g-i***). Models restricted to random segregation or ecDNA-scaled fitness alone failed to capture the speed and symmetry of restoration (***Fig. 6g***). In contrast, models consistent with both asymmetric inheritance and ecDNA-dependent division recapitulated rapid recovery of the continuum (***Fig. 6h***).

Best-fitting solutions converged on two features (***Fig. 6i***): ecDNA^high^ cells divide faster than ecDNA^low^ cells and asymmetric ecDNA inheritance, with daughter cells inheriting approximately 60% and 40% of ecDNA copies. Together, these parameters restored the continuum within three generations (***Fig. 6j*, *Extended Data Fig. 10j-k***). Imaging of post-mitotic daughter cells was qualitatively consistent with asymmetric DNA partitioning patterns consistent with the segregation bias inferred from the model. (***Fig. 6k***), and growth assays demonstrated faster proliferation of ecDNA^high^ cells. Proportional-mix experiments demonstrated that population growth scales with the fraction of ecDNA^high^ cells (***Fig. 6l***). Recovery was asymmetric, with ecDNA^low^ cells increasing their copy number nearly fourfold faster than ecDNA^high^ cells lost copies, despite their initially slower proliferation rates. (***Fig. 6l***).

## Discussion

Cancer evolution is classically framed as clonal selection, in which a limited number of advantageous genomic states expand through discrete sweeps. Our results indicate that ecDNA-driven tumors operate differently. Rather than collapsing into dominant clonal states, they maintain a broad, ordered continuum of oncogene dosage across cells. This continuum is not stochastic noise but a structured, heritable feature that generates a wide spectrum of molecular and phenotypic states. We refer to this population-level strategy as distributed evolution. In contrast to clonal systems, which rely on the emergence of rare advantageous variants, ecDNA-bearing populations are poised to respond immediately to environmental pressures by sampling diverse phenotypic states in parallel.

We show that graded ecDNA dosage produces coordinated continua of genomic, epigenomic, transcriptomic, and proteomic traits, reorganizing networks that govern proliferation, signaling, and chromatin regulation. Amplification architecture itself encodes regulatory logic: genes amplified on ecDNA follow distinct transcriptional scaling regimes that differ from those of the same genes amplified within HSRs. Thus, amplification on ecDNA is not merely an increase in copy number. It alters how gene dosage is interpreted, reshaping the quantitative relationship between genotype and phenotype.

We further demonstrate that the ecDNA dosage continuum is actively maintained. When cells at the extremes of copy number were isolated, the full distribution was rapidly restored within a few generations. This recovery cannot be explained by random segregation alone. Instead, the observed dynamics are most consistent with a model incorporating asymmetric inheritance together with ecDNA-dependent division, indicating that ecDNA-bearing populations possess intrinsic mechanisms to rebuild their evolutionary landscape. Elucidating the molecular mechanisms underlying this asymmetric inheritance will be an important direction for future work.

Together, these findings redefine ecDNA as a structured system for generating and preserving adaptive potential (***Fig. 6m***). ecDNA-driven cancers are not aggressive solely because oncogenes are amplified, but because amplification architecture confers enhanced evolvability. By maintaining a continuously distributed set of heritable states and the capacity to regenerate them, ecDNA-positive tumors achieve population-level resilience that may underlie their aggressiveness and therapeutic resistance. Targeting the architecture, dynamics, or regulatory consequences of ecDNA may therefore disrupt not only oncogene dosage but the evolutionary engine that sustains these cancers.

## Data Availability

All scripts are available at GitHub (https://github.com/Brunk-Lab). NCIH2170 single-cell multiomics data and ONT long-read sequencing data are available on Gene Expression Omnibus (GEO accession numbers GSE293273 and GSE293274). Images and tutorials are available on FigShare (10.6084/m9.figshare.28784999).

## Methods

### Primary cell culture

NCI-H2170, SNU16, COLO320DM, NCI-H716, COLO320HSR, SKGT2 and KATOIII cells were purchased from ATCC and were grown in RPMI media (Gibco) supplemented with 10% heat-inactivated fetal bovine serum (Gibco) in a humidified incubator with 5% CO_2_. Adherent cells were harvested with 0.25% trypsin in DPBS (Gibco). Viable cells were stained with trypan blue (Invitrogen, T10282) and counted using a Countess 3 (Invitrogen) counter. All cells were collected for metaphase and karyotype (G-banding) experiments within three passages. Cell doubling time for adherent NCI-H2170 cells was estimated from real-time impedance growth curves measured using the xCELLigence RTCA system, by fitting exponential growth models to the log-phase portion of the cell index trajectory.

### Metaphase sample preparation and imaging

Cells were arrested at metaphase by treating with 0.1 µg/mL colcemid (FUJIFILM, Irvine Scientific) for 12–20 hours when they reached ∼70% confluency. Colcemid inhibits spindle formation and cytokinesis. After treatment, adherent cells were detached via trypsinization, quenched with cold colcemid-spiked media and PBS, and centrifuged at 1.5g for 2 minutes. The cells were incubated with 600 µL of 0.075 M KCl (Gibco) at 37 °C for 15 minutes to induce osmotic swelling. Freshly prepared modified Carnoy’s fixative (3:1 methanol:glacial acetic acid) was added dropwise, followed by centrifugation and resuspension. This fixation was repeated three times, with the final density adjusted to ∼6 x 10^6^ cells/mL for single-cell imaging. Slides were prepared using Superfrost™ Microscope Slides (Fisherbrand, Cat. No: 12550123). A drop of the prepared cell suspension was placed on each slide and allowed to air dry for 1 hour. The slides were then equilibrated in 2x saline sodium citrate (SSC, Invitrogen, Cat. No: 15557-036) and dehydrated with an ethanol series (70%, 85%, 100%). DNA FISH probes (purchased from Empire Genomics) targeting ERBB2 (17q12), MYC (8q24), CDC6 (17q21.2) and PVT1 (8q24) were labeled with spectrally distinct fluorophores and hybridized according to standard protocols. Genomic coordinates and probe details are provided in **Supplementary Table 3**. Fluorescent DNA probes in hybridization buffer (5 µL) were added onto slides and applied with cover slips to denature at 72 °C for 2 minutes. Slides were stored at 37 °C for 16–20 hours in a humidified slide moat. Slides were washed in 0.4x and 2x SSC with 0.05% Tween20, and SlowFade™ Diamond Antifade Mountant with DAPI (Invitrogen, Cat. No: S36964) was added followed by addition of cover slips (Fisherbrand, Cat. No: 12541036). Images were captured with an Echo Revolution Microscope at 60x magnification. Consistency was ensured by imaging from the same slide or two slides prepared from the same metaphase spread. Images were visualized with Fiji (v.2.1.0/1.53c)^53^. G-band karyotyping^19^ indicated that NCI-H2170 cells are polyploid (approximately triploid to hexaploid), whereas ERBB2, MYC and CDC6 amplification occur predominantly on ecDNA and not on HSRs. EcDNA copy number was quantified by counting extrachromosomal DNA foci in metaphase spreads from DNA FISH experiments, with approximately ∼150 metaphases analyzed per condition. Images were processed through a computer-vision based pipeline (Supplementary Methods, **Supplementary Figure 2, Supplementary Table 6**, https://github.com/Brunk-Lab/ecEnhance) and using a previously described AI-based image annotation pipeline^18^.

### Fluorescence-assisted cell sorting

FACS was used to isolate subpopulations of NCI-H2170, NCI-H716, and SNU16 cells with high and low expression of specific surface markers. HER2-FITC (Cell Signaling Technology, **Supplementary Table 1**) was used for NCI-H2170 cells, and FGFR2-670 (Cell Signaling Technology, **Supplementary Table 1**) was used for NCI-H716 and SNU16 cells. Cells were sorted to select the top ∼10% (high) and bottom ∼10% (low) based on fluorescence intensity. Live/dead discrimination was performed using Annexin V and Cytox Blue staining, and cells were gated to exclude debris and select live cells (**Supplementary Fig. 1,3,4**). Unsorted populations were used as controls, and post-sort purity was validated to ensure minimal overlap between high and low subpopulations (**Supplementary Fig. 3**). Sorting and analysis were performed using Becton Dickinson FACSAria II and FACSymphony S6 cytometers. Data analysis was performed using FlowJo software. Sorted cells were collected for downstream molecular and phenotypic assays.

### Real-time growth monitoring analysis

Cell growth was monitored using an xCELLigence real-time cell analyzer (Agilent) with RTCA Software Pro 2.6.0. This label-free approach monitors electrical impedance across microelectrodes, correlating with cell attachment, growth, and proliferation, with data recorded every 15 minutes over 192 hours (8 days) for high-resolution, time-course analysis. The SP station was placed in a humidified CO_2_ incubator, and background readings were recorded with 80 μL of media in a 96-well PET E-plate. For the FACS-sorted cells with high and low levels of HER2 cells and unsorted control cells, 17,000 cells were seeded per well in 100 μL of media. Impedance data were translated into normalized Cell Index values. Outer wells were filled with DPBS to minimize evaporation. To assess the impact of varying ecDNA levels on growth, ecDNA^high^ and ecDNA^low^ cells were mixed at 1:0, 0.25:0.75, 0.5:0.5, and 0:1 ratios, and growth was compared across these mixtures over 120 hours, with pre- and post-sorted control cells included for comparison. Impedance data were translated into normalized Cell Index values.

### Analytical flow cytometry and cell-cycle distribution analysis

NCI-H2170, SNU-16, and NCI-H716 cells were pulsed with 10 μM EdU (Santa Cruz Biotechnology, Cat # Sc-284628) for 1 hour at 37 °C to label actively dividing cells in S phase. Three independent cultures of each cell line were harvested and processed. After trypsinization and filtration through a 50-μm Celltrics filter (Sysmex), cells were washed with PBS, fixed in 4% paraformaldehyde (Sigma-Aldrich, Cat # 15714-S) for 15 minutes, and resuspended in 1% BSA-PBS. Cells were stored at 4 °C until further processing. For flow cytometry, cells were permeabilized with 0.5% Triton X-100 (Sigma-Aldrich, Cat # 9002-93-1) in 1% BSA-PBS for 15 minutes. EdU labeling was done by incubating cells with 1 μM Alexa 647-azide (Life Technologies, Cat # a10277), 1 mM CuSO_4_ (Sigma-Aldrich, Cat # 209198-100G), and 100 mM ascorbic acid (Fisher Scientific, Cat # A61-100) in PBS for 30 minutes in the dark at room temperature. NCI-H2170 cells were then incubated with 10 μL HER2-FITC antibody (Thermo Fisher Scientific, Cat # BMS120FI) at 1:10 dilution for 30 minutes in the dark. For DNA content analysis, cells were stained with 1 μg/mL DAPI (Life Technologies, Cat # D1306) and treated with 100 μg/mL RNAse A (Sigma-Aldrich, Cat # R5503) overnight at 4 °C or for 1 hour at 37 °C. Samples were analyzed using an Attune NxT flow cytometer (Thermo Fisher Scientific) with gating based on forward scatter area vs. side scatter area for cell selection, and DAPI area vs. height for singlet identification (**Supplementary Fig. 1**). Positive/negative gates for EdU and HER2-FITC were determined using unstained controls. FCS Express 7 Research (De Novo Software) was used for data analysis. In each biological replicate performed per cell line, at least 10,000 cells per sample were analyzed.

### Iterative indirect immunofluorescence imaging

4i was performed to analyze 29 proteins (**Supplementary Table 1-2**) across 12 staining and elution cycles. NCI-H2170, SNU16, and NCI-H716 cells were seeded onto poly-L-lysine-coated glass-bottomed 96-well plates for adhesion. For each cell line, two biological replicates were seeded into separate wells of a 96-well plate. Cells were fixed with 4% paraformaldehyde for 30 minutes, permeabilized with 0.1% Triton X-100 for 15 minutes, and blocked with 1% BSA in PBS supplemented with 100 mM maleimide and 100 mM NH₄Cl for 1 hour. Each cycle involved overnight incubation with primary antibodies in 1% BSA in PBS (cBS) at 4 °C, followed by secondary antibodies conjugated to fluorophores (1:500 in cBS) for 1 hour at room temperature. Hoechst 33258 (1:2500) was included for nuclear segmentation. After each cycle, fluorescent signals were removed by washing with 3 M guanidine chloride, 3 M urea, and 0.5 M glycine (pH 2.5) for three 10-minute washes. Fluorescence images were captured after each cycle, with overview images at 10x and detailed imaging at 100x magnification. For each biological replicate, multiple tiled fields of view were imaged across the well to capture large numbers of single cells. These tiles represent technical sampling of cells within a replicate rather than independent biological replicates, and all segmented cells were aggregated for downstream single-cell analysis. Signal intensity and antibody specificity were confirmed using positive and negative controls. After 14 cycles, samples were stored in PBS with 0.02% azide at 4 °C for long-term preservation. Composite protein program scores were derived from multiplexed 4i measurements to quantify proliferation and checkpoint-associated signaling. Protein intensities were residualized against per-cell global protein abundance to control for global staining differences, and standardized using z-score normalization. Proliferation (CDC6, Cyclin D1, E2F1, cFOS, MDM2, p21, MYC) and checkpoint/stress (FOXO3A, BRD4, pRb S807/811, cPARP, Cyclin A) program scores were computed as the mean of standardized residuals within each marker set. A net cell-cycle activity score was defined as the difference between proliferation and checkpoint scores. Differences between ecDNA^high^ and ecDNA^low^ cells (top and bottom 10% of HER2 intensity) were evaluated using Mann-Whitney tests.

### RNA isolation and digital qPCR sample preparation

High-throughput digital qPCR was performed to quantify gene expression across ecDNA and HSR model systems and in FACS-sorted subpopulations. Experiments included multiple cancer cell lines (NCI-H2170, SKGT2, SUM159, COLO320DM, COLO320HSR, SNU16, and NCI-H716) collected under control growth conditions. In addition, NCI-H2170 cells were analyzed following fluorescence-activated cell sorting (FACS) to isolate ecDNA-enriched populations based on HER2 expression (pre-sort, ecDNA^high^, ecDNA^low^, and post-sort fractions). Sorted populations were expanded for approximately ten generations prior to RNA analysis. Total RNA was isolated using the Zymo Quick-RNA Microprep Kit (Zymo Research, R1050). RNA concentration and purity were assessed by NanoDrop spectrophotometry and confirmed using the Qubit RNA High Sensitivity Assay (Thermo Fisher Scientific, Q32852). RNA concentrations were normalized to 20 ng/µL, and input volumes were calculated to provide ≥300 ng total RNA per sample. In total, 60 RNA samples spanning cell lines, time points, and sorted populations were analyzed. Reverse transcription was performed using the Standard BioTools protocol (PN 100-6472) to generate cDNA for subsequent amplification. Specific target amplification was carried out using pooled TaqMan assays (final concentration 180 nM) with the following cycling conditions: 2 minutes at 95 °C, followed by 20 cycles of 15 seconds at 95 °C and 4 minutes at 60 °C. Preamplified cDNA was diluted 1:5 with DNA Suspension Buffer and stored at −20 °C until analysis. Quantitative PCR was performed on the BioMark HD system (Standard BioTools, PN 100-6174) using a 192.24 Dynamic Array integrated fluidic circuit (IFC). Sample pre-mixes and TaqMan assays targeting a panel of 24 genes were prepared and loaded according to the manufacturer’s instructions. Thermal cycling conditions consisted of 60 seconds at 95 °C followed by 35 cycles of 5 seconds at 96 °C and 20 seconds at 60 °C. Each sample was measured in technical replicate. Gene expression profiles were compared across cell lines and sorted ecDNA subpopulations to assess transcriptional changes associated with ecDNA dosage.

### 10X Genomics multi-omics single-cell sequencing

To characterize the transcriptional and chromatin accessibility landscapes of NCI-H2170 cells, we performed multiomics single-cell sequencing using the Chromium Single Cell Multiome ATAC + Gene Expression platform (10X Genomics). Cells were analyzed before their third passage to minimize culture-induced artifacts. Single-cell suspensions were prepared as per the 10X Genomics protocol, ensuring >85% viability. Cell concentration and quality were assessed using a Countess II Automated Cell Counter (ThermoFisher). Approximately 10,000 cells per sample were loaded into the Chromium Controller for partitioning into Gel Bead-In Emulsions, enabling RNA and chromatin accessibility profiling in the same cells. Library preparation followed manufacturer guidelines, including reverse transcription for gene expression, transposition for chromatin accessibility, and amplification of both cDNA and ATAC libraries. Libraries were quantified using a Qubit dsDNA High Sensitivity Assay (ThermoFisher) and assessed for fragment size with an Agilent 4200 TapeStation. Sequencing was performed on an Illumina NovaSeq 6000 platform with paired-end reads. Data processing was done with Cell Ranger ARC (v2.0.1) and analyzed in R (v4.2.1/v4.3.1) using Seurat (v4.4.0)^54^ and Signac (v1.10.0)^55^. A Seurat object was created following the Signac protocol. Cells passing specific quality criteria were retained for further analysis, and DoubletFinder (v2.0.4) was used to identify and filter out doublets. ecDNA copy numbers were inferred from scATAC-seq data using a method adapted from a previous publication^38^. We compared distributional properties of ecDNA counts inferred from scATAC-seq, scDNA-seq, and FISH imaging (***Extended Data Fig. 4d-e***). Scripts are available at https://github.com/Brunk-Lab/ecMultiOME.

The 10X Single Cell Multiome ATAC + Gene Expression data of SNU16 was acquired from NCBI with SRA run accession number of SRR29521417 under the project accession number of PRJNA1127616. The 10X Single Cell Multiome ATAC + Gene Expression data of COLO320DM and COLO320HSR was acquired from NCBI under GEO accession number of GSE159986. The 10X single-cell multiome data of SNU16, COLO320DM, and COLO320HSR was aligned and processed similarly as NCIH2170.

The DNA/ATAC/RNA coverage track plots for MYC and PVT1 regions were made using the pseudobulk data of NCIH2170, SNU16, COLO320DM, SKGT2, KATOIII and COLO320HSR from 10X Single-cell multiome and Bioskryb ResolveOME platforms with the tools deeptools (v.3.5.6) and IGV (v.2.19.4). The coverage is normalized by CPM by setting the parameter “--normalizingUsing” as “CPM”.

### Nanopore long-read sequencing

Nanopore long-read sequencing^39^ with adaptive sampling was used to profile specific regions on chromosomes 17 and 8 in NCI-H2170 cells. Adaptive sampling enriched DNA fragments from these regions for targeted sequencing without physical enrichment steps. Two protocols were tested: a ligation-based protocol and a rapid sequencing protocol. For the ligation protocol, high-molecular weight DNA was extracted, processed to optimize read lengths, and libraries were prepared using ONT’s ligation kit. Sequencing was performed on MinION or PromethION platforms. The rapid protocol used the Oxford Nanopore Technologies (ONT) rapid sequencing kit. Sequencing runs applied adaptive sampling to target specific chromosomes, controlled by ONT’s algorithms in MinKNOW software. We compared enrichment efficiency, coverage depth, and read length distribution between the protocols. Basecalling was performed using Guppy (v.6.4.2; https://community.nanoporetech.com) in high-accuracy mode, and reads were aligned to GRCh38 with minimap2^56^ (v.2.26-r1175). NanoPlot (v.1.44.1) was used for quality control (**Supplementary Fig. 5**). Coverage was calculated for specific regions (chr8:125,000,000–130,000,000 and chr17:39,000,000–40,500,000) (**Fig. 2b, Supplementary Fig. 6**), and Flye^57^ (v.2.9.2) was used to assemble the consensus ecDNA structure. Variant calling was done using Clair3^58^ within a Singularity container, with alignment to GRCh38 using minimap2^56^.

The nanopore long-read WGS data of SNU16 was acquired from the public database NCBI with the SRA run accession number of SRR2951399 under the project accession number PRJNA1127616. The package CorRAL (v.2.2.0) was used to predict ecDNA structures in SNU16. The function “coral seed –cn-seg” was used to acquire the copy number profile across the genome in the cell line. The function “coral reconstruct” was used to reconstruct the ecDNA structures. The function “coral plot” was used to generate the ecDNA visualization plot. The package sniffles (v.2.6.3) was used to detect structural variants/breakpoints in the long-read WGS data of SNU16.

The nanopore long-read genomic DNA data of COLO320DM was acquired from the public database NCBI with the SRA run accession number of SRR12880625 under the project accession number PRJNA670737. Genomic coverage of the bin amplified on ecDNA used to estimate ecDNA counts was plotted similarly as NCIH2170.

### Single-cell ResolveOME multi-omics sequencing

Cells were prepared for FACS as described above. Single cells were isolated via FACS and plated into microtiter plates, ensuring one cell per well (based on ERBB2 intensity for NCI-H2170 and SKGT2 cells and FGFR2 intensity for SNU-16 and KATOIII cells). Paired DNA and RNA libraries were prepared from individual cells using the Bioskryb Genomics ResolveOME Whole Genome and Whole Transcriptome Single Cell Core Kit, according to the manufacturer’s instructions. Cytosolic lysis was performed to release RNA, and first-strand cDNA was synthesized from cytosolic mRNA via reverse transcription. Following nuclear lysis, whole genome amplification was conducted on the same cell using Primary Template-directed Amplification. The cDNA was then purified and amplified, and libraries were prepared. DNA and RNA sequencing was performed using the Illumina NextSeq 2000 platform. Paired-end 50-base-pair sequencing was conducted on both DNA (targeting 2×10^6^ reads) and RNA (targeting 2×10^5^ reads) arms per single-cell library. The analysis of the data is described in the Supplementary Methods.

### Machine learning

Single-cell RNA sequencing data were used to predict ecDNA copy numbers with 2600 genes (log-transformed transcript-per-million, logTPM) retained for analysis. Missing values were imputed using median imputation, and data were standardized before splitting into 80% training and 20% test sets. Four models were implemented: Random forest, gradient boosting, extreme gradient boosting (XGBoost^59^), and a neural network. The random forest model was trained with 500 trees, maximum depth of 15, and split constraints. The gradient boosting model was trained with 500 boosting stages, learning rate 0.05, and maximum depth 5. XGBoost used 500 rounds, a learning rate of 0.05, and regularization (L1: 0.1, L2: 1). The neural network, built in PyTorch, had three hidden layers (512, 256, 128 neurons) with Swish activation and early stopping after 20 epochs. Models were evaluated with MAE, R², Pearson, and Spearman correlation, with XGBoost performing the best (lowest MAE: 66, highest R²: 0.8). The trained XGBoost model was applied to new datasets after identical preprocessing, and predicted ecDNA counts were stored for analysis. All methods were implemented in Python 3.8+ using scikit-learn, XGBoost, PyTorch, and joblib, with random seeds for reproducibility. The pipeline, including scripts and trained models, is available upon request.

### Statistical analyses

To assess differences in ecDNA abundance across FISH data from FACS-sorted cells cultured at various time points (0, 48, 96, 432 hours), we applied both linear mixed effects modeling and ordinary least squares regression. For global analysis, a linear mixed effects model was applied with the formula: ecDNA ∼ Timeline * Group + (1 | folder), where “Timeline” indicates timepoints (G0, G1, G2, G9), “Group” reflects FACS-sorted populations (High, Low, Post-sort Control, Pre-sort Control), and “folder” accounts for batch variability. Post-sort Control at G0 was used as the reference group. In cases where Post-sort Control at specific timepoints was missing, synthetic rows were generated. Model fitting was done with maximum likelihood estimation using the L-BFGS optimizer. For subset comparisons, ordinary least squares models were applied to evaluate differences between High vs. Low populations at individual timepoints, with High as the reference level. Interaction models (ecDNA ∼ Timeline * Group) were also used to assess changes in group-level differences across timepoints. Estimated coefficients, standard errors, p-values, and confidence intervals were extracted, and p-values for specific combinations (e.g., G9 High vs. baseline) were computed using t_test() with custom contrast vectors. Group means and standard errors were visualized using bar plots, and Welch’s t-tests were applied for group differences. All analyses were conducted in Python using statsmodels, scipy, pandas, and seaborn. All measurements were obtained from biologically independent samples unless otherwise noted; no repeated measurements were taken from the same sample across conditions. All statistical tests were two-sided unless otherwise specified.

### Computational modeling and analysis

To infer ecDNA dosage without relying on prior amplicon annotations, we performed an unbiased genome-wide co-variation screen using single-cell DNA copy number profiles from ResolveOME data. For each genomic bin, we quantified copy number across individual cells and computed pairwise Spearman correlations between bins across the genome. We then identified genomic regions exhibiting the strongest continuous and coordinated co-variation patterns. Regions were ranked based on (i) magnitude of pairwise correlation (Spearman ρ), (ii) smoothness of variation across the population, and (iii) degree centrality within the genome-wide co-variation matrix. Clusters of bins showing high mutual correlation and forming a coherent module were considered candidate ecDNA-associated regions. For each model, the composite ecDNA dosage axis was defined as the aggregate copy number across the most strongly co-varying segments. This approach recovered the full ecDNA amplicon structure in NCI-H2170, SNU16 and COLO320DM cells, matching long-read assembly coordinates and DNA metaphase FISH measurements. Validation against DNA FISH confirmed that the inferred dosage axis accurately reflected ecDNA copy number distributions. Following inference of the ecDNA dosage axis, single cells were ordered along this continuous axis. For transcriptomic analyses, normalized gene expression values were correlated with the inferred dosage metric using Spearman correlation. Genes exhibiting significant monotonic scaling (Benjamini–Hochberg FDR < 0.05) were considered dosage-responsive. Single-cell gene set enrichment analysis score was performed using MSigDB hallmark pathways. For proteomic analyses using 4i data, single-cell protein intensities were log-transformed and batch-corrected prior to analysis. Correlations between protein abundance and ecDNA dosage proxies (HER2 or FGFR2) were computed to identify dosage-dependent scaling relationships. To assess continuous scaling, protein abundance values were ordered along the ecDNA proxy dosage axis and locally smoothed using a nonparametric regression approach (LOESS). Smoothed trajectories were used to identify monotonic and non-linear scaling relationships while reducing noise from single-cell variability. Correlation coefficients were computed between raw protein values and ecDNA dosage, and scaling patterns were visually confirmed using smoothed trends.

To determine whether ecDNA dosage reorganizes regulatory architecture, protein–protein correlation networks were constructed separately for ecDNA^high^ and ecDNA^low^ subpopulations. Pairwise Spearman correlations were calculated among all quantified proteins (n = 29). Significant correlations (FDR < 0.05) were retained as network edges. Network density, connected components, and clustering coefficients were computed to assess structural differences between subpopulations. Centrality metrics, including degree and betweenness centrality, were calculated using NetworkX. Statistical significance of network differences was evaluated by permutation testing (10,000 label shuffles), preserving protein distributions while randomizing ecDNA group assignments.

To characterize PVT1 isoform splice-junction usage across ecDNA dosage states, we analyzed full-length transcript reads generated by ResolveOME single-cell sequencing. The RNA-seq aligner STAR (v.2.7.11a) was used to align the FASTQ files from each single cell produced by Bioskryb ResolveOME platform to the hg38 genome. PVT1 splice junction reads spanning different exons were acquired from the “_SJ.out.tab” output file for each single cell. Genomic information for PVT1 exons was acquired from GRCh38. For each cell, junction spanning reads were quantified by counting uniquely mapped reads bridging annotated exon-exon boundaries. Junction diversity was quantified as the number of distinct exon-exon junctions detected per cell. Isoform-level patterns were visualized by aggregating junction counts across ecDNA^high^ and ecDNA^low^ subpopulations and across models. These single-cell splice-junction patterns were compared with bulk RNA-seq profiles across model systems to assess concordance. RNA-seq data of COLO320DM was acquired from GEO accession number GSE249656. RNA-seq data of SNU16 was acquired from GEO accession number GEO236987. The package StringTie (v.2.1.5) was used to predict different PVT1 isoforms from RNA-seq data of NCIH2170, COLO320DM, SKGT2 and COLO320HSR.

### Evolutionary modeling of ecDNA copy number and fitness dynamics

To investigate how ecDNA heterogeneity is maintained and restored across generations, we developed a stochastic population model based on the Gillespie algorithm^60^. Unlike prior short-term models focused primarily on co-segregation or clonal expansion^5,17,38^, our framework incorporates experimentally informed, ecDNA-dependent fitness effects and simulates population dynamics over multiple generations. Each cell is assigned an ecDNA copy number and ecDNA-dependent division and death rates. These rates are modeled as flexible log-linear functions of ecDNA dosage, allowing for non-linear fitness scaling that captures both positive and negative selection regimes. During simulation, birth and death events are selected probabilistically according to these rates. At division, duplicated ecDNA is partitioned between daughters using binomial sampling, with the daughter fraction p set to 0.5 (unbiased) or drawn from a biased distribution centered near 0.6, based on live imaging. To model experimentally observed asymmetry, we introduced a split inheritance parameter r, which permits biased ecDNA segregation during mitosis. To initialize simulations with realistic starting distributions and reduce sampling bias from FACS-sorted data, we implemented a Metropolis–Hastings upsampling procedure based on kernel density estimates of experimentally measured ecDNA distributions. This ensured that simulated populations matched observed baseline distributions prior to perturbation. Model parameters, including division rate, death rate, and segregation bias, were inferred using a genetic algorithm. Parameter sets were scored based on their ability to reproduce (i) the steady-state ecDNA distribution and (ii) the rapid recentralization of high and low populations toward the post-sort control distribution, quantified using Kullback–Leibler divergence. Iterative selection, mutation, crossover, and elitism were applied until convergence. The optimized model recapitulated both the experimentally observed rapid restoration of ecDNA heterogeneity and long-term stabilization of dosage distributions. Detailed equations, parameter bounds, and optimization procedures are provided in the Supplementary Methods. All simulations were implemented in Python, and the full codebase is available at: https://github.com/Brunk-Lab/ecRestore/.

### Live-cell Imaging and quantification of DNA partitioning

NCI-H2170 and RPE-hTERT cells were plated on cyclic olefin–bottom 96-well imaging plates (Greiner) and allowed to adhere for at least 24 hours prior to imaging. Cells were fixed with 4% formaldehyde (Electron Microscopy Sciences), permeabilized with 0.5% Triton X-100, and stained with 1 µg mL⁻¹ DAPI for 15 minutes at room temperature to visualize nuclear DNA. Images were acquired on a Nikon TiE inverted microscope equipped with a spinning-disk confocal system (Crest Optics X-light) and a Hamamatsu ORCA Flash 4.0 camera. For each field, seven confocal z-sections were collected using a 20× objective (NA 0.8). Fields were selected randomly across wells to avoid sampling bias. Image stacks were processed in FIJI to generate average-intensity projections and nuclei were segmented using the StarDist plugin. Post-mitotic daughter cells were manually identified based on nuclear morphology and DAPI intensity patterns indicative of recent cytokinesis. For each mitotic event, integrated nuclear fluorescence intensity was measured for both daughter nuclei. Partitioning asymmetry was quantified by calculating the fraction of total DNA signal inherited by each daughter cell. Distributions of daughter-cell intensity ratios were compared to simulated binomial expectations under symmetric inheritance (r = 0.5) to evaluate deviations consistent with asymmetric DNA partitioning.

## Acknowledgements

The authors thank the UNC Flow Cytometry Core, with special acknowledgment to Janet Dow and Ramiro Diz, for their expertise and support. We also extend our gratitude to staff of the UNC Microscopy Core and the Advanced Analytics Core, especially Gabrielle Cannon, for their valuable contributions. J.C., O.C., Y.W., and P.B. were supported by funding from the IBM Junior Faculty Development Grant, the Computational Medicine Pilot Grant, the PhRMA Foundation and the Lung Cancer Research Foundation Boehringer Ingelheim Early Career Investigator Award. J.G.C. and D.F are supported by the National Institute of General Medical Sciences (NIGMS) of the National Institutes of Health (NIH) under award number R35GM141833. D.F. also acknowledges support from the American Heart Association fellowship (Award No.1027147). T.E. is supported by the Maximizing Investigators’ Research Award (MIRA) from the NIH/NIGMS under award number R35GM127145. J.E.P., S.W, J.G.C., and P.S. are supported by the National Cancer Institute of the NIH under award number R01CA280482. The UNC Flow Cytometry Core Facility (RRID:SCR_019170) is supported in part by P30 CA016086 Cancer Center Core Support Grant to the UNC Lineberger Comprehensive Cancer Center. The UNC Advanced Analytics Core Facility is supported by the CGIBD center grant, P30 DK034987.

## Author Contributions

Conceptualization, E.B.; methodology, E.B., C.G.F., J.P., J.W., J.G.C.; formal Analysis, Y.W., O.C., J.C., A.M., D.F., C.G.F., P.B., S.H., S.G., L.S., E.B.; funding Acquisition, E.B.; investigation, Y.W., O.C., J.C., A.M., D.F., C.G.F., P.B., S.H., S.G., E.B.; resources, E.B., J.W., J.P., J.G.C; supervision, E.B., S.W., J.P., P.S., T.E., C.A.T., A.M.R., T.T., J.M.D., J.W., J.G.C.; validation, Y.W., O.C., J.C., D.F.; visualization, Y.W., J.C., D.F., E.B.; writing—original draft, E.B.; writing—review and editing, All authors. All authors have read and agreed to the published version of the manuscript.

## Inclusion and Ethics Statement

All experiments involving cell lines were conducted in compliance with institutional biosafety and research integrity guidelines approved by the University of North Carolina at Chapel Hill. No human or animal subjects were involved in this study. We are committed to fostering a supportive research environment. Our team reflects a range of backgrounds and training levels, and we actively mentor junior scientists across disciplines. We strive to ensure that our research practices are transparent, reproducible, and accessible to the broader scientific community. Data and code from this work are openly shared to promote collaboration and accountability.

## Extended Data Figures

**Ex. Data Fig. 1:**
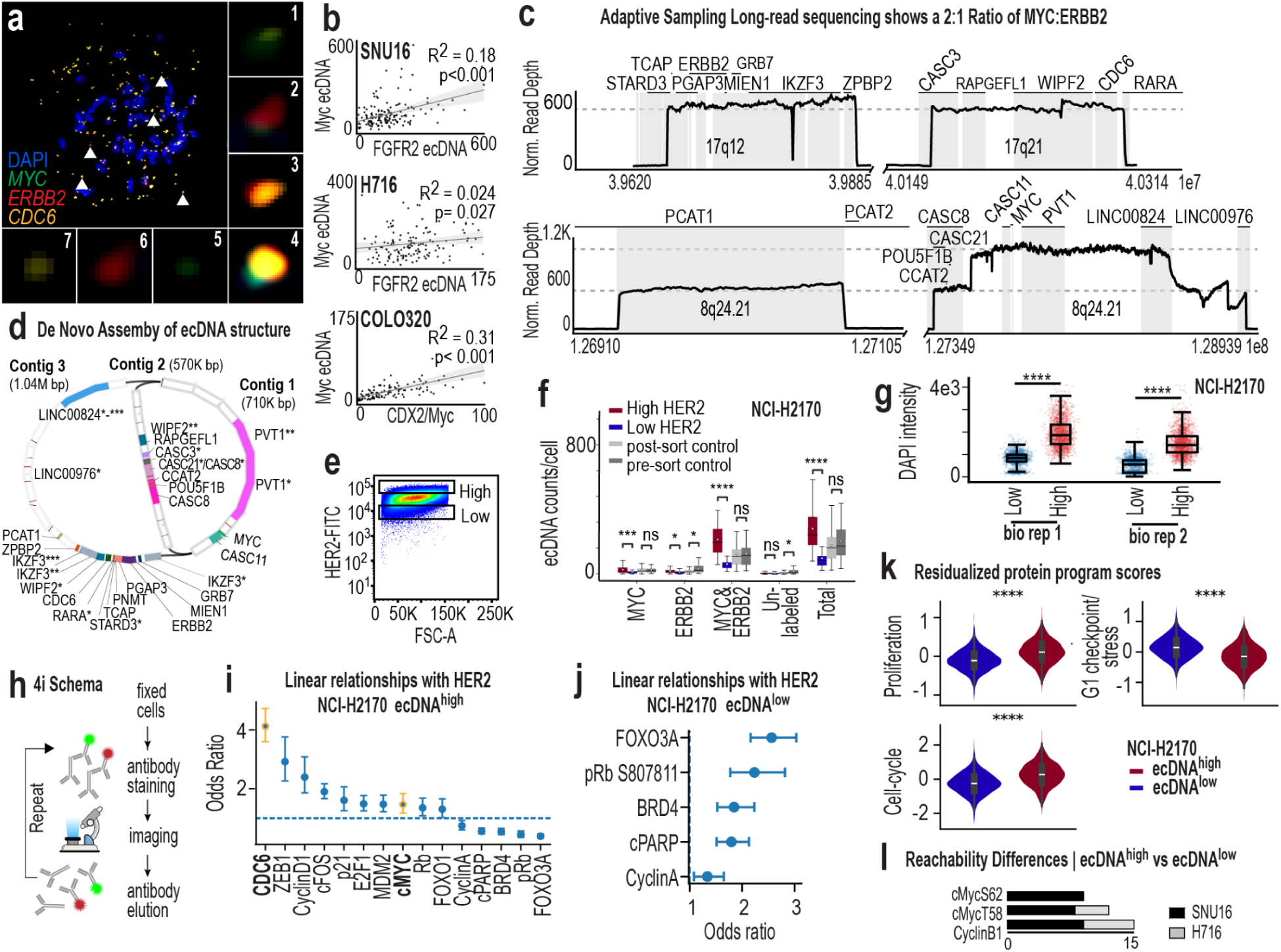
Characterization of ecDNA sequence and its relationship to protein expression. **a**, Tri-color DNA metaphase FISH image showing seven distinct ecDNA species in NCI-H2170 cells, with *MYC*, *ERBB2*, and *CDC6* amplified either individually or co-amplified on shared amplicons. **b,** Scatterplots illustrating variance in ecDNA dosage when two or more ecDNA species are present (top: SNU16 with *MYC* and *FGFR2* ecDNA species; middle: NCI-H716 with *MYC* and *FGFR2* ecDNA species; bottom: COLO320DM with *MYC* and *CDX2* ecDNA species). **c,** Oxford Nanopore adaptive long-read sequencing of chromosome 17 and chromosome 8 regions in NCI-H2170 cells, highlighting genomic segments amplified on ecDNA. **d,** *De novo* assembly of adaptive long-read sequencing data showing the predicted structure of the dominant ecDNA amplicon in NCI-H2170 cells. **e,** Flow cytometry plot showing FITC-HER2 intensity (y-axis) and forward scattering area, correlating to cell size (x-axis). Boxes indicate the gating strategy used to isolate ecDNA^high^ and ecDNA^low^ cells based on the top 13.7% and bottom 13.7% of HER2 intensity, respectively. Representative of three independent biological replicates. **f,** Box plots showing quantitative ecDNA counts from DNA metaphase FISH. NCI-H2170 cells were sorted into ecDNA^high^, ecDNA^low^, and control groups based on HER2 fluorescence intensity, and ecDNA copies were manually quantified. Distinct ecDNA species are shown separately, including species co-amplifying *MYC* and *ERBB2*, species amplifying these genes independently, and unlabeled species. Data representative of three biological replicates. **g,** Differences in total DNA between ecDNA^high^ and ecDNA^low^ NCI-H2170 cells using iterative indirect immunofluorescence imaging, representative of two biological replicates. **h,** Schema illustrating iterative indirect immunofluorescence imaging (4i). Fixed cells were sequentially labeled with antibodies, imaged, chemically stripped, and re-labeled across multiple rounds. Twenty-nine proteins were profiled in NCI-H2170, SNU16, and NCI-H716 cells. **i,** Forest plot showing odds ratios and 95% confidence intervals from a logistic regression model predicting ecDNA^high^ cells based on single-cell protein measurements from the 4i panel. Points indicate estimated odds ratios for each protein feature and horizontal bars denote 95% confidence intervals. The dashed vertical line indicates no association (odds ratio = 1). Proteins with odds ratios greater than 1 are positively associated with the ecDNA^high^ state. **j,** Forest plot showing odds ratios reoriented to represent enrichment in ecDNA^low^ cells from the same logistic regression model. Odds ratios were inverted so that values greater than 1 indicate stronger association with the HER2-low state. Points indicate estimated odds ratios and horizontal bars represent 95% confidence intervals. The dashed vertical line indicates no association (odds ratio = 1). **k,** Residualized protein program scores derived from multiplexed 4i measurements are shown for ecDNA^high^ and ecDNA^low^ cells. Proliferation markers (CDC6, Cyclin D1, E2F1, cFOS, MDM2, p21, MYC) and checkpoint/stress-associated markers (FOXO3A, BRD4, phosphorylated Rb, cPARP, Cyclin A) were aggregated to generate composite program scores. The net cell-cycle activity score represents the difference between proliferation and checkpoint program scores. Statistical comparisons were performed using two-sided Mann–Whitney tests. **l,** Network modeling of 4i protein data. Bar plots show differences in reachability properties for key regulatory proteins, including MYC post-translational modifications, between ecDNA^high^ and ecDNA^low^ states in SNU16 and NCI-H716 cells.

**Ex. Data. Fig. 2:**
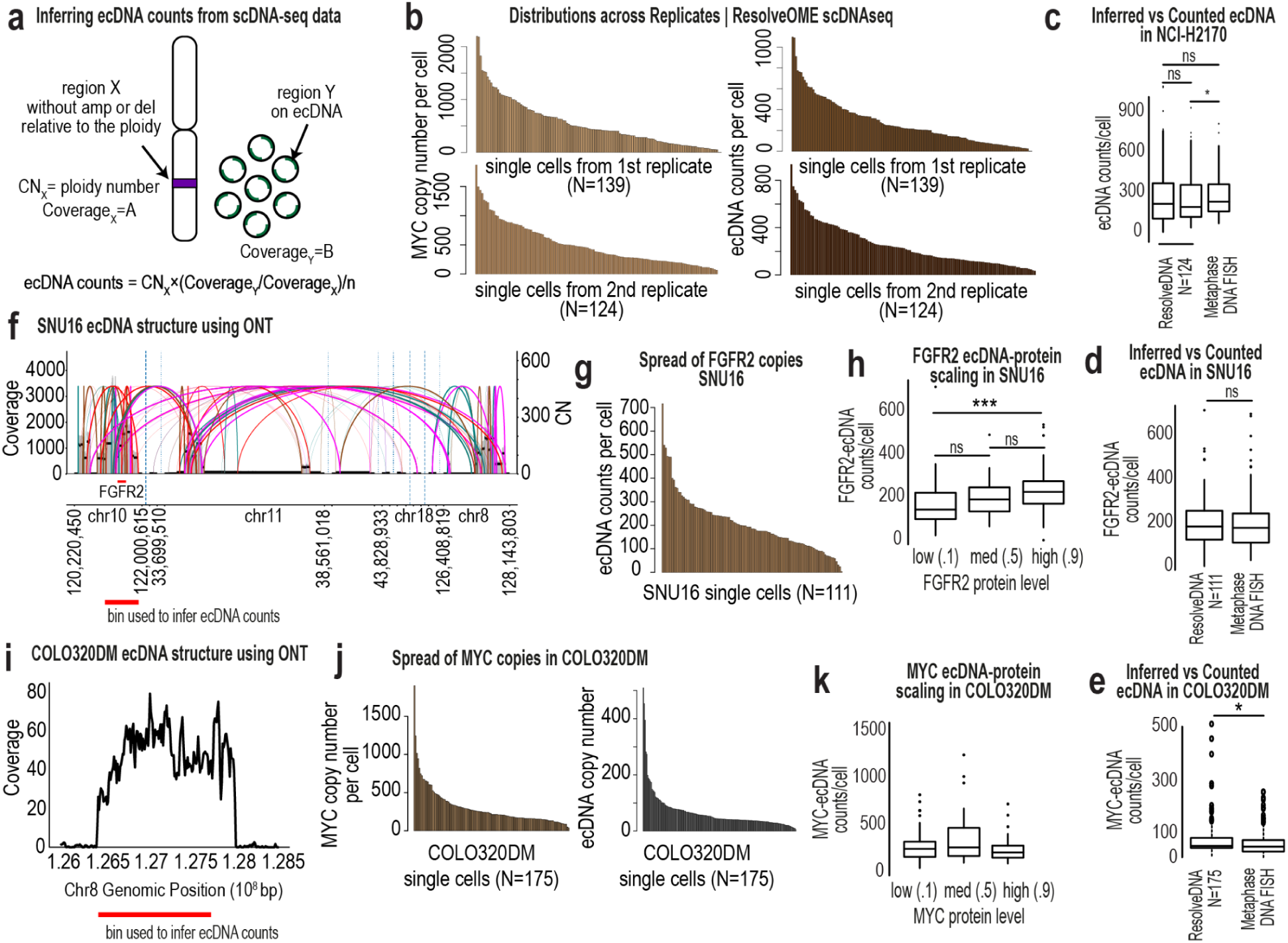
Functional and structural studies in multiple cell line models with ecDNA. **a**, Schema illustrating inference of ecDNA copy number from single-cell copy number variation profiles generated using ResolveOME. A chromosomal reference region (region X) with copy number equal to ploidy (CNₓ) and sequencing coverage A is used to normalize an amplified region (region Y) with coverage B. **b,** Distribution of MYC copy numbers across single cells profiled by ResolveOME in NCI-H2170 cells, shown for two independent replicates (left, replicate 1 top; replicate 2 bottom). Right, inferred ecDNA copy number distributions for the same samples. **c,** Box plots comparing inferred ecDNA copy numbers from ResolveOME scDNA data and counts obtained from DNA metaphase FISH. Wilcoxon rank-sum testing shows no significant difference between inferred and FISH-measured ecDNA counts. The ResolveDNA boxplots represent two replicates of NCI-H2170 sequenced cells. **d,** Distribution of inferred *FGFR2*-ecDNA copy numbers in 150 SNU16 cells estimated from ResolveOME data and quantified by DNA metaphase FISH. Wilcoxon rank-sum testing shows no significant difference between methods. **e,** Box plots comparing inferred ecDNA copy numbers from ResolveOME scDNA data with DNA metaphase FISH quantification in 150 COLO320DM cells. **f,** *De novo* assembly of ecDNA in SNU16 cells using available Oxford Nanopore long-read sequencing data and CoRAL structural reconstruction. **g,** Distribution of inferred *FGFR2*-ecDNA copy numbers across single SNU16 cells using genomic bins derived from the assembled ecDNA structure in (f). **h,** Box plots showing the relationship between inferred *FGFR2*-ecDNA copy number and FGFR2 protein abundance measured by flow cytometry prior to plating and sequencing 111 single cells. Wilcoxon rank-sum testing indicates significant differences between ecDNA^high^ and ecDNA^low^ cells where indicated. **i,** Identification of genomic bins corresponding to amplified ecDNA regions in COLO320DM cells using prior Oxford Nanopore sequencing. **j,** Distribution plots showing *MYC* copy number estimated from genomic bins identified in (i) and inferred ecDNA copy number across single COLO320DM cells from ResolveOME scDNA data. **k,** Box plots showing the relationship between inferred *MYC*-ecDNA copy number and MYC protein abundance measured by flow cytometry prior to single-cell plating and sequencing. Wilcoxon rank-sum testing indicates no significant protein differences were detected.

**Ex. Data Fig. 3:**
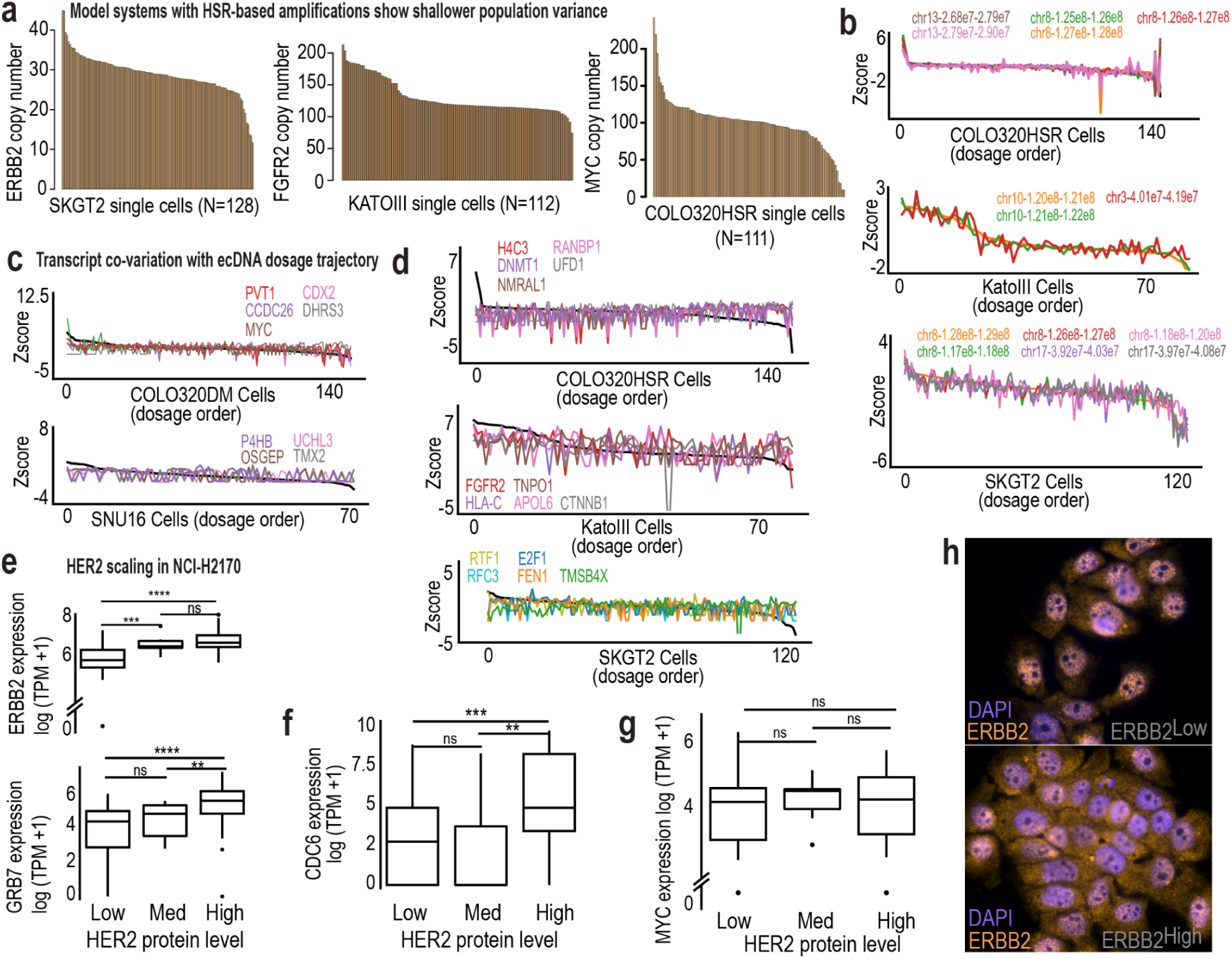
Single-cell relationships between ecDNA dosage and gene expression across models. **a**, Distribution of HSR-amplified gene copy numbers across single cells in SKGT2 (left), KATOIII (middle), and COLO320HSR (right) cells, illustrating the narrower and more uniform copy number spread characteristic of chromosomal amplifications. **b,** Ordering of single cells ranked by HSR-amplified copy number (x-axis) with z-scored DNA bin intensity shown on the y-axis. Cells with higher copy number are positioned on the left and lower copy number on the right. Co-varying genomic bins are overlaid to illustrate coordinated copy-number changes along the axis. Top, COLO320HSR; middle, KATOIII; bottom, SKGT2. **c,** Co-variation of transcript abundance with ecDNA dosage in ecDNA models COLO320DM (top) and SNU16 (bottom), demonstrating graded RNA responses across the copy number continuum. **d,** Co-variation of transcript abundance along MYC dosage trajectory in COLO320HSR (top), KATOIII (middle, FGFR2), and SKGT2 (bottom, ERBB2/MYC), showing comparatively reduced transcriptional spread. **e–h,** Box plots showing relationships between transcript levels and HER2 protein abundance measured by flow cytometry in 97 NCI-H2170 single cells profiled by ResolveOME. **e-g,** Boxplots showing transcript levels of ERBB2, GRB7, CDC6, and MYC across single cells binned by HER2 protein intensity (low, medium, high; n = 139 cells). Boxes represent the interquartile range with median indicated; whiskers denote 1.5× IQR. Statistical significance between groups was assessed using the Wilcoxon rank-sum test. **h,** Representative RNA-DNA FISH images illustrating heterogeneity in ERBB2 transcript signal across individual cells.

**Ex. Data. Fig. 4:**
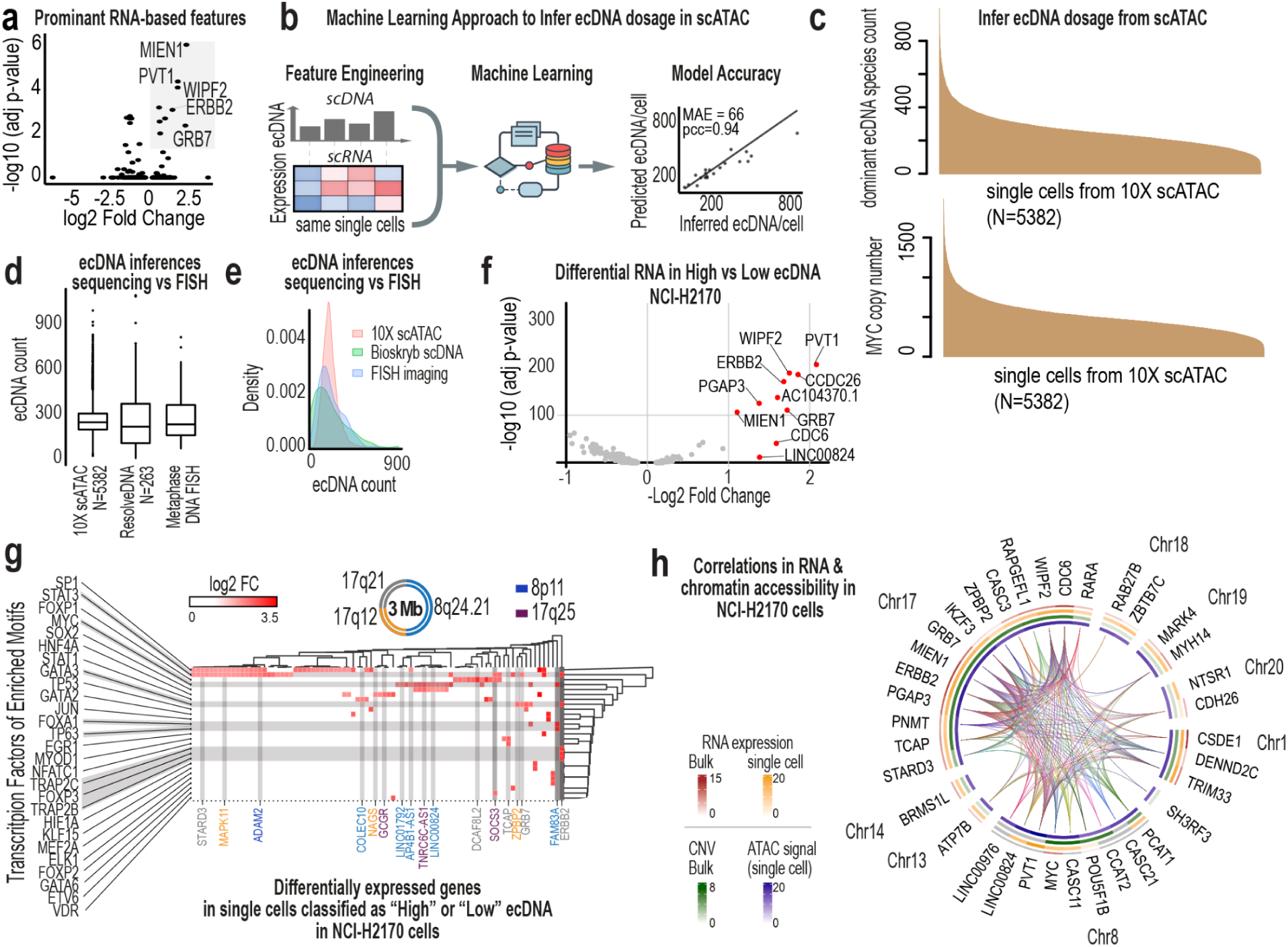
Predictive and multimodal structure linking ecDNA dosage to transcription and chromatin state. **a**, Volcano plot of differential gene expression between ecDNA^high^ and ecDNA^low^ NCI-H2170 cells profiled by ResolveOME. Highlighted genes represent the most significant transcript-level differences associated with ecDNA dosage. **b,** Supervised machine learning framework for predicting ecDNA copy number from matched single-cell DNA and RNA data. An XGBoost model was trained using transcript levels of the top ecDNA-associated RNA features identified in (a). Predicted versus inferred ecDNA counts per cell are shown. The model achieved high accuracy (MAE = 66 copies per cell; Pearson r = 0.94), demonstrating strong predictive coupling between transcriptional state and ecDNA dosage. **c,** Application of the trained model to independent 10X Multiome datasets. Top, distribution of predicted ecDNA copy numbers inferred from scATAC profiles. Bottom, computed MYC copy number per cell derived from scATAC signal. **d,** Box plots comparing ecDNA copy number estimates derived from scATAC data (10X), scDNA data (ResolveOME), and DNA metaphase FISH (N=150). **e,** Kernel density plots corresponding to (d), illustrating distributional differences across platforms. scDNA-based inference closely recapitulates FISH measurements, whereas scATAC-based estimates show narrowed variance. **f,** Volcano plot of differential gene expression between ecDNA^high^ and ecDNA^low^ cells using 10X Multiome data, confirming reproducibility of dosage-associated transcriptional shifts across platforms. **g,** Heatmap linking differentially expressed genes in ecDNA^high^ versus ecDNA^low^ cells (10X data) to transcription factors enriched for binding motifs in accessible chromatin regions. Axes represent differentially expressed genes and motif-enriched transcription factors; intersections indicate coordinated regulatory relationships. **h,** Multimodal integration of gene expression and chromatin accessibility in NCI-H2170 cells. Arcs denote significant gene–peak correlations. Inner circular heatmaps show concordance between bulk and single-cell RNA and ATAC signals, illustrating coordinated epigenetic remodeling associated with ecDNA dosage.

**Ex. Data. Fig. 5:**
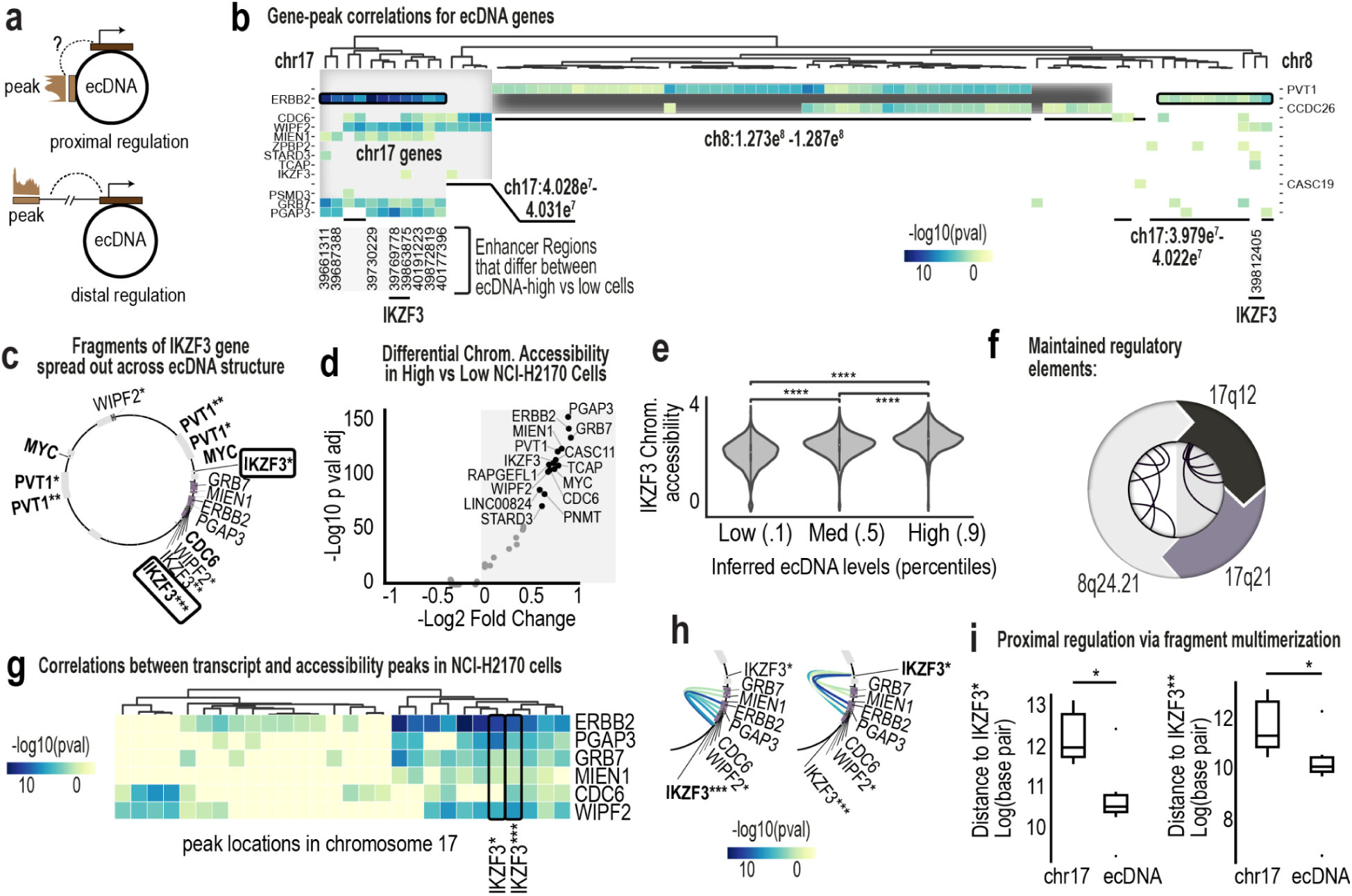
ecDNA reshapes cis-regulatory architecture and enhancer proximity. **a**, Schematic illustrating proximal and distal modes of epigenetic regulation. EcDNA multimerization can juxtapose regulatory elements that were originally separated on chromosomes, enabling both local and long-range interactions. **b,** Heatmap of gene–peak correlations for ecDNA-amplified genes in NCI-H2170 cells. Color intensity reflects correlation strength between chromatin accessibility and gene expression. Hierarchical clustering reveals regulatory modules. Although chromosome 8 and chromosome 17 genes co-amplify on the same ecDNA, most cluster according to their chromosome of origin, indicating preserved cis-regulatory organization. IKZF3 exhibits the highest number of gene–peak correlations. **c,** Structural model of the dominant NCI-H2170 ecDNA amplicon showing multimerized fragments of IKZF3 distributed across distinct positions within the circular structure, creating novel breakpoint configurations. **d,** Volcano plot of differential chromatin accessibility between ecDNA^high^ and ecDNA^low^ NCI-H2170 cells. Peaks associated with ecDNA-amplified loci are among the most significantly altered regions. **e,** Violin plots showing IKZF3 chromatin accessibility across ecDNA^low^, ecDNA^medium^, and ecDNA^high^ subpopulations defined by inferred ecDNA levels. Accessibility scales with ecDNA dosage. **f,** Schema showing preserved cis-regulatory organization, according to the chromosome of origin. **g,** Zoomed heatmap of gene–peak correlations on chromosome 17 ecDNA regions. IKZF3 fragments flanking ERBB2, GRB7, MIEN1, and related genes show the strongest and most frequent correlations. **h,** Arc diagram connecting IKZF3 fragments to chromosome 17 ecDNA genes with which their accessibility is significantly correlated. Strongest correlations occur with genes that are spatially proximal on the ecDNA amplicon. **i,** Boxplots comparing physical, linear genomic distances between IKZF3 fragments and highly correlated chromosome 17 genes on ecDNA versus their original chromosomal configuration. Distances are significantly shorter on ecDNA, indicating that multimerization repositions regulatory elements to enhance local transcriptional coupling.

**Ex. Data. Fig. 6:**
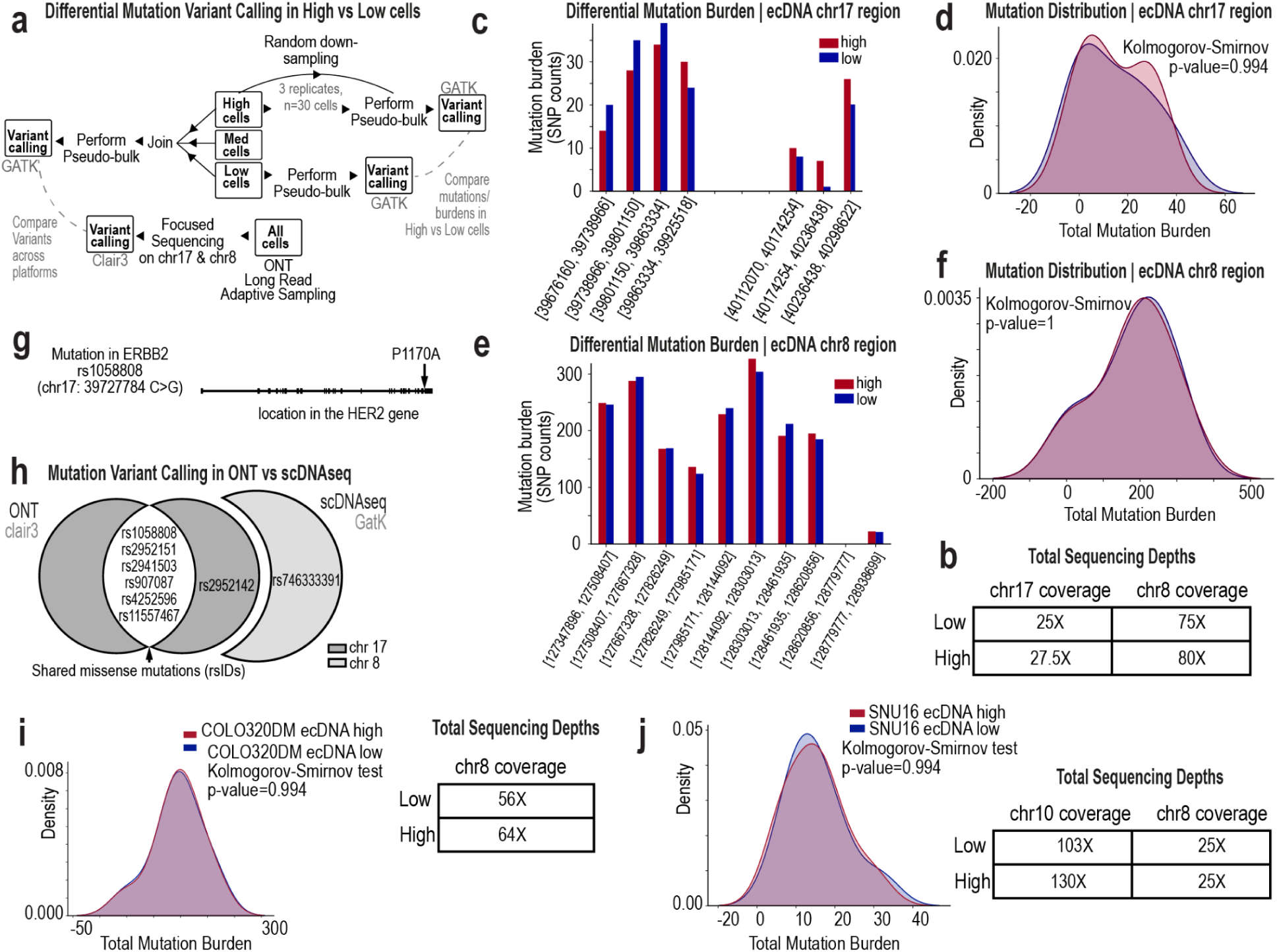
EcDNA-driven phenotypic variation occurs without changes in mutational burden. **a**, Schematic of the variant-calling workflow used to compare ecDNA^high^ vs ecDNA^low^ NCI-H2170 cells following FACS isolation. Single-cell DNA profiles were aggregated to generate pseudobulk variant calls within ecDNA-amplified regions. **b,** Sequencing depth across chromosome 17 and chromosome 8 ecDNA regions in ecDNA^high^ and ecDNA^low^ cells cells after downsampling to equalize coverage. **c,** SNP counts per genomic bin across the chromosome 17 ecDNA region in FACS-sorted ecDNA^high^ (N=30) and ecDNA^low^ (N=64) cells. **d,** Distribution of SNP counts within the chromosome 17 ecDNA region showing no significant difference in mutational burden between ecDNA^high^ and ecDNA^low^ cells. **e,** SNP counts per genomic bin across the chromosome 8 ecDNA region in ecDNA^high^ and ecDNA^low^ cells. **f,** Distribution of SNP counts within the chromosome 8 ecDNA region again demonstrating no significant difference in mutational burden between states. **g,** Detection of a functionally relevant *ERBB2* missense variant present in both ecDNA^high^ and ecDNA^low^ cells. Exons are shown as blocks along the gene model. **h,** Concordance of missense variants identified by long-read Oxford Nanopore sequencing and ResolveOME single-cell DNA sequencing. **i,** Distribution of SNP counts within the chromosome 8 ecDNA region of COLO320DM cells demonstrating no significant difference in mutational burden between ecDNA^high^ and ecDNA^low^ cells. Sequencing depth across chromosome 8 regions in ecDNA^high^ and ecDNA^low^ cells after downsampling to equalize coverage. **j,** Distribution of SNP counts within the chromosome 8 ecDNA region of SNU16 cells demonstrating no significant difference in mutational burden between ecDNA^high^ and ecDNA^low^ cells. Sequencing depth across chromosome 8 regions in ecDNA^high^ and ecDNA^low^ cells after downsampling to equalize coverage.

**Ex. Data. Fig. 7:**
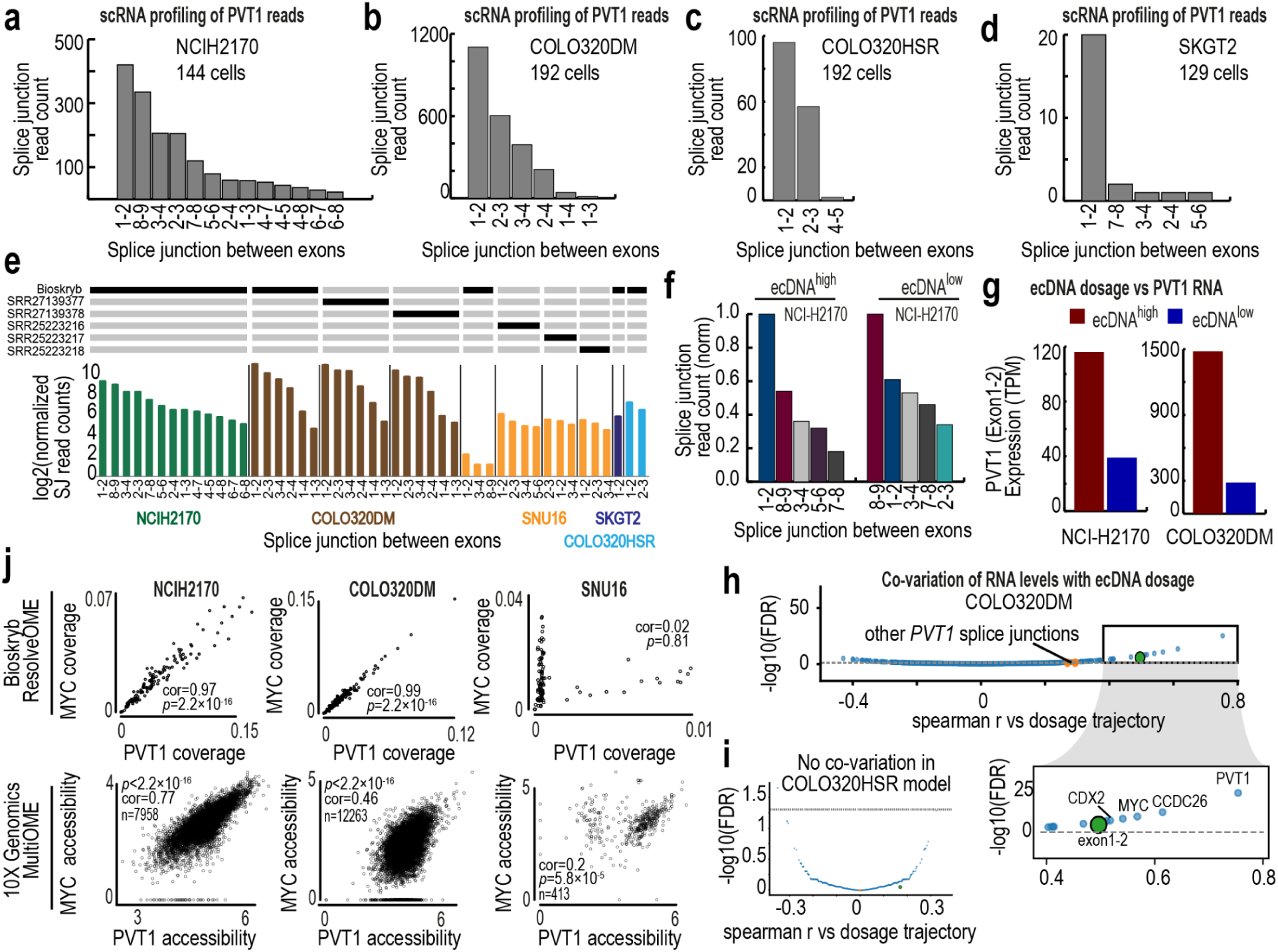
PVT1 splice architecture and dosage-dependent coupling vary by amplification architecture. **a**, Distribution of PVT1 splice-junction reads in NCI-H2170 single-cell RNA sequencing data. PVT1 is amplified on ecDNA in this model. The x-axis shows exon–exon junctions and the y-axis shows splice-junction read counts. **b,** Distribution of PVT1 splice-junction reads in COLO320DM cells, where PVT1 is amplified on ecDNA. **c,** Distribution of PVT1 splice-junction reads in COLO320HSR cells, where PVT1 is amplified on homogeneous staining regions. **d,** Distribution of PVT1 splice-junction reads in SKGT2 cells, where PVT1 resides on HSRs. **e,** Distribution of PVT1 splice-junction reads across model systems. Comparisons are drawn between Bioskryb pseudobulk data and pre-existing bulk RNA-sequencing data. **f,** Differences in PVT1 splice-junction read counts between 64 ecDNA^high^ and 64 ecDNA^low^ NCI-H2170 cells. **g,** Exon 1–2 PVT1 expression (TPM) predicted from StringTie in ecDNA^high^ and ecDNA^low^ cells from NCI-H2170 (left, N=128) and COLO320DM (right, N=106). **h,** Association volcano plot of Spearman correlation versus –log10(FDR) for RNA features co-varying with the ecDNA dosage axis in COLO320DM cells. Exon 1–2 of PVT1 is among the strongest and most significant dosage-correlated features. Inset highlights the most significant transcripts, including ecDNA-amplified genes. **i,** Equivalent analysis in COLO320HSR cells, showing minimal significant co-variation with MYC dosage trajectory. **j,** Cross-model comparison of *MYC* and *PVT1* coupling. Top, scDNA coverage relationships between *MYC* and *PVT1* from ResolveOME data (N=139 for NCI-H2170, N=175 for COLO320DM, N=111 for SNU16). Bottom, chromatin accessibility relationships from 10X Multiome data. NCI-H2170 and COLO320DM show strong linear coupling, whereas SNU16 exhibits a bimodal, two-state pattern.

**Ex. Data. Fig. 8:**
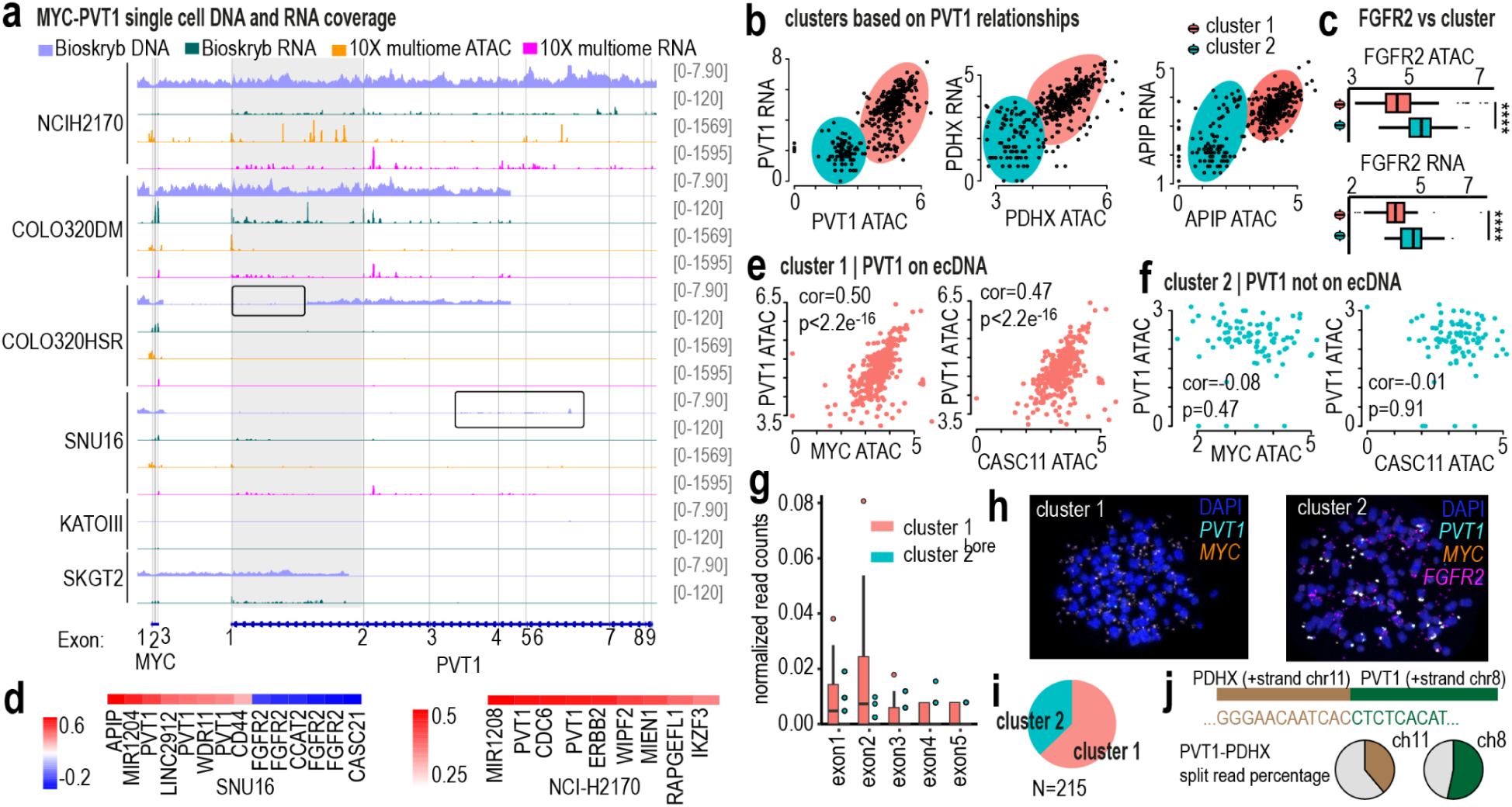
Architecture-dependent decoupling of PVT1 across genomic, epigenetic, and transcriptional layers. **a**, Integrated genomic view of MYC and PVT1 loci across six cell lines. Tracks show scDNA coverage (ResolveOME pseudobulk data), scRNA coverage (ResolveOME), scATAC coverage (10X Multiome), and scRNA coverage (10X Multiome). Highlighted regions indicate segments of PVT1 with reduced scDNA coverage in specific models, suggesting structural heterogeneity in amplification. **b,** Scatter plots from SNU16 10X Multiome data (N=413) showing RNA expression (CPM) versus chromatin accessibility for PVT1 (left), PDHX (middle), and APIP (right). Each plot reveals two distinct cellular clusters: one exhibiting coordinated accessibility–expression coupling and another lacking this relationship. **c,** FGFR2 chromatin accessibility (top) and RNA expression (bottom) stratified by the two clusters identified in (b), demonstrating differential regulatory behavior between states in SNU16 cells. **d,** Heatmap of promoter peak–gene correlations associated with PVT1 RNA levels. Left, SNU16; right, NCI-H2170. In SNU16, FGFR2 promoter accessibility negatively correlates with PVT1 RNA, consistent with cluster separation in (c). This antagonistic relationship is absent in NCI-H2170, indicating architecture-specific regulatory wiring. **e,** Cluster 1 cells (N=324) in SNU16 showing positive relationships between PVT1 ATAC signal and MYC (left) or CASC11 (right), consistent with co-amplification on ecDNA. **f,** Cluster 2 cells (N=89) in SNU16 showing loss of correlation between PVT1 ATAC and MYC or CASC11 accessibility, consistent with absence of PVT1 on ecDNA in this subset. **g,** Normalized splice-junction read counts from Bioskryb SNU16 data for PVT1 exons in cluster 1 (N=15) versus cluster 2 (N=96) cells. Only cluster 1 cells display robust exon 1–2 junction usage, supporting ecDNA-associated transcriptional activity. The small number of cluster 1 cells likely reflects enrichment of the dataset for FGFR2-positive cells during FACS isolation, and PVT1 and FGFR2 signals appear partially anticorrelated in this model. **h,** Tri-color DNA metaphase FISH in SNU16 cells showing heterogeneous amplification states: some cells lack PVT1 on ecDNA, while others co-amplify PVT1 with MYC on ecDNA and amplify FGFR2 on a separate amplicon. **i,** Quantification of 215 metaphase FISH images showing that 63% of cells harbor PVT1 on ecDNA, while 37% lack PVT1 amplification on ecDNA. **j,** Quantification of PVT1-PDHX split read percentage (breakpoints) at specific loci using Oxford Nanopore long-read WGS data. Two genomic locations were quantified at chr11:350,580,77 and chr8:127,949,634. Only 35-48% of reads show sequences that indicate PVT1-PDHX breakpoints are occurring, which indicates that not all PVT1 copies are amplified on ecDNA.

**Ex. Data. Fig. 9:**
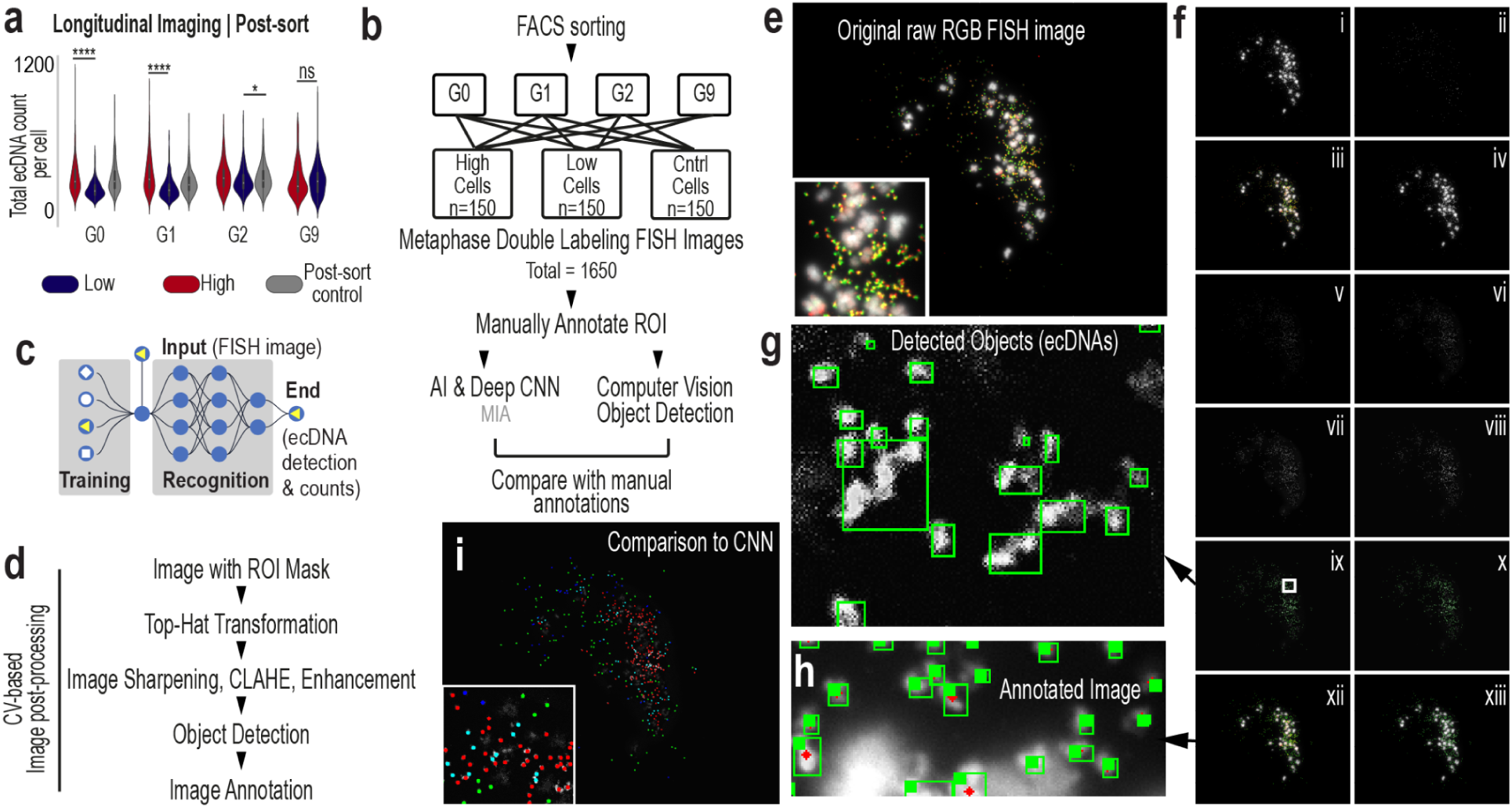
Automated Object Detection Accelerates Longitudinal Imaging of ecDNA variation dynamics. **a**, Differences between high, low, and control cells diminish rapidly in subsequent generations post-sorting. **b**, Schema of complete sorting, longitudinal imaging, and quantification strategy to quantify ecDNA variation restoration. **c**, Schema of AI model trained to automatically annotate DNA metaphase FISH images using convolutional neural networks (CNN). **d**, Steps involved in the image post-processing approach. **e**, Initial step of the image post-processing pipeline using a raw RGB DNA metaphase FISH image of a NCI-H2170 bursted nuclei. Fluorescent probes label ERBB2 and MYC genes, amplified solely on ecDNA in this model system. **f**, DNA metaphase FISH image transformation and enhancement steps within the pipeline. **g**, Example of ecDNA objects detected within DNA metaphase FISH image using computer vision methods. **h**, Final annotated image showing detected ecDNA in DAPI-stained DNA metaphase FISH images. **i**, Comparison between computer vision-based object detection and convolutional neural network (CNN)-based annotation methods.

**Ex. Data. Fig. 10:**
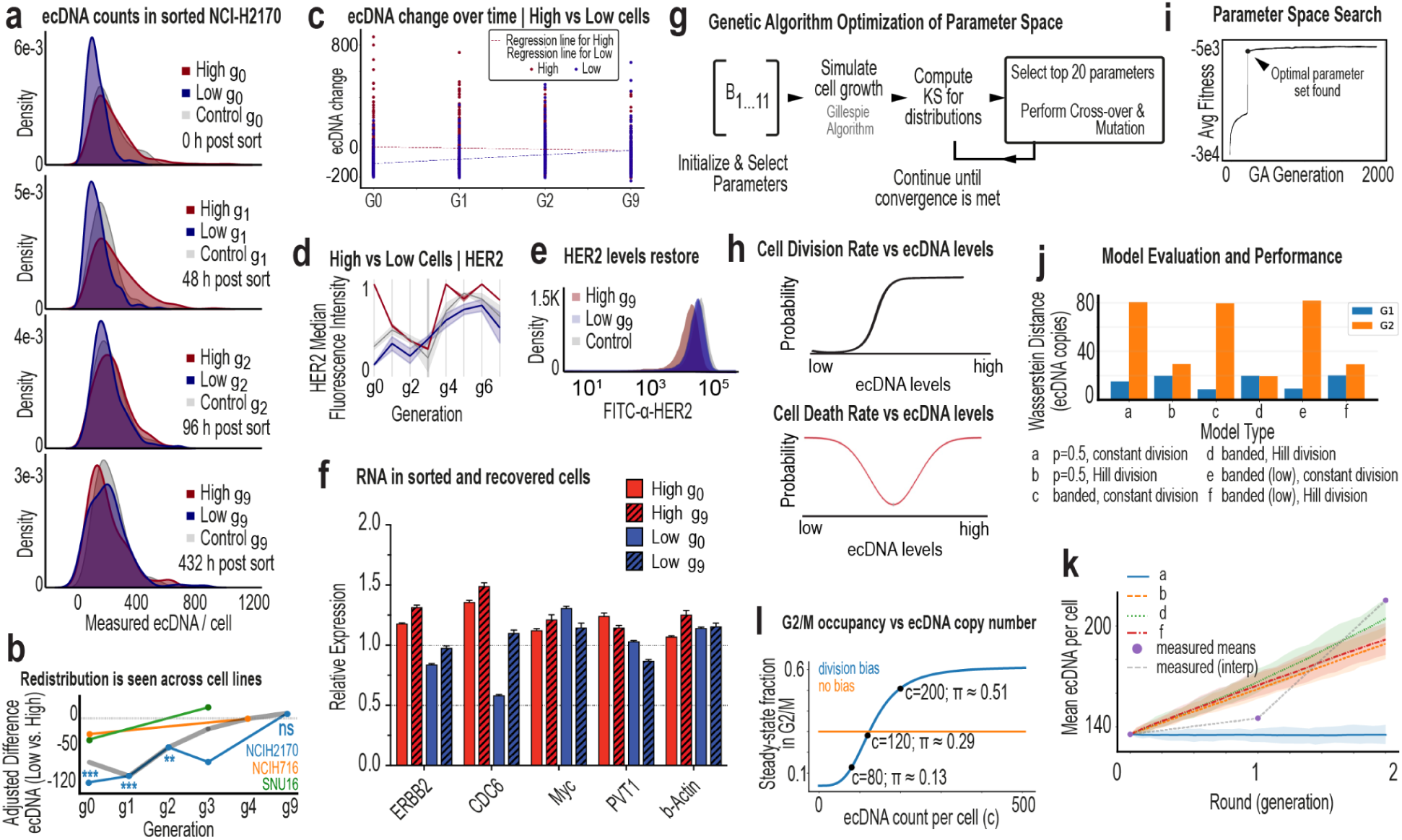
Longitudinal imaging and modeling consistent with biased inheritance and division drive rapid restoration in ecDNA variation post cell sorting. a, Distributions of ecDNA^high^ and ecDNA^low^ cells at each generation post sort. b, Longitudinal imaging plots showing rebound in ecDNA variation within several generations post sort in multiple ecDNA-bearing cell lines. c, Rate of change in ecDNA levels over time for high versus low cells, assessed via regression analysis. d, HER2 protein levels measured by flow cytometry in the first six generations post cell sorting. Shaded regions indicate 95% CI. e, Distributions of HER2 protein levels, measured by flow cytometry, at nine generations post sort (single biological replicate). f, RNA levels were quantified by digital qPCR in sorted cells at generation 0 and after nine generations of culture. Data from sorted cells represent two biological replicates with three technical replicates per condition. Data from passaged cells represent one biological replicate with three technical replicates. g, Overview of the genetic algorithm (GA) framework used to optimize parameters within a Gillespie-based stochastic simulation of ecDNA inheritance and cell population dynamics. h, Functional forms describing cell division rate (top) and death rate (bottom) as a function of ecDNA copy number per cell. Integration of optimized parameters into the Gillespie simulation framework to model stochastic cell division, death, and ecDNA segregation across generations. i, Computed fitness gains along GA-based generations in search of effective parameters for the Gillespie algorithm. Optimized parameters indicate that both inheritance bias (∼0.6 skew toward one daughter cell) and ecDNA-dependent division bias (high-ecDNA cells dividing approximately 60% faster) are required to restore baseline variation after sorting. j, Model comparison across tested architectures. Wasserstein distance between simulated and experimental ecDNA distributions at generations 1 (G1) and 2 (G2) after sorting. Models incorporating both biased inheritance and biased division best match experimental recovery. k, Comparison of simulated and experimentally measured mean ecDNA copy number per generation. Models closest to experimental trajectories demonstrate highest predictive performance. l, Steady-state fraction of cells in G2/M as a function of ecDNA level as predicted by the division bias model.

