## Supplementary Information for "Extrachromosomal DNA Gives Cancer a New Evolutionary Pathway"

#### Keywords

#### Supplementary Information

### 1. Experimental Methods

#### 1.1 Flow Cytometry and Cell Cycle Distribution Analysis

For labelling actively dividing cells in S phase, cultured NCI-H2170 or SNU-16 or NCI-NCI-H716 cells were pulsed with 10 μM EdU (Santa Cruz Biotechnology) for 1 hour at 37°C. For NCI-H2170, cells were then trypsinized with 0.25% trypsin, harvested, strained through a 50 μm Celltrics filter (Sysmex) and centrifuged for 5 minutes at 2650 rcf. For SNU-16 or NCI-H716, cells were strained through the filter and centrifuged for 5 minutes at 2350 rcf. Cells were then washed in 1x phosphate buffered saline (PBS) then fixed in 4% paraformaldehyde (Sigma-Aldrich Chemistry) diluted in 1x PBS for 15 minutes at room temperature. 1% BSA-PBS was added to the fixed cells, which were then centrifuged at 2650 rcf for 10 minutes. Cells were then resuspended in 1% BSA-PBS and stored at 4°C. All the following centrifugation steps were carried out at 2650 rcf for 10 minutes.

For cell permeabilization and EdU detection, fixed cells were permeabilized using 0.5% Triton X-100 diluted in 1% BSA-PBS for 15 minutes at room temperature, then centrifuged to remove the supernatant. Cells were then incubated with 1 μM Alexa 647-azide for NCI-H2170 cells or 488-azide for SNU-16 and NCI-H716 cells (Life Technology), 1 mM $CuSO_4$ and 100 mM ascorbic acid (prepared fresh) in 1x PBS for 30 minutes in the dark at room temperature. Then, 1% BSA-PBS with 0.5% Triton X-100 was added, and samples were strained through a 50 μm Celltrics filter and centrifuged to remove the supernatant. For primary antibody staining, NCI-H2170 cells were resuspended in 90 μl of FACS buffer (2% FBS in 1x PBS) and 10 μL of the HER2-FITC antibody (Thermo Fisher Scientific, Cat# BMS120FI) at 1:10 dilution, and incubated for 30 minutes in the dark at room temperature. For SNU-16 and NCI-H716 cells, samples were incubated in FGFR2 antibody (CST, Cat# 23328S) diluted in 1x PBS at 1:100 dilution for 30 minutes in the dark at 4 degrees. After 30 minutes, 1% BSA-PBS with 0.5% Triton X-100 was added, and samples were centrifuged to remove the supernatant. For SNU-16 and NCI-H716 cells, samples were then incubated in a donkey anti-rabbit secondary antibody conjugated to Alexa-fluor 647 (Life Technology) diluted in 1x PBS at 1:1000 for 30 minutes in the dark at 4 degrees. After 30 minutes, 1% BSA-PBS with 0.5% Triton X-100 was added, and samples were centrifuged to remove the supernatant. Finally, for analyzing DNA content using DAPI staining, samples were resuspended in 1% BSA-PBS with 0.5% Triton X-100, 1 μg/mL DAPI (Sigma-Aldrich Chemistry) and 100 μg/mL RNAse A (Sigma-Aldrich Chemistry). Samples were incubated overnight at 4°C in the dark and run the next day on Attune NxT flow cytometer (Thermo Fisher Scientific). Data was analyzed using FCS Express 7 Research (De Novo Software).

For gating, FS-area versus SS-area was used to gate cells, DAPI area versus DAPI height was used to gate singlets. The positive/negative gates for EdU and HER2-FITC or FGFR2 staining were gated on an unstained negative control sample, which was incubated with DAPI only to distinguish background from positive staining. For both HER2 or FGFR2, the low gate was determined as the bottom ~ 10% positive signal above background, while the high gate was determined as the top ~ 10% positive signal above background. The HER2+ or FGFR2 total gate was gated from the singlets gate. A representative example showing the gating strategy is illustrated in **Supplementary Fig.1**.

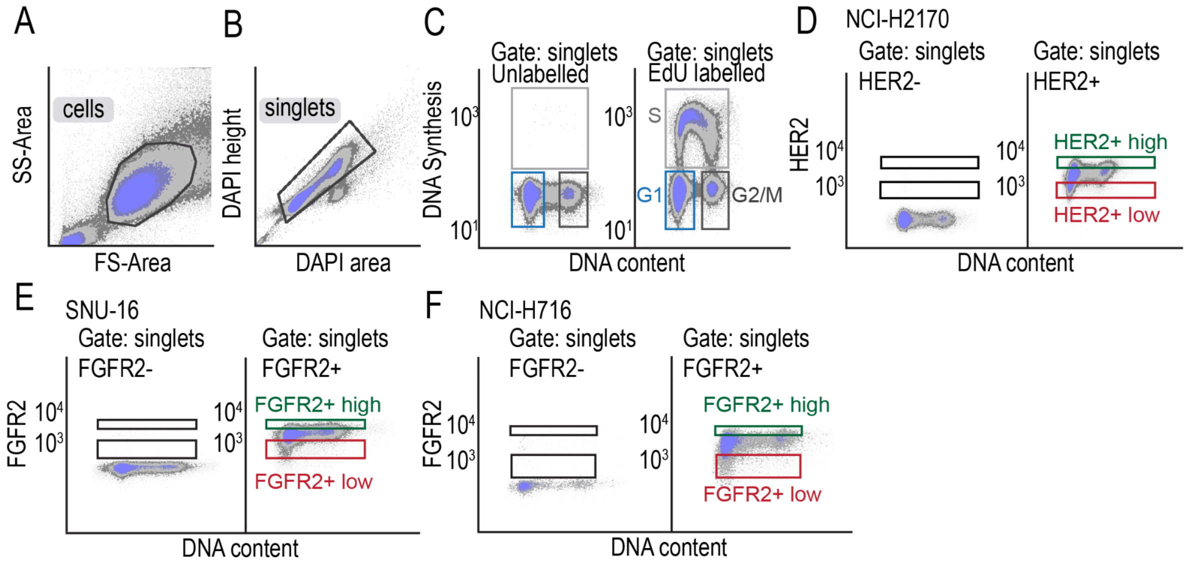

**Supplementary Fig.1: Gating scheme for flow cytometry. A**. Representative example of NCI-H2170 cells. FS-area versus SS-area is used to gate on cells and exclude any debris. **B**. Representative example of NCI-H2170 cells. DAPI area versus DAPI height is used to gate on single cells and exclude doublets. **C**. Representative example of NCI-H2170 cells. G1, S and G2/M cell cycle phases are determined based on EdU (for DNA synthesis) versus DAPI (for DNA content) staining. An unlabelled sample (left) is used to determine the gates in the EdU labelled sample (right). EdU positive cells are actively undergoing DNA replication in S phase. Both G1 and G2/M cells are EdU negative, G1 cells have 2C DNA content while G2/M cells have 4C DNA content. The y-axis values are arbitrary units on a biexponential scale. **D**. Representative gating strategy of three independent biological replicates in NCI-H2170 cells. HER2 is detected using anti-HER2-FITC antibody. HER2 negative sample (left) is used to determine background staining versus HER2 positive staining. Thresholds for high versus low HER2 positive cells (right) are determined as the top or bottom 10-15% of positive HER2 antibody staining, respectively. The y-axis values are arbitrary units on a biexponential scale. **E-F**. Representative gating strategy of three independent biological replicates in SNU-16 (**E**) or NCI-H716 (**F**) cells. FGFR2 is detected using anti-FGFR2 antibody and Alexa-fluor 647 secondary antibody. FGFR2 negative sample (left) is used to determine background staining versus FGFR2 positive staining. Thresholds for high versus low FGFR2 positive cells (right) are determined as the top or bottom 10-15% of positive FGFR2 antibody staining, respectively. The y-axis values are arbitrary units on a biexponential scale.

Doubling time for NCI-H2170 cells and cell cycle lengths measurements:
**Cell cycle phase length (hours) =** Cell cycle phase distribution (%) * Doubling time (DT) (hours) / 100

#### 1.2 Experimental Details for Iterative Indirect Immunofluorescence Imaging (4i)

4i was performed to analyze 29 proteins across 12 iterative staining and elution cycles[1]. NCI-H2170, H716, SNU16 cells were plated on a glass bottomed 96-well plate. Fibronectin (Sigma #F1141, 1ug/cm2) was used to increase cell adhesion. For pre-staining steps, cells were fixed by 4% fixing solution (formaldehyde solution 16%, Thermo 28908) for 30 min at room temperature, followed by being permeabilized by 0.1% Triton X-100 solution for 15 min at room temperature. Before the first round of antibody labeling, Hoechst staining (Hoechst 33258, 1:2500) was performed to confirm that cells were well distributed and suitable to continue 4i. For each round of antibody labeling, the blocking solution was used to incubate cells for 1h at room temperature. For every 1 ml of the blocking solution, it requires 14.6mg maleimide (Sigma #129585) and 5.35mg $NH_4Cl$ (Sigma #A9434). After washing and rinsing with PBS, the primary antibody was incubated with cells overnight at 4℃. After washing and rinsing with PBS, the secondary antibody and Hoechst stain were incubated with cells at room temperature for 1h. After washing and rinsing with PBS, the imaging buffer (700mM N-acetyl-cysteine (NAC, Sigma #A7250) $H_2O$ solution with pH=7.4) was added to get cells ready for imaging.

Fluorescence images were captured after each staining cycle using a high-resolution fluorescence microscope. Overview images (10x magnification) were used to identify regions of interest, while detailed imaging was performed at 100x magnification with tiled regions stitched using automated software. Signal intensity and antibody specificity were confirmed through positive and negative controls, ensuring high-quality data.

After each round of imaging, cells were washed 3 times with $H_2O$ and then antibodies were eluted off with the elution buffer. The elution buffer is made of 0.5M L-Glycine (Sigma #50046), 3M Urea (Sigma #U4883), 3M Guanidine chloride-GC (Invitrogen #15502-016), 70mM TCEP-HCL (Sigma #646547) with pH finally adjusted to 2.5. All wash and incubation steps were performed on a shaker. All PBS washes and rinses were performed in 200μl volume.

After completing all 12 cycles, samples were stored in PBS with 0.02% azide at 4°C for long-term preservation.

##### 1.2.1 Antibody information

**Supplementary Table 1. Primary Antibody Information**

| Iteration | Protein Target | Host Species | Dilution | Vendor | Catalog Number |
|---|---|---|---|---|---|
| 1st | c-Myc | rabbit | 1:500 | Cell Signaling Technology | 5605 |
| 1st | BRD4 | mouse | 1:1000 | Cell Signaling Technology | 63759S |
| 1st | CDK2 | goat | 1:500 | R&D Systems | AF4654 |
| 2nd | HER2 | rabbit | 1:200 | Cell Signaling Technology | 29D8 |
| 2nd | Cyclin D1 | mouse | 1:100 | Santa Cruz | sc-20044 |
| 2nd | EGFR | goat | 1:200 | R&D Systems | AF231 |
| 3rd | FGFR2 | rabbit | 1:400 | Cell Signaling Technology | 23328S |
| 3rd | CDC6 | mouse | 1:100 | Santa-Cruz | sc-9964 |
| 3rd | EZH2 | goat | 1:200 | R&D Systems | AF4767-SP |
| 4th | phospho-myc (S62) | rabbit | 1:200 | Cell Signaling Technology | 13748S |
| 4th | FOXO1 | mouse | 1:100 | Cell Signaling Technology | 14952 |
| 4th | HER2 | goat | 1:100 | R&D Systems | AD1129 |
| 5th | Phospho-myc (T58) | rabbit | 1:200 | Cell Signaling Technology | 46650S |
| 5th | Cyclin A | mouse | 1:50 | Santa-Cruz | sc-271682 |
| 5th | Cyclin B1 | goat | 1:100 | R&D Systems | AF6000 |
| 6th | FOXO3a | rabbit | 1:200 | Cell Signaling Technology | 2497 |
| 6th | AKT | mouse | 1:100 | Cell Signaling Technology | 2920 |
| 6th | ZEB1 | goat | 1:100 | abcam | ab81972 |
| 7th | phospho-Rb (S807/811) | rabbit | 1:1000 | Cell Signaling Technology | 8516 |
| 7th | Rb | mouse | 1:500 | Cell Signaling Technology | 9309 |
| 7th | p21 | goat | 1:200 | R&D Systems | AF1047 |
| 8th | CDT1 | rabbit | 1:200 | Cell Signaling Technology | 8064 |
| 8th | MDM2 | mouse | 1:200 | abcam | ab16895 |
| 9th | SKP2 | rabbit | 1:800 | Cell Signaling Technology | 2652 |
| 9th | m-TOR | mouse | 1:100 | Cell Signaling Technology | 4517 |
| 10th | TGFbeta | rabbit | 1:500 | abcam | ab215715 |
| 10th | E2F1 | mouse | 1:100 | Santa Cruz | sc-251 |

| | | | | | |
|---|---|---|---|---|---|
| 11th | c-FOS | rabbit | 1:200 | Cell Signaling Technology | 2250 |
| 11th | CDH1 | mouse | 1:100 | Santa Cruz | sc-56312 |
| 12th | EZH2 | rabbit | 1:200 | Cell Signaling Technology | 5246T |
| 12th | cPARP | mouse | 1:400 | Cell Signaling Technology | 32563 |

**Supplementary Table 2. Secondary Conjugated Antibody Information**

| Name | Host Species | Dilution | Vendor | Catalog Number | Wavelength (nm) |
|---|---|---|---|---|---|
| anti-Mouse | Donkey | 1:500 | Invitrogen | A32773 | 555 |
| anti-Rabbit | Donkey | 1:500 | Invitrogen | A32790 | 488 |
| anti-Goat | Donkey | 1:500 | Invitrogen | A21447 | 647 |

#### 1.3 10X Multiome Single-Cell Sequencing (scRNA-seq and scATAC-seq)

To characterize the transcriptional and chromatin accessibility landscapes of NCI-H2170 cells, we performed multiomics single-cell sequencing using the Chromium Single Cell Multiome ATAC + Gene Expression platform (10X Genomics). Experiments were conducted on cells prior to their third passage to ensure the analysis captured the early cellular states with minimal culture-induced artifacts.

Single-cell suspensions were prepared following the 10X Genomics protocol, ensuring a high viability (>85%) for optimal cell recovery. Cell concentration and quality were assessed using a Countess II Automated Cell Counter (ThermoFisher). Approximately 10,000 cells per sample were loaded into the Chromium Controller to partition individual cells into Gel Bead-In Emulsions (GEMs), enabling parallel profiling of RNA transcripts and chromatin accessibility within the same cells.

Library preparation was carried out according to the manufacturer's guidelines, including reverse transcription for gene expression, transposition for chromatin accessibility, and amplification of both cDNA and ATAC libraries. Libraries were quantified using a Qubit dsDNA High Sensitivity Assay (ThermoFisher) and analyzed for fragment size distribution using an Agilent 4200 TapeStation. Sequencing was performed on an Illumina NovaSeq 6000 platform with paired-end reads to ensure high-resolution data.

These results provide a comprehensive view of the transcriptional and epigenetic heterogeneity in NCI-H2170 cells, offering insights into how molecular programs are coordinated within single cells.

#### 1.4 Nanopore Long Read Sequencing with Adaptive Sampling

Nanopore long-read sequencing with adaptive sampling was employed to profile specific genomic regions of interest in NCI-H2170 cells, focusing on chromosomes 17 and 8. Adaptive sampling, a feature unique to nanopore sequencing, was used to enrich DNA fragments originating from these chromosomes, enabling targeted sequencing without the need for physical enrichment steps.

Two sequencing approaches were tested to evaluate performance: a ligation-based protocol and a rapid sequencing protocol, both following the standard workflows recommended by Oxford Nanopore Technologies (ONT). For the ligation protocol, high molecular weight DNA was extracted, quantified, and processed to ensure optimal read lengths. DNA libraries were prepared using ONT's ligation sequencing kit, and sequencing was conducted on a MinION or PromethION platform. The rapid protocol involved the use of ONT's rapid sequencing kit, streamlining library preparation while maintaining sufficient read quality and throughput.

Sequencing runs were performed using adaptive sampling to selectively sequence reads from chromosomes 17 and 8 by dynamically rejecting reads originating from other chromosomes. This real-time targeting was managed through ONT's adaptive sampling algorithms integrated into MinKNOW software. Metrics such as enrichment efficiency, coverage depth, and read length distribution were compared across the two protocols to assess the suitability of each for future experiments.

This approach provided a detailed view of the genomic architecture of chromosomes 17 and 8, enabling high-resolution analysis of regions of interest while leveraging the versatility of adaptive sampling for targeted sequencing.

#### 1.5 G-band Karyotyping

Cytogenetic analysis was conducted on 25 G-banded metaphase spreads from the human cancer cell line NCI-H2170. In 23 spreads, the chromosome count ranged from 62 to 68, consistent with a near-triploid karyotype. Two spreads exhibited near-hexaploid chromosome counts of 126 and 129. Most spreads displayed a sex chromosome complement consisting of two X chromosomes and one Y chromosome. However, two spreads showed one apparently normal X chromosome and an X chromosome with additional chromatin of unknown origin attached to its p arm, while one spread contained a single X chromosome.
All spreads demonstrated multiple chromosomal aberrations, with minor variations between individual spreads. Multiple copies of certain chromosomes, particularly chromosomes 7 and 20, were frequently observed. Common structural aberrations included two variants of a derivative chromosome 1 with a p arm deletion replaced by chromatin of unknown origin; an isochromosome of the chromosome 13 q arm; an additional chromatin segment of unknown origin attached to the p arm of one copy of chromosome 13; and two copies of chromosome 14 with an interstitial duplication of the distal 14q arm.

Additionally, each spread displayed approximately 8 to 100 double minutes and two to seven marker chromosomes. Marker chromosomes, defined as structurally abnormal chromosomes that cannot be unambiguously identified by conventional banding techniques, were a consistent feature across spreads. These findings highlight the extensive genomic instability characteristic of NCI-H2170 cells.

#### 1.6 Bioskryb ResolveOME Single-Cell Sequencing (scRNA-seq + scDNA-seq)

To sequence cells with different amounts of protein expression and ecDNA abundance, we collaborated with BioSkryb Genomics, Inc. to collect different subpopulations of cells from three different cells (NCI-H2170, SNU16, and KATO III) using FACS and then performed sequencing.

Trypsinization was used to detach and collect NCI-H2170 and KATO III cell lines. Only the cells in suspension for the SNU16 cell line were collected. The cells from each line were resuspended in 10 mL of FACS buffer (2% FBS in a solution of 1X PBS). A 70 micron cell strainer was used to strain the cells and remove large clumps. The cells were then counted and resuspended in a smaller volume of FACS buffer at a concentration of 1.33x10^7 cells/mL. Control samples, consisting of unstained and live/dead stained cells, were prepared prior to adding antibodies and used to establish the baseline fluorescence and to gate on viable cells. The NCI-H2170 cells were stained with the antibody HER2-FITC (Thermo Fisher Scientific, Cat# BMS120FI). The KATO III and Snu16 cell lines were each stained with the antibody FGFR2-FITC (FabGennix, Cat# FGFR2-FITC). All three cell lines were stained at a concentration of 1:50 (1uL of antibody added for every 49uL of cells).

After cell preparation, single cells were isolated via FACS and plated based on HER2 intensity into microtiter plates with one cell for each well. Paired DNA and RNA libraries from single cells were prepared using ResolveOME Whole Genome and Whole Transcriptome Single Cell Core Kit following manufacturer's instructions. Briefly, cytosolic lysis was performed, and the first strand cDNA was synthesized from cytosolic mRNA via reverse transcription. Following nuclear lysis, the same cell underwent whole genome amplification via Primary Template directed Amplification (PTA). Then the transcriptome cDNA was purified, amplified, and prepared into individual NGS libraries. Illumina NextSeq 2000 platform was used for paired end 50 BPs sequencing of both the DNA arm, targeting 2×10^6 reads, and the RNA arm, targeting 2×10^5 reads per single cell library.

#### 1.7 Metaphase Sample Preparation and Fluorescence In Situ Hybridization (FISH)

Generating condensed chromatin during metaphase allows for optimal imaging of ecDNA. Before karyotyping, cells underwent a four-stage preparation: arrest at metaphase, incubation in a hypotonic solution, cell fixation, and staining. Samples were prepared using cells cultured by the Brunk Lab.

Cells were arrested at metaphase by treating with colcemid at 0.1 µg/mL (10 µg/mL Colcemid Solution, FUJIFILM Irvine Scientific) in cell culture media when cells reached ~70% confluency. Colcemid arrests cellular division during mitosis by binding to tubulin, preventing spindle formation and cytokinesis. Cells were incubated with colcemid for 12–20 hours before being harvested following standard cell culture procedures. For adherent or semi-adherent cells, trypsinization was used to detach cells, and this was quenched with a mixture of cold colcemid-spiked media and PBS wash to maximize yield. Cells were resuspended in 1 mL of 1x PBS by pipetting and transferred to 1.5 mL microcentrifuge tubes for centrifugation at 5000 rpm for 2 minutes.

The cells were incubated with 600 µL of pre-warmed 37°C 0.075M KCl (Gibco), added dropwise with gentle agitation to resuspend cells. After 15 minutes at 37°C, the cells became swollen and fragile due to osmotic pressure, making them ready for fixation. Freshly prepared modified Carnoy's fixative (3:1 methanol:glacial acetic acid) was added dropwise to each sample. Tubes were immediately centrifuged at 5000 rpm for 2 minutes. After leaving ~150 µL of supernatant, pellets were gently agitated and resuspended. Another 600 µL of fixative was added dropwise, followed by agitation and centrifugation for 2 minutes at 5000 rpm. This fixation step was repeated three times, with the final addition of fixative adjusted to achieve ~6 million cells/mL (0–1 mL), ensuring optimal density for single-cell imaging.

Microscope slides were prepared using Superfrost™ Microscope Slides (Fisherbrand, Cat. No: 12550123), which are uncharged. Slides were humidified using water vapor immediately before a drop (10 µL) of the prepared cell suspension was dropped from a height of ~60–70 cm onto the slide. The slides were left to air dry for an hour. Dried slides were equilibrated in 2x saline sodium citrate (SSC) (Ultrapure™ 20X SSC buffer, Invitrogen, Cat. No: 15557-036) and dehydrated through an ascending ethanol series (70%, 85%, 100%) for 2 minutes each. Slides were stored at 37°C for 16–20 hours in a slide moat before staining.

Slides were then washed in 0.4x SSC and 2x SSC + 0.05% Tween20 for 2 minutes each, followed by a final dip in 2x SSC. Fluorescent DNA probes (Empire Genomics) were applied (5 µL), and SlowFade™ Diamond Antifade Mountant with DAPI (Invitrogen, Cat. No: S36964) was added to the center. Microscope cover glass slips (Fisherbrand, Cat. No: 12541036) were applied and sealed with nail polish.

All images were captured using an Echo Revolve Microscope (Echo, San Diego, CA) at 60x magnification. Images were taken from the same slide or occasionally from two slides prepared from the same metaphase spread to ensure consistency in experimental analysis. While uncharged slides were used for all metaphase spreads, potential differences in ecDNA adherence between charged and uncharged slides were not specifically tested.

#### 1.8 Fluorescence-Activated Cell Sorting (FACS)

##### 1.8.1 Preparation of Metaphase Spreads for ecDNA Quantification in Sorted Subpopulations

To measure the abundance of ecDNA associated with differential protein expression, we combined FACS sorting with FISH imaging of metaphase-arrested cells across three cell lines: NCI-H2170, NCI-H716, and SNU16. Sixteen hours before FACS sorting, cells were treated with colcemid (10 µg/mL stock diluted to 0.1 µg/mL final concentration) to arrest them in metaphase.

Cells were harvested using trypsinization, resuspended in FACS buffer (PBS + 2% FBS), and stained with the following antibodies:

For NCI-H2170: HER2-FITC (Thermo Fisher Scientific, Cat# BMS120FI, 1:40 dilution, 30 min incubation).

For NCI-H716 and SNU16: FGFR2 primary antibody (Cell Signaling Technology, Cat# 23328S, 1:400 dilution, 30 min), followed by Alexa Fluor 647 secondary antibody (Cell Signaling Technology, Cat# 4414S, 1:1000 dilution, 60 min).

After each staining step, cells were washed twice in the FACS buffer.

Live/dead discrimination was performed using Annexin V and Cytox Blue staining. Controls (unstained and live/dead stained) were used to establish gating parameters. Cells were sorted into three groups: the top 10% (high expression), bottom 10% (low expression), and a live/dead control population gated without regard to protein expression. Post-sort analysis confirmed minimal overlap between high- and low-expression groups (**Supplementary Fig. 2**).

Sorted cells were processed for metaphase spread preparation. Cells were counted, resuspended in 1 mL PBS, and centrifuged at 5000 rpm for 2 minutes. Pellets were resuspended in 600 µL of pre-warmed 0.075 M KCl, incubated for 15 minutes at 37°C to induce swelling, and fixed using dropwise addition of Carnoy's fixative (3:1 methanol:acetic acid). Fixation was repeated three times, and final cell concentrations were adjusted to approximately 1 million cells/mL for optimal spread density.

Metaphase spreads were prepared by dropping 10 µL of the suspension from ~60–70 cm height onto uncharged Superfrost™ slides (Fisherbrand, Cat# 12550123) pre-humidified with water vapor. Slides were air-dried for 1 hour, then equilibrated in 2x SSC buffer and dehydrated through a graded ethanol series (70%, 85%, 100%). Slides were stored overnight at 37°C before hybridization.

For FISH, slides were washed sequentially in 0.4x SSC, 2x SSC + 0.05% Tween20, and 2x SSC. Fluorescent DNA probes (Empire Genomics) were applied (5 µL per slide) and sealed with SlowFade™ Diamond Antifade Mountant with DAPI (Invitrogen, Cat# S36964) and coverslips (Fisherbrand, Cat# 12541036).

All images were captured at 60× magnification on an Echo Revolve Microscope (Echo, San Diego, CA). Images were acquired from the same slide or occasionally two slides prepared from the same metaphase harvest to ensure experimental consistency. Only uncharged slides were used; potential differences between charged and uncharged substrates were not systematically evaluated.

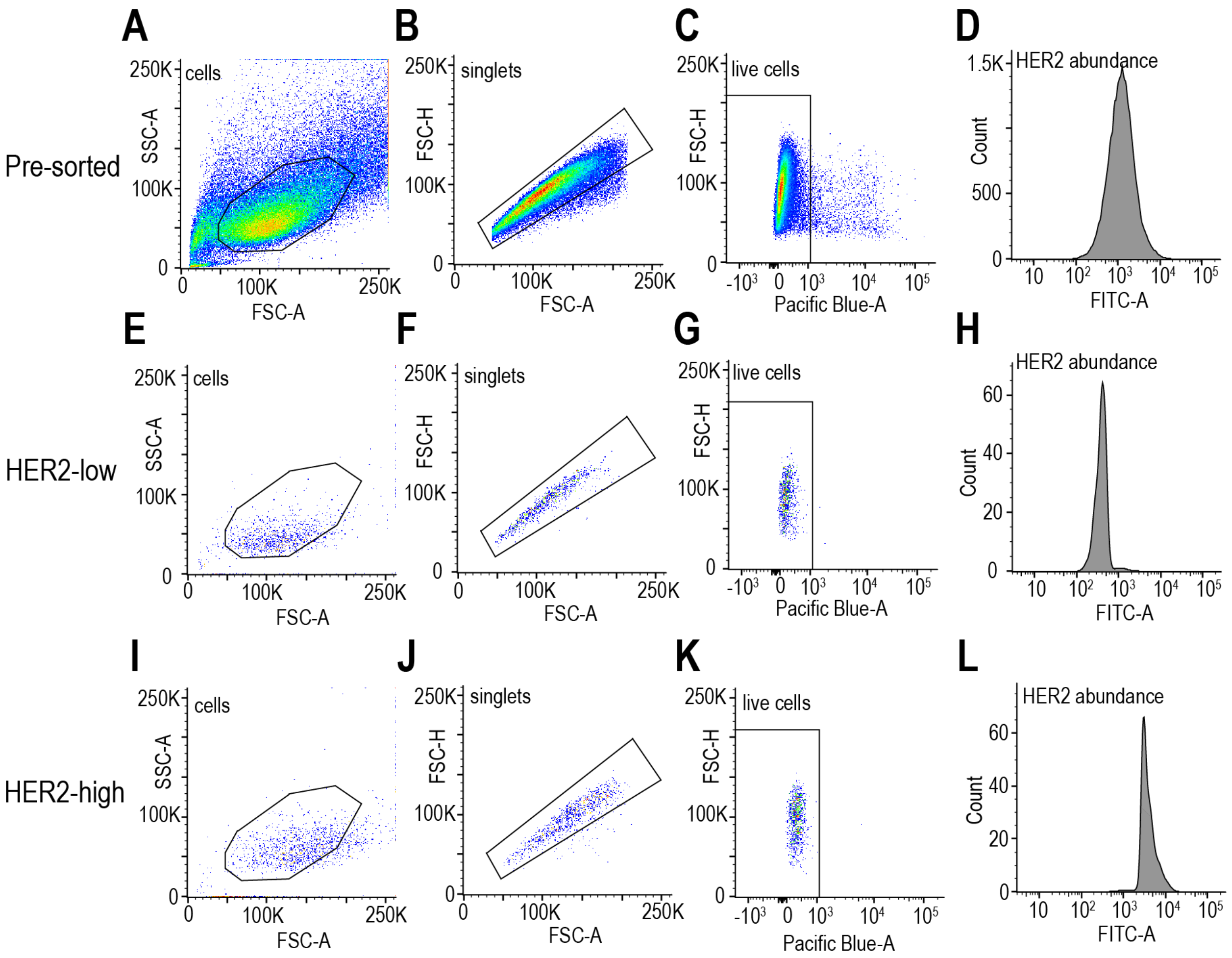

**Supplementary Fig.2: Flow cytometry gating and cell sorting strategy. A**. Pre-sorted NCI-H2170 cells. SSC-A versus FSC-A plot is used to gate on cells and exclude any debris. **B**. Pre-sorted NCI-H2170 cells. FSC-H vs FSC-A plot is used to identify single cells and remove doublets or clumps. **C**. Pre-sorted NCI-H2170 cells. FSC-H vs Pacific Blue-A plot is used to identify live cells. **D**. FITC intensity, indicating HER2 protein amount, for all pre-sorted NCI-H2170 cells. Cells with the top and bottom 10% of HER2 intensities are collected and regarded as HER2-high and HER2-low cells. **E.** SSC-A vs FSC-A plot shows the post-sorted HER2-low NCI-H2170 cells. **F.** FSC-H vs FSC-A plot shows the post-sorted HER2-low NCI-H2170 cells are single cells. **G.** FSC-H vs Pacific Blue-A plot shows the post-sorted HER2-low NCI-H2170 cells are live cells. **H.** FITC intensity for post-sorted HER2-low cells. **I.** SSC-A vs FSC-A plot shows the post-sorted HER2-high NCI-H2170 cells. **J.** FSC-H vs FSC-A plot shows the post-sorted HER2-high NCI-H2170 cells are single cells. **K.** FSC-H vs Pacific Blue-A plot shows the post-sorted HER2-high NCI-H2170 cells are live cells. **L.** FITC intensity for post-sorted HER2-high cells.

#### 1.8.2 FACS-Based Growth and Recovery

To monitor changes in ecDNA abundance and protein expression following sorting, FACS was performed on live cells, and sorted populations were maintained in culture. To ensure viability, antibody staining prior to sorting targeted cell surface proteins specific to each cell line.

Immediately after FACS, cells from each subpopulation and control group were resuspended in antibiotic-supplemented media (1% Penicillin-Streptomycin, 10,000 U/mL, Gibco; 0.1% Gentamicin, 50 mg/mL) to prevent bacterial contamination from the sorters.

Sorted cells were plated into appropriate vessels based on growth characteristics: adherent NCI-H2170 cells were plated into 6-well dishes, while suspension NCI-H716 and SNU16 cells were plated into T25 flasks.

Cells were maintained in antibiotic-containing media throughout recovery and expansion. Cultures were passaged into new plates or flasks upon reaching 70–80% confluency.

##### 1.8.3 FACS-Based Redistribution Kinetics

To measure the redistribution kinetics of sorted populations, cells were collected every 48 hours for two weeks following the initial FACS sort. At each time point, ecDNA abundance and protein expression levels were measured and recorded.

For each subpopulation and control group, a portion of cells (one T25 flask or one well of a 6-well dish) was collected for both FISH analysis and flow cytometry.

*FISH Sample Preparation:*

To collect cells for metaphase FISH imaging, cultures were treated with colcemid (0.1 μg/mL) 12–20 hours before harvest to arrest cells in metaphase. Cells were then processed according to the Metaphase Sample Preparation and Fluorescence In Situ Hybridization (FISH) Imaging protocol. Prepared metaphase samples were stored at –20°C until slides were made and imaged.

*Flow Cytometry Sample Preparation:*

To collect cells for flow cytometry, samples were harvested (using trypsin if necessary), resuspended in 200 μL of 4% paraformaldehyde (PFA), and incubated for 15 minutes on ice. After fixation, cells were washed in an excess of 1X PBS and stored at 4°C for up to one week prior to antibody staining and flow cytometry.

For NCI-H2170 cells, staining was performed with HER2-FITC antibody (Thermo Fisher Scientific, Cat# BMS120FI) at 1:40 dilution for 30 minutes. Cells were then washed twice with 1X PBS and resuspended in 500 μL PBS for analysis on a Thermo Fisher Attune NxT flow cytometer. Gating strategies for early time points (g1, g2, and g3) are provided in **Supplementary Fig. 3**.

For NCI-H716 and SNU16 cells, samples were first stained with an FGFR2 primary antibody (Cell Signaling Technology, Cat# 23328S) at 1:400 dilution for 30 minutes at 4°C, washed twice, then stained with an Alexa Fluor 647-conjugated secondary antibody (Cell Signaling Technology, Cat# 4414S) at 1:1000 dilution for 60 minutes at 4°C. After two additional PBS washes, cells were resuspended in 500 μL PBS for flow cytometry.

*Data Analysis:*

Flow cytometry data were analyzed using FlowJo software (v10.10.0) to assess population purity and quantify fluorescence intensity distributions over time.

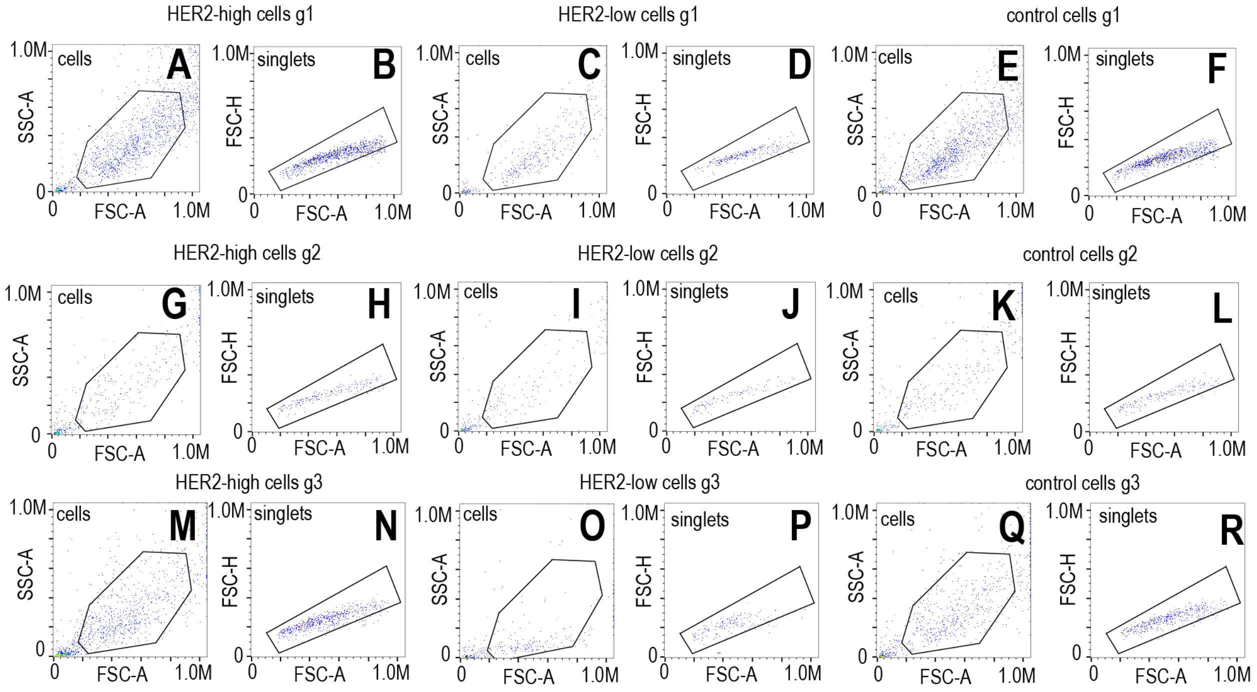

**Supplementary Fig.3: Flow cytometry gating strategy for post-sorted NCI-H2170 cells at g1, g2, and g3 time points. A**. Post-sorted NCI-H2170 HER2-high cells at g1. SSC-A vs FSC-A plot is used to gate on cells and exclude any debris. **B**. Post-sorted NCI-H2170 HER2-high cells at g1. FSC-H vs FSC-A plot is used to identify single cells and remove doublets or clumps. **C**. SSC-A vs FSC-A plot of post-sorted NCI-H2170 HER2-low cells at g1. **D**. FSC-H vs FSC-A plot of post-sorted NCI-H2170 HER2-low cells at g1. **E.** SSC-A vs FSC-A plot of post-sorted NCI-H2170 control cells at g1. **F.** FSC-H vs FSC-A plot of post-sorted NCI-H2170 control cells at g1. **G**. SSC-A vs FSC-A plot of post-sorted NCI-H2170 HER2-high cells at g2. **H**. FSC-H vs FSC-A plot of post-sorted NCI-H2170 HER2-high cells at g2. **I**. SSC-A vs FSC-A plot of post-sorted NCI-H2170 HER2-low cells at g2. **J**. FSC-H vs FSC-A plot of post-sorted NCI-H2170 HER2-low cells at g2. **K.** SSC-A vs FSC-A plot of post-sorted NCI-H2170 control cells at g2. **L.** FSC-H vs FSC-A plot of post-sorted NCI-H2170 control cells at g2. **M**. SSC-A vs FSC-A plot of post-sorted NCI-H2170 HER2-high cells at g3. **N**. FSC-H vs FSC-A plot of post-sorted NCI-H2170 HER2-high cells at g3. **O**. SSC-A vs FSC-A plot of post-sorted NCI-H2170 HER2-low cells at g3. **P**. FSC-H vs FSC-A plot of post-sorted NCI-H2170 HER2-low cells at g3. **Q.** SSC-A vs FSC-A plot of post-sorted NCI-H2170 control cells at g3. **R.** FSC-H vs FSC-A plot of post-sorted NCI-H2170 control cells at g3.

### 2. Computational Methods

#### 2.1 Analysis of DNA composition Differences

The percentages of **50%** and **180%** represent the fractional increase in genome size when moving from a normal **diploid** genome (46 chromosomes) to either a **near-triploid** genome (69 chromosomes) or a **near-hexaploid** genome (129 chromosomes). Here's the step-by-step breakdown:

##### 2.1.1. Define the Baseline Diploid Genome Size

The diploid human genome contains two sets of 23 chromosomes, for a total of 46 chromosomes, with an estimated total size of approximately 6.4 billion base pairs. This corresponds to 2 × the haploid genome size, where the haploid genome (23 chromosomes) contains about 3.2 billion base pairs.

Diploid Genome Size = 2 × Haploid Genome Size = 6.4 billion base pairs

##### 2.1.2. Calculate the Genome Sizes for Each Ploidy

Genome size scales linearly with chromosome count, so we can estimate total DNA content in higher ploidy states by comparing chromosome numbers:

Diploid (46 chromosomes):
Genome Size = 6.4 billion base pairs

Near-Triploid (69 chromosomes):
Genome Size = (69 / 46) × 6.4B = ~9.6 billion base pairs

Near-Hexaploid (129 chromosomes):
Genome Size = (129 / 46) × 6.4B = ~18.0 billion base pairs

##### 2.1.3. Calculate the Fractional Increase

To quantify how much larger these genomes are relative to diploid:

Near-Triploid:

Fractional Increase = (9.6B − 6.4B) / 6.4B = 0.5 → 50% increase

Near-Hexaploid:

Fractional Increase = (18.0B − 6.4B) / 6.4B = 1.8 → 180% increase

To put this in context, the average human chromosome contains approximately 150 million base pairs (Mb). The difference in ecDNA content between “high” and “low” cells—estimated at 540 Mb—is therefore:

(540,000,000 bp / 150,000,000 bp) × 100 = 360% of an average chromosome

##### 2.1.4 Summary

**Near-Triploid Genome: ~50% larger than diploid**

**Near-Hexaploid Genome: ~180% larger than diploid**

**ecDNA gain of 540 Mb is equivalent to adding ~3.6 average-sized chromosomes**

These calculations illustrate how DNA content increases with chromosome count and underscore the significant genomic burden imposed by elevated ploidy or high levels of ecDNA in cancer cells.

#### 2.2 Computational Analysis of 4i Data

Python (v3.7.1) was used to process the images acquired from the previous iterative indirect immunofluorescence imaging step. Image segmentation was performed using the package Cellpose[2] (v2.0.5). pyStackReg (v0.2.5) library was used to align the segmented images. Manual adjustment was made to align a problematic round if necessay. Areas of artifacts were drawn and excluded from further analysis using Napari[3] (v0.4.18). Scikit-image[4] was used to extract features from images. A very detailed image processing workflow can be found in this publication[5] and this GitHub repository (https://github.com/fjorka/4i_analysis). After feature extraction, z-scores were calculated for each of the features. And for each feature, the cells that have the top 5% and bottom 5% of signal intensity were excluded to get rid of outliers.

For HER2 protein intensity, the top 10% and bottom 10% of cells were selected as “HER2-high” and “HER2-low”. Then the z-scores of other proteins of interest, such as MYC and CDC6, were compared between the “HER2-high” and “HER2-low” sub-populations using the Wilcoxon Rank Sum Test. For the “HER2-high” and “HER2-low” subpopulations respectively, the Pearson correlations between all protein pairs were calculated using the corr() function in python. Hierarchical clustering was performed on the correlation matrix using the linkage and squareform functions from the scipy package (v.1.10.1), and a clustered triangular protein correlation heatmap was generated using the package seaborn (v.0.11.1).

For protein network analysis, a correlation network was created based on protein-protein correlations between all protein pairs using the python package networkx (v.3.1) for “HER2-high” and “HER2-low” cells respectively. Edges connecting two proteins were added if the protein pair correlation was above 0.5. Degree of centrality was computed using the degree_centrality() function in the networkx package.

Logistic regression analysis was conducted using the sklearn.linear_model module (v.1.3.2) and statsmodels.api (v.0.14.0) to classify single cells as HER2-high or HER2-low based on nuclear protein expression in NCI-H2170 cells. The model was trained on the following features:

['12_cPARP_nuc_mean', '11_CDH1_nuc_mean', '12_EZH2_nuc_mean', '01_CDK2_nuc_mean', '02_CyclinD1_nuc_mean', '02_EGFR_nuc_mean', '03_FGFR2_nuc_mean', '03_CDC6_nuc_mean', '03_EZH2_nuc_mean', '04_FOXO1_nuc_mean', '05_CyclinB1_nuc_mean', '05_cMycT58_nuc_mean', '06_AKT_nuc_mean', '10_TGFbeta_nuc_mean', '09_mTOR_nuc_mean', '07_p21_nuc_mean', '06_ZEB1_nuc_mean', '11_cFOS_nuc_mean'].

From the trained model, z-scores and p-values were extracted to quantify the statistical significance and direction of each feature's contribution to the HER2 classification. Odds ratios were calculated from the logistic regression coefficients, offering interpretable effect sizes for each protein marker.

#### 2.3 Computational Analyses of 10X Single Cell Multiomics Data

##### 2.3.1 General Analysis Details

Cell Ranger ARC (v2.0.1) was used to align raw FASTQ data against the GRCh38 genome. Downstream data processing and analysis was performed in R (v.4.2.1/v.4.3.1) language. Seurat (v4.4.0) and Signac (v1.10.0) were used to further process and analyze the 10X single-cell multiome data. A seurat object was built for the NCI-H2170 single cell multiomic data following the standard protocol provided on the Signac webpage (https://stuartlab.org/signac/articles/pbmc_multiomic) Briefly, single-cell barcodes meeting all the following criteria were kept for further analysis: ATAC read counts between 1,000 and 70,000; RNA read counts between 1,000 and 25,000; >500 genes detected in each individual cell; percentage of mitochondrial gene transcripts < 20%; nucleosome signal < 2; TSS.enrichment > 1. DoubletFinder[6] (v2.0.4) was used to identify and filter out doublets.

For single-cell RNA data, SCTransform() function in Seurat was used for data normalization and the results were saved in the "SCT" assay in the seurat object. For single-cell ATAC data, the CallPeaks() function was used for peak calling. The GeneActivity(biotypes = NULL) function from Signac was used to calculate gene activity as counts in the gene body and promoter region for each gene, and the results were saved in the "gene_activity" assay in the seurat object.

To identify genes associated with the genes amplified on ecDNA, single cells with the top (ecDNA-high) and bottom (ecDNA-low) 10% of ecDNA gene signals (such as mRNA or gene activity) were selected. FindMarkers(test.use = Wilcoxon Rank Sum Test, min.pct = 0.25, logfc.threshold = 0.10) function in Seurat was used to find differentially expressed genes (DEG) or differential gene activities between the ecDNA-high and ecDNA-low sub-populations. Genes whose chromatin accessibility were strongly correlated with ecDNA gene chromatin accessibility were plotted as hollow circular heatmap with mRNA-ATAC links between pairs of genes in the middle using the package Circlize (v.0.4.15). For each gene in the circular heatmap, the levels of CCLE/DepMap bulk-level copy number (v.22Q1), 10X scATAC pseudobulk chromatin accessibility, CCLE/DepMap bulk-level expression (v.23Q2), and 10X scRNA pseudobulk expression are shown. AggregateExpression() function was performed to calculate summed counts for the "gene_activity" and "SCT" assay.

##### 2.3.2 Inference of ecDNA counts

To infer ecDNA counts from scATAC data, we used a method adapted from the previous publications[7–9]. The R package epiAneufinder (v.1.0.2)[10] was used to calculate the GC content corrected coverages for each 100kb bin across the GRCh38 genome. The chr8 region (chr8-127300001-128900000) amplified on ecDNA was used to infer copy numbers of MYC and the dominant ecDNA species in NCI-H2170. From the DepMap bulk copy number data, the mean copy number (CN) across all genes in NCI-H2170 is around 1, which means the genes with a relative CN of 1 don't have any amplification or deletion relative to its ploidy and the exact CN of these genes are 1*NCI-H2170 ploidy= NCI-H2170 ploidy. Therefore, it is speculated that in 10X scATAC data where gene dosage contributes the most to chromatin accessibility, the same thing holds true. In each single cell, the mean normalized coverage for all bins is the coverage for chromosome regions that don't have amplification or deletion relative to its ploidy. And this region can be used as the chromatin accessibility baseline to infer ecDNA copy number. Since from G-band Karyotyping the ploidy of NCI-H2170 is roughly 4.5 (the average number of triploid and hexaploid), the exact CN for these regions is 4.5. Then if the copy number of the region “chr8-127300001-128900000” is A, the normalized coverage for this region is B, the mean normalized coverage for all bins is C, we have A/B=4.5/C. Since the values of B and C for each single cell are available, we can calculate the copy number of the bin “chr8-127300001-128900000”, which is 4.5*B/C. The ecDNA counts inferred from 10X scATAC data is shown in **Supplementary Fig.4**.

To predict ecDNA counts for single cells in this 10X sc-multiome dataset, gene expression count per million (CPM) was calculated from the raw unique molecular identifier (UMI) counts across single cells for further machine learning analysis.

##### 2.3.3 Comparison of inferred ecDNA in scATAC and scDNA with FISH

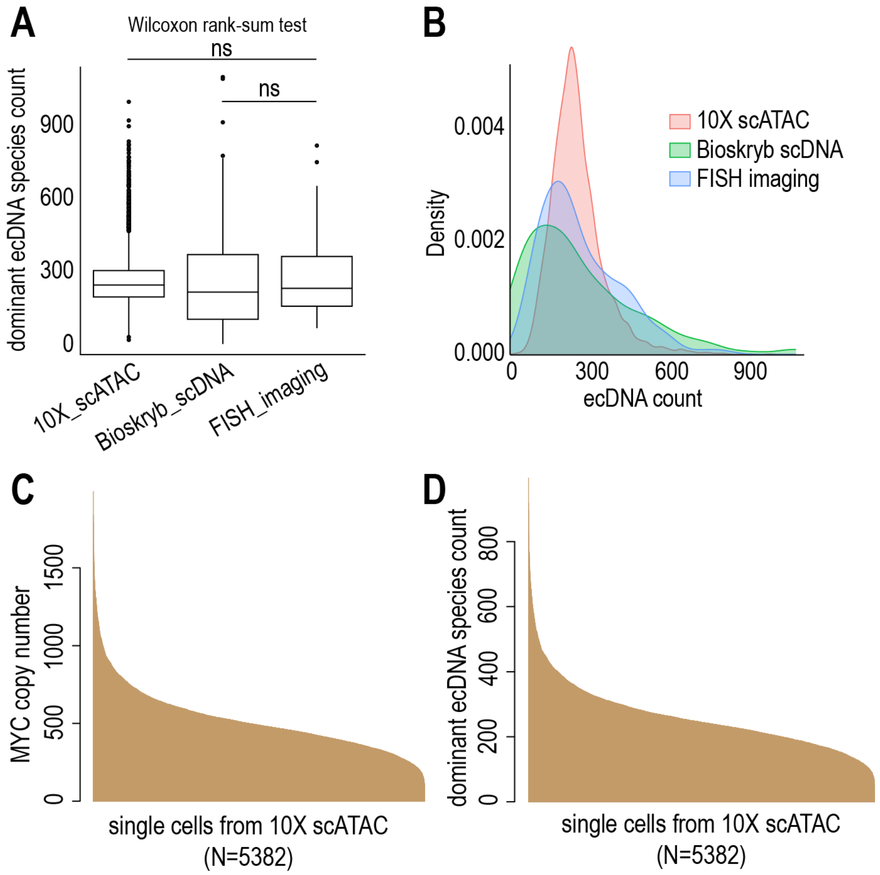

**Supplementary Fig.4: Inferred ecDNA counts from 10X scATAC data. A.** Comparison of inferred ecDNA counts between different techniques (10X scATAC-seq, Bioskryb scDNA-seq and FISH imaging) **B.** KDE plots of inferred ecDNA counts from 10X scATAC-seq data, inferred ecDNA counts from Bioskryb scDNA-seq data, ecDNA counts from FISH imaging. **C.** inferred MYC copy number barplot across single cells in 10X scATAC data **D.** inferred dominant ecDNA species count barplot across single cells in 10X scATAC data.

Although single-cell ATAC-seq (scATAC) can provide indirect signals of ecDNA abundance through chromatin accessibility, it systematically underestimates ecDNA copy number variation compared to direct DNA-based measurements like single-cell DNA-seq (scDNA) and FISH imaging. While Wilcoxon rank-sum tests show no significant difference in the median distributions across scATAC, scDNA, and FISH datasets, Levene’s tests reveal a striking compression of variance: scATAC-inferred ecDNA counts exhibit significantly reduced spread relative to FISH ($p = 5.96\times10^{-13}$), whereas scDNA-inferred counts match FISH variance much more closely ($p = 0.0061$).

This discrepancy arises because scATAC measures chromatin accessibility, not DNA abundance. Amplified ecDNAs can exist in closed or transcriptionally silenced chromatin states, rendering them invisible to Tn5-based profiling despite high copy number. Additionally, ecDNAs often contain small focal hubs of enhancer activity surrounded by inaccessible DNA, which can be missed when accessibility is integrated over broader genomic regions. Technical factors further contribute: scATAC library preparation and PCR amplification biases underrepresent high-copy repetitive regions, and limited read depth at the single-cell level masks high-variance amplification events. Finally, because Tn5 preferentially targets nucleosome-free regions, any inaccessible ecDNA fragments—including structural variants or breakpoints—remain undetected.

Together, these biological and technical limitations explain why scATAC-based ecDNA inference produces a compressed distribution and reduced dynamic range compared to scDNA and FISH. For downstream analyses, we therefore use scATAC primarily to infer regulatory associations with gene expression rather than to quantify absolute ecDNA burden. To analyze these accessibility-transcription associations, we adapted a method from Regner et al.[11] and Signac's LinkPeaks() function[12] to identify both proximal and distal ATAC peaks linked to gene expression, as described in our GitHub repository (https://github.com/Brunk-Lab/ecMultiOME).

#### 2.3.4 Analysis of Gene Expression to Accessible Peak Correlations

Transcription factor (TF) binding motifs were extracted from the database JASPAR2020 (v.0.99.10) and the motifs were quantified across single cells using the package chromVAR (v.1.20.2). Differential accessible peaks were found between cells with top and bottom 10% of ERBB2 expression or chromatin accessibility using the function FindMarker(test.use="LR"). Enriched TF motifs were discovered in the differential accessible peaks using the function FindMotifs().

Step 1: Sets of neighboring cells (default = 50) based on dimension reduction (e.g., LSI, PCA, UMAP, Harmony) were combined using the function "generate_metacells()" to generate a "metacell" object. Using metacells instead of single cells provides more robust results by accounting for the usual sparsity of ATAC and RNA data in single cells.

Step 2: To prepare for correlation analysis, genomic ranges of the seurat object were extracted using the function "extract_genomic_ranges()". Gene genomic coordinates were extracted using the function "extract_genomic_ranges()". Genomic range objects were created using the function "create_gene_granges()".

Step 3: To look for nearby ATAC peaks that regulate gene expressions, for each gene in the seurat object, ATAC peaks located within a specific distance from the gene (default = 500 Kbp) were identified using the function "find_nearby_peaks()".

Step 4: The function "calculate_gene_peak_correlations()" was used to calculate the correlation for every gene-peak pair within a 500 Kbp proximity from the metacell object.

Step 5: To calculate p-values and adjusted p-values, for any gene-peak pair with the absolute value of the computed correlation above a given threshold (default > 0.2), the null distribution of the correlations was calculated using the function "calculate_null_correlations_filtered()". This function permutes the gene expression values and calculates gene-peak correlations for the permuted values and peak counts. The empirical p-values and the adjusted p-values using Benjamini-Hochberg procedure were calculated using the function “calculate_fdr()”.

Step 6: To analyze the distal peak correlations (unrestricted to genome proximity) of specific genes, the expression of the gene of interest was from the “metacell” object, and the Pearson correlation between the gene expression and each peak was calculated.

Step 7: To visualize the distribution of the gene-peak links in the genome, we calculated the number of significant gene-peak correlations in bins of 50 Kpb throughout the genome and generated a density heatmap with each chromosome.

#### 2.4 Computational Analyses of Long-Read Sequencing Data

Guppy (v.6.4.2) high-accuracy mode was used for basecalling to generate the FASTQ files from the two libraries produced by ligation and rapid sequencing kits. The software NanoPlot (v.1.44.1)[13] was used to perform quality control and generate quality control relevant plots (**Supplementary Fig.5**). To identify chromosomal sequences on ecDNA, the FASTQ files were aligned to genome GRCh38 using minimap2[14] (v.2.26-r1175) with the command “minimap2 -cx map-ont”. From the resulting PAF file, reads from each chromosome were quantified. Further, for specific chromosomal regions, chr8: 125000000-130000000 and chr17: 39000000-40500000, normalized read coverage was calculated with ±1,000,000bp around these two specific regions. The results were saved in DAT files for coverage visualization shown in **Fig.3b** and **Supplementary Fig.6**.

To construct the consensus ecDNA structure, Flye (v.2.9.2)[15], a software designed for de novo assembly, was used on the FASTQ file generated by the ligation-based protocol with parameters “–nano-raw -g 4m -asm-coverage 40”. The potential ecDNA structure was stored in the resulting GFA (Graphical Fragment Assembly) file, which was further visualized using Bandage (v.0.8.1)[16]. In Bandage, BLAST (Basic Local Alignment Search Tool) search was done using the FASTA files of genes that are potentially amplified on ecDNA. Gene sequence FASTA files were downloaded from the NCBI database. After comparison, we found that the FASTQ data from the ligation sequencing library had better performance in terms of re-constructing the dominant ecDNA circle than the FASTQ data from the rapid sequencing library, which makes sense given the potentially higher sequencing accuracy and coverage provided by the ligation sequencing kit.

To detect variants, the FASTQ file was aligned to genome GRCh38 using miminap2 with the command “miminap2 -ax map-ont”. The resulting SAM file was used to generate the BAM file using samtools (v.1.21). Then the BAM file was sorted and indexed. The software Clair3[17] was used to call variants on the ONT long-read sequencing data. Clair3 was run within a Singularity container. The container image was pulled from a pre-built Docker image. The Clair3 model “r941_prom_hac_g360+g422” was used for variant calling.

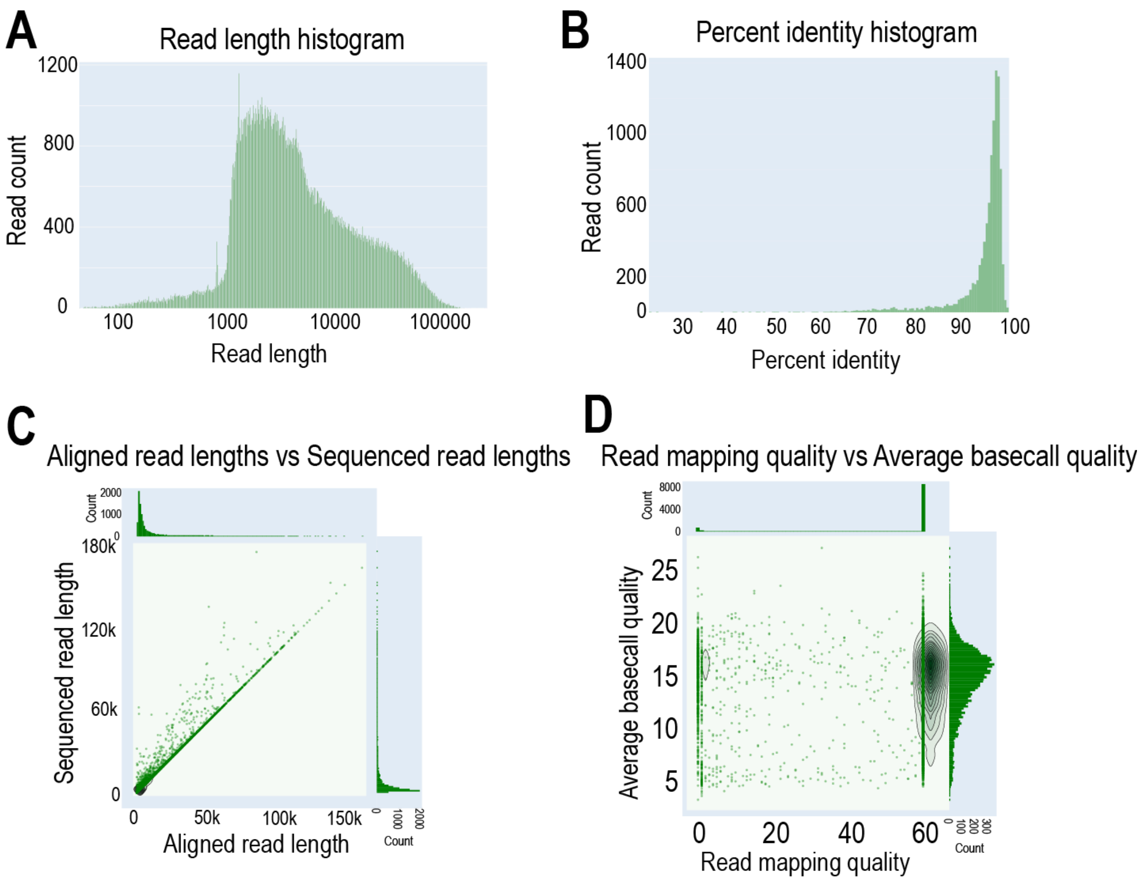

**Supplementary Fig.5: Quality control plots for NCI-H2170 ONT long-read sequencing. A.** ONT long-read sequencing Read length distribution **B.** Percent identity distribution across reads **C.** Visualization of aligned read length vs sequenced read lengths **D.** Read mapping quality vs average baseball quality

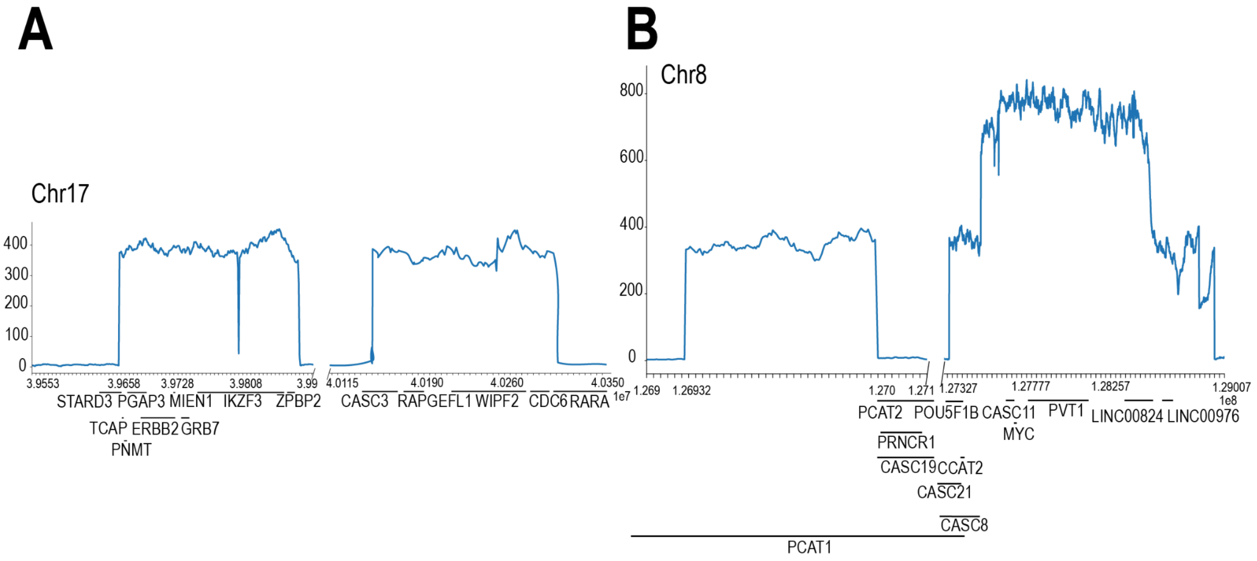

**Supplementary Fig.6: Coverage plots for ecDNA chr17 and chr8 regions from the ONT long-read rapid sequencing library. A.** Normalized chr17 ecDNA region coverage plot **B.** Normalized chr8 ecDNA region coverage plot. These two plots confirm the 2:1 copy number ratio of MYC to ERBB2

#### 2.5 Computational Analyses of Bioskryb ResolveOME sequencing data (scRNA-seq + scDNA-seq)

The Bioskryb Genomics BaseJumper bioinformatics platform (v.1.14), which covers pre-processing, quality control, read alignment, and downstream analysis, was used to analyze NCI-H2170 ResolveOME data. Genome GRCh38 was used for alignment.

The pipeline BJ-DNA-QC (v.2.0.3) was used for DNA data processing and analysis. In the BJ-DNA-QC pipeline, the following tools are implemented: Seqtk (v.1.3-r106), fastp (v.0.20.1), FastQC (v.0.11.9), Sentieon (v.202308.01), QualiMap (v.2.2.2-dev), Preseq (v.2.0.3), Kraken2 (v.2.1.3), bam-lorenz-coverage (v.2.3.0 GNU), Ginkgo (v.0.0.2), and bedtools (v.2.28.0). DNA FASTQ files with more than 10K reads were sent into the pipeline.

The software Ginkgo[18] was used for ecDNA copy number amplification analysis and the following parameters were chosen: bin_size=1,000,000bp; min_ploidy=1.5; max_ploidy=6; is_haplotype=2; n_reads=2000000. After running the pipeline, for each single cell, the normalized coverage for each chromatin bin is provided in the "SegNorm" slot in a RDS file. The bin "chr8-127412678-128444056" that covers the MYC gene was used to evaluate ecDNA counts across single cells given that it has the maximum overlap with known ecDNA sequence across all the bins.

From the DepMap bulk copy number data, the mean copy number (CN) across all genes in NCI-H2170 is around 1, which means the genes with a relative CN of 1 don't have any amplification or deletion relative to its ploidy and the exact CN of these genes are 1*NCI-H2170 ploidy= NCI-H2170 ploidy. Therefore, it is speculated that in Bioskryb scWGS data, the same thing holds true. In each single cell, the mean normalized coverage for all bins is the coverage for chromosome regions that don't have amplification or deletion relative to its ploidy. Since from G-band Karyotyping the ploidy of NCI-H2170 is roughly 4.5 (the average number of triploid and hexaploid), the exact CN for these regions is 4.5. Then if the copy number of the bin "chr8-127412678-128444056" is A, the normalized coverage for this bin is B, the mean normalized coverage for all bins is C, we have A/B=4.5/C. Since the values of B and C for each single cell are available, we can calculate the copy number of the bin "chr8-127412678-128444056", which is 4.5*B/C. From ONT long-read sequencing, we know that the dominant ecDNA species has two MYC and one ERBB2, therefore the dominant ecDNA species count in each cell would be 0.5*4.5*B/C.

For variant detection, The Genome Analysis Toolkit (GATK)[19] (v.4.6.1.0) was used on the concatenated pseudobulk WGS data for all single cells including FACS-sorted high, medium and low cells. The function "gatk HaplotypeCaller" was used to call the variants. The output variants were filtered according to GATK hard filtering basic thresholds. Due to the coverage limitation, we focused on ecDNA regions since this highly amplified region was able to receive enough coverage (>30X) in the sparse pseudobulk WGS data. The analysis-ready variants were annotated using the Genome Aggregation Database (gnomAD). For variant detection in FACS-sorted HER2 "high" and "low" cells, the same workflow was used on the concatenated pseudobulk WGS data for the FACS-sorted HER2 "high" and "low" cells respectively. To test of there is any mutation burden difference between ecDNA-high vs ecDNA-low cells, the pseudobulk WGS data for the FACS-sorted HER2 "high" condition was randomly downsampled for 3 replicates (Seed1, Seed2, and Seed3) (**Supplementary Fig.7-8**) so that the pseudobulk WGS data for FACS-sorted HER2 high and low conditions has the same coverage in the ecDNA regions.

The pipeline BJ-Expression (v.1.8.3) was used for scRNA analysis. In the BJ-Expression pipeline, the following tools are implemented: Seqtk (v.1.3-r106), fastp (v.0.20.1), Salmon (v.1.6.0), STAR (v.2.7.6a), QualiMap (v.2.2.2-dev), Samtools (v.1.10), and HTseq (v.0.13.5). RNA FASTQ files with more than 10K reads were sent into the pipeline. Raw count outputs from Salmon were used to calculate counts per million (CPM) and transcripts per million (TPM) for further analysis. In the data filtering step, cells with no more than 1000 genes were filtered out and genes present in less than 50% of cells were filtered out. A Seurat object was created for the normalized TPM data using the Seurat library (v.4.4.0), and the inferred ecDNA counts were added to the seurat object as metadata. Differential gene expression analyses for the FACS-sorted “high” vs “low” groups and the inferred ecDNA count “high” vs “low” groups were performed using the FindMarkers() function in the Seurat library. The Pearson correlation was calculated between inferred ecDNA counts and each gene expression for all the genes across single cells using the function cor.test() in R. Bonferroni correction was performed across all genes to calculate adjusted p-values.

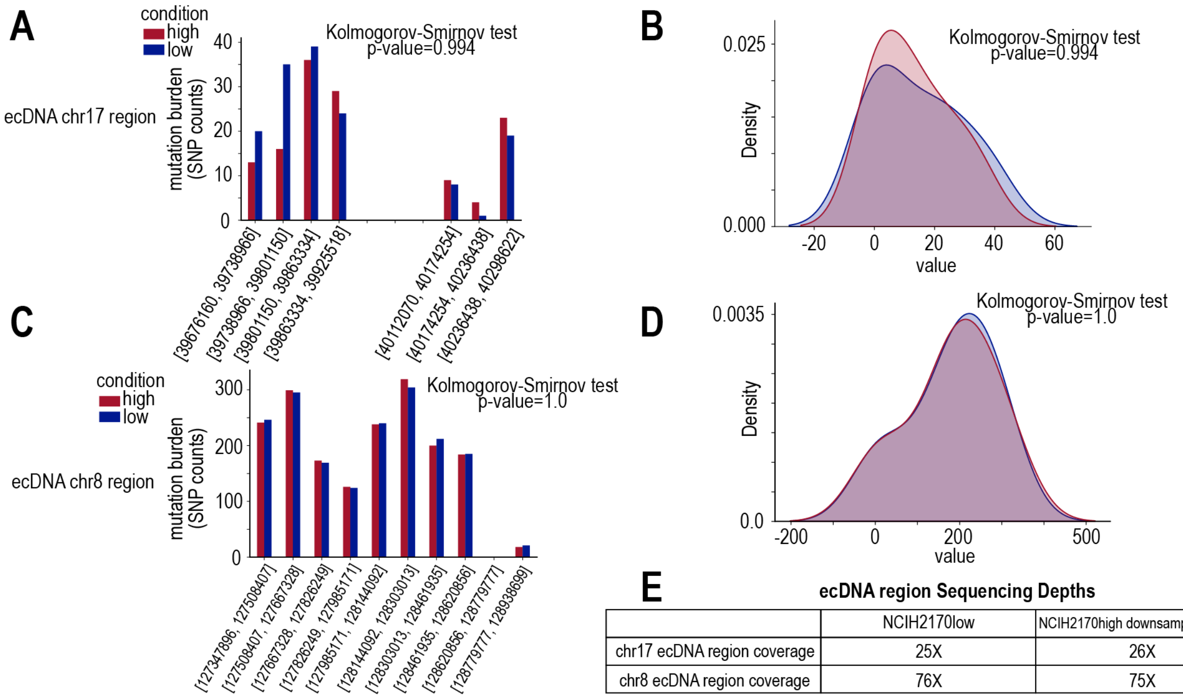

**ecDNA region Sequencing Depths**

| | NCIH2170low | NCIH2170high downsampling seed2 |
|---|---|---|
| chr17 ecDNA region coverage | 25X | 26X |
| chr8 ecDNA region coverage | 76X | 75X |

**Supplementary Fig.7: Mutation burden comparison between ecDNA high and low cells (ecDNA high cells random downsampling seed No.2) A.** SNP counts over different bins in ecDNA chr17 region for FACS-sorted high HER2 and low HER2 cells. **B.** Distribution comparison between SNP counts in FACS-sorted high HER2 and low HER2 cells in ecDNA chr17 region. **C.** SNP counts over different bins in ecDNA chr8 region for FACS-sorted high HER2 and low HER2 cells. **D.** Distribution comparison between SNP counts in FACS-sorted high HER2 and low HER2 cells in ecDNA chr8 region. **E.** Sequencing depths for ecDNA chr17 and chr8 regions in FACS-sorted high HER2 and low HER2 cells.

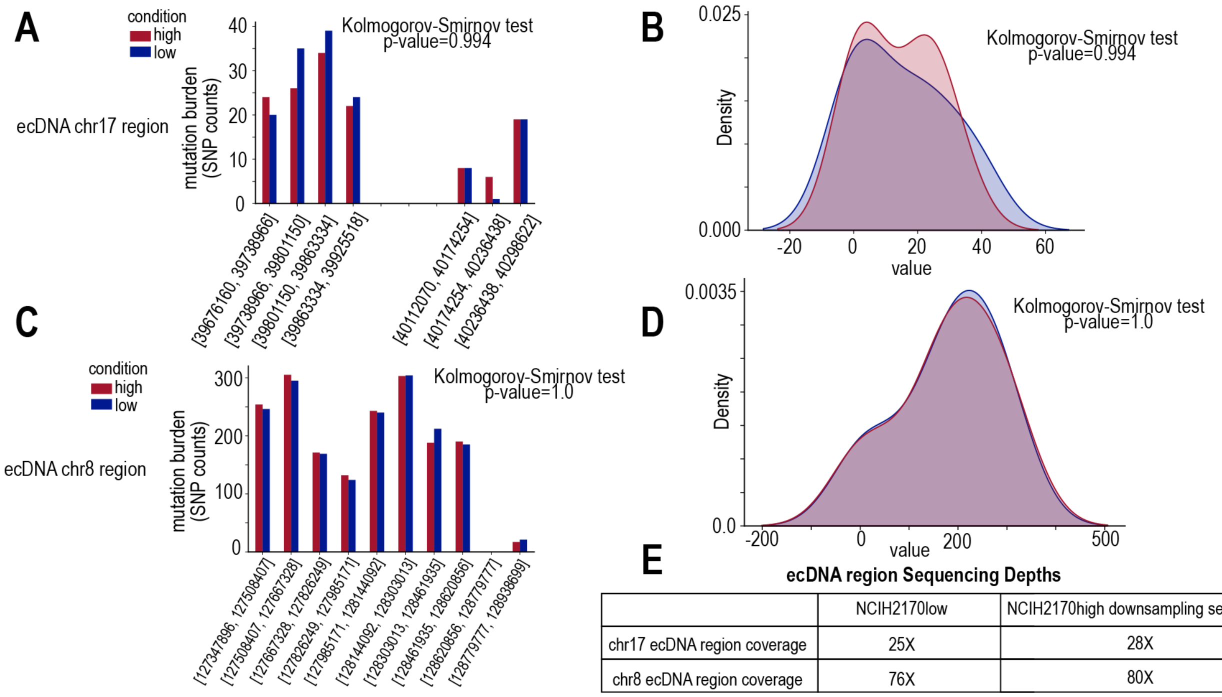

**E**

**ecDNA region Sequencing Depths**

| | NCIH2170low | NCIH2170high downsampling seed3 |
|---|---|---|
| chr17 ecDNA region coverage | 25X | 28X |
| chr8 ecDNA region coverage | 76X | 80X |

**Supplementary Fig.8: Mutation burden comparison between ecDNA high and low cells (ecDNA high cells random downsampling seed No.3) A.** SNP counts over different bins in ecDNA chr17 region for FACS-sorted high HER2 and low HER2 cells. **B.** Distribution comparison between SNP counts in FACS-sorted high HER2 and low HER2 cells in ecDNA chr17 region. **C.** SNP counts over different bins in ecDNA chr8 region for FACS-sorted high HER2 and low HER2 cells. **D.** Distribution comparison between SNP counts in FACS-sorted high HER2 and low HER2 cells in ecDNA chr8 region. **E.** Sequencing depths for ecDNA chr17 and chr8 regions in FACS-sorted high HER2 and low HER2 cells.

#### 2.6 Machine Learning of ecDNA Copy Number per Cell

*Data Preprocessing*

Single-cell RNA sequencing (scRNA-seq) data was used to predict inferred extrachromosomal DNA (ecDNA) copy number counts. The dataset contained 2600 gene expression features (log-transformed transcript-per-million values, logTPM) for each single cell. Prior to model training, the following preprocessing steps were applied. First, only the 2600 genes present in both the training and new prediction datasets were retained. Missing values were handled using median imputation via the SimpleImputer function in scikit-learn. Gene expression values were then standardized using StandardScaler() to ensure zero mean and unit variance. The dataset was split into 80% training and 20% test data using train_test_split with random_state=42 for reproducibility. Model performance was evaluated using mean absolute error (MAE), $R^2$ score, Pearson correlation, and Spearman correlation.

*Machine Learning Models*

To predict ecDNA copy number, we implemented four different models: Random Forest (RF), Gradient Boosting (GB), Extreme Gradient Boosting (XGBoost), and a deep neural network (NN).

The Random Forest (RF) model was trained using the RandomForestRegressor function with 500 decision trees (n_estimators=500), a maximum depth of 15 (max_depth=15), and a minimum of five samples required to split an internal node (min_samples_split=5). The minimum number of samples required in a leaf node was set to two (min_samples_leaf=2).

The Gradient Boosting (GB) model was implemented using GradientBoostingRegressor, with 500 boosting stages (n_estimators=500), a learning rate of 0.05 (learning_rate=0.05), and a maximum tree depth of 5 (max_depth=5). Additional hyperparameters included row sampling (subsample=0.8) and constraints on splits (min_samples_split=5, min_samples_leaf=2).

The Extreme Gradient Boosting (XGBoost) model was trained using XGBRegressor with optimized hyperparameters determined via grid search. The final model consisted of 500 boosting rounds (n_estimators=500), a learning rate of 0.05 (learning_rate=0.05), a maximum tree depth of 6 (max_depth=6), and feature subsampling (colsample_bytree=0.8). Regularization was applied using L1 (reg_alpha=0.1) and L2 (reg_lambda=1) penalties.

The Neural Network (NN) model was implemented using PyTorch. The architecture consisted of an input layer with 2600 features, followed by three hidden layers with 512, 256, and 128 neurons, respectively. The Swish activation function (x * sigmoid(x)) was applied to all hidden layers. Dropout regularization (p=0.2) was used to prevent overfitting. The output layer consisted of a single neuron predicting ecDNA copy number. Training was performed using Huber loss (SmoothL1Loss()), with the AdamW optimizer (lr=0.00005, weight_decay=1e-4). The model was trained with a batch size of 64 and early stopping with a patience of 20 epochs.

*Model Evaluation and Selection*

All models were evaluated on the test dataset using MAE, $R^2$ score, Pearson correlation, and Spearman correlation. Among the four models, XGBoost demonstrated the best overall performance, achieving the lowest MAE (67.47) and the highest $R^2$ score (0.7276). The neural network model showed competitive performance in rank-based correlation, with a Spearman correlation of 0.7774. Given its strong predictive accuracy and interpretability, XGBoost was selected as the final model for predicting ecDNA copy number from scRNA-seq data.

*Application of XGBoost Model for New Data*

The trained XGBoost model was applied to new single-cell RNA sequencing datasets to infer ecDNA copy number. To ensure consistency, new datasets were processed identically to the training data. This included selecting the same 2600 genes, applying median imputation for missing values, and standardizing features using the previously trained StandardScaler. The trained XGBoost model was then loaded and applied to predict ecDNA counts. The predicted values were stored in a CSV file for downstream analysis.

To facilitate reproducibility, all model training and evaluation steps were implemented in Python 3.8+ using scikit-learn, XGBoost, PyTorch, and joblib. Random seeds were set to ensure consistent results. The full pipeline, including preprocessing scripts and trained model files, is available upon request.

#### 2.7 Statistical Analyses of Longitudinal FISH Data

To assess differences in ecDNA abundance across experimental groups and timepoints, we applied both linear mixed effects modeling and ordinary least squares (OLS) regression, depending on the structure and availability of data for each comparison.

*Global mixed effects model*

To estimate ecDNA changes across all groups and timepoints while accounting for batch effects, we fit a linear mixed effects model using the statsmodels Python package. The model was specified as:

**ecDNA ~ Timeline * Group + (1 | condition)**

where Timeline indicates the timepoint (e.g., G0, G1, G2, G9), Group reflects the FACS-sorted population (High, Low, Post-sort Control, Pre-sort Control), and folder was included as a random effect to account for variability across experimental batches. Baseline levels were explicitly set such that Post-sort Control at G0 served as the reference group. This allowed for direct interpretation of fixed effect coefficients as differences from this baseline. In cases where Post-sort Control samples were not available for a particular time point (e.g., G9), synthetic rows were generated by duplicating G2 Post-sort Control data and re-labeling as G9. Model fitting was performed using maximum likelihood estimation (reml=False) with the L-BFGS optimizer.

*Subset comparisons using OLS*

To specifically evaluate differences between High vs. Low populations at individual timepoints, we subset the dataset to include only samples from the relevant timeline and group. For each timepoint (G0, G1, G2, G9), we fit an OLS model of the form:

**ecDNA ~ Group**

with High set as the reference level. This allowed us to directly quantify the difference in ecDNA abundance between Low and High cells at each timepoint.

To evaluate whether the group-level difference changed across timepoints (e.g., whether the High-Low difference at G9 differed from G2), we also fit interaction models of the form:

**ecDNA ~ Timeline * Group**

where Timeline and Group were both treated as categorical variables with specified reference levels.

*Inference and visualization*

For all models, we extracted estimated coefficients, standard errors, p-values, and confidence intervals. In select cases, we used t_test() with custom contrast vectors to compute p-values for predicted ecDNA levels at specific combinations (e.g., G9 High vs. baseline). Group means and standard errors were visualized using bar plots with error bars, and group differences were further evaluated using Welch's t-tests (unequal variances).

All analyses were conducted in Python using the statsmodels, scipy, pandas, and seaborn libraries.

#### 2.8 Stochastic Modeling of ecDNA Redistribution

##### 2.8.1 Comparison to Existing Stochastic/Evolutionary Models

Prior models of ecDNA dynamics, including studies in Nature (2017)[20], Nature Genetics (2022)[21], and Nature (2024)[7], have generally treated ecDNA as a passive feature of genomic instability. These frameworks rely on fixed probabilistic rules for segregation, proliferation, and selection. The 2017 Nature study, for example, used a Galton–Watson branching process to simulate oncogene amplification, imposing copy-number ceilings through logistic functions. While it provided insights into selection thresholds, the model assumed simplified population dynamics and did not account for how heterogeneity might be re-established after disruption.

More recent models simulated co-assortment of ecDNA species using binomial segregation and fixed selection coefficients. These forward-time simulations, which included parameter inference via Approximate Bayesian Computation, were valuable for evaluating co-selection trends and copy-number correlations. However, they operated under static assumptions about ecDNA inheritance and did not address how variation could be restored following a selective bottleneck.

To evaluate these approaches, we implemented their core assumptions in our simulator. In each case, we found that these models failed to reproduce the experimentally observed restoration of ecDNA heterogeneity within two to three generations. Instead, they predicted a monotonic loss of diversity or stable equilibria, without recovery of the original distribution.

##### 2.8.2 Novelty and Conceptual Advances of Our Approach

Our model offers a new conceptual framework for ecDNA population dynamics. Rather than assuming passive inheritance, we simulate the active restoration of ecDNA variation over time, using a generation-resolved, Gillespie-based stochastic birth-death process.

Key innovations include modeling division and death rates as non-linear functions of ecDNA copy number, enabling us to represent both fitness advantages at low levels and growth penalties at high levels. We also introduce a split inheritance parameter (r), which allows for asymmetric ecDNA segregation during mitosis. This feature is essential to capturing the rapid re-establishment of heterogeneity seen in our experiments and is jointly optimized alongside fitness parameters using a Genetic Algorithm. To initialize the model, we apply a Metropolis–Hastings algorithm to upsample ecDNA distributions from limited G0 (post-sort) data (see section 2.8.3). This KDE-based approach improves starting conditions and avoids sampling bias.

When we implemented prior modeling assumptions, such as symmetric segregation or fixed selection, we consistently failed to replicate the observed restoration of ecDNA variability. Only by combining asymmetric inheritance, dose-sensitive fitness effects, and parameter optimization were we able to reproduce both the timing and structure of ecDNA redistribution across generations.

Together, these innovations allow us to simulate not only the maintenance of ecDNA heterogeneity, but also its active recovery, a hallmark of ecDNA-driven adaptation that previous models could not explain.

##### 2.8.3 Data up-sampling with Metropolis Hastings

To reduce the effect of bias and extreme values caused by the small G0 data set, a Metropolis-Hastings scheme was used to up sample the data. We implemented this algorithm in Python. A KDE (Kernel Density Estimate) was created for each of the high, post-sort control, and low HER2 expression groups. For each expression group, new ecDNA counts were generated by comparing a candidate ecDNA count to the previously accepted ecDNA count. If a candidate count had a higher likelihood according to the KDE, it was automatically accepted. If it had a lower likelihood according to the KDE, it was accepted with probability $p(x')/p(x)$, where p is the KDE, x' is the candidate value, and x is the previous value. The first 100 values are removed to avoid initialization bias. For each of the HER2 expression groups (high, low, and post-sort control) 1000 new samples were generated and the distributional similarity was verified using a Kolmogorov-Smirnov Test ($p > 0.05$).

#### 2.8.4 Gillespie Algorithm

The Gillespie Algorithm, implemented in Python, was used to simulate cell growth stochastically while keeping a global pseudotime. First cellular event rates were summed based on the fitness-based events within each cell: either division or death. These were either static rates across each cell in the case of the neutral model, or ecDNA dependent rates in the case of the non-neutral models. In the latter case, the parameters for calculating the rates from the ecDNA counts were passed into the Gillespie Algorithm as a parameter. Once all the rates of the cells were calculated, the rates were summed to provide a global event rate. A global pseudotime was simultaneously initialized as t = 0.

An event time Δt is sampled from an exponential using the global event rate as a parameter. Then the event is chosen (either a division or a death) by randomly sampling a real number from a uniform distribution from 0 to the global event rate. If the randomly generated number is less than the sum of the division rates, the event is a division, otherwise the event is a death. Finally, the cell for which that event happens is determined by drawing a random real number between 0 and either the sum of the death or division rates (depending on the previous step). The chosen cell is determined by partitioning the reals between 0 and the death or division rate into segments based on the individual cell rates and checking which cell rate interval the randomly generated number falls into.

Once an event and a cell is chosen, the event occurs and Δt is added to the global pseudotime. In the case of a death event, this removes the cell's death and division rate from the global event rate and no longer keeps track of the ecDNA within the cell (they are effectively destroyed). In the case of a division, an integer is drawn from a binomial distribution with the number of the trials corresponding to twice the number of ecDNA. Then the global event rates are updated based on the division and death rate of the new cells. This process repeats using the updated event rates until the global pseudotime hits the maximum amount of time (48 hours).

#### 2.8.5 Genetic Algorithm

The Genetic Algorithm was used to optimize the parameters for cell death rates, cell division rates, and the split probability in the binomial distribution, even when the high dimensional space is non-convex. This code was written in Python. 2000 parameter sets were generated from uniform distributions of parameters related to the cell death, cell division, and split probability. Each of these were then loaded onto a separate CPU and the Gillespie Algorithm was run for 7 generations (336 hours) under each parameter. The distribution determined after each generation and an objective score was calculated for all 2000 parameter sets based on the KL-divergence between the total ecDNA distribution of subsequent generations (Steady-State objective) and the KL-divergence between the HER2 expression groups within each generation after G0 (Recentralization objective). The top 20 parameter sets based on this objective were collected from among the 2000 and returned to a single CPU.

2000 new parameter sets were then created by randomly sampling parameters from the top 20 parameter sets and randomly mutating each within 5% of its original value. This process was repeated on this set of 2000 new parameters. This repetition was done for 2000 iterations or until the maximum objective scores plateaued. The parameter set optimized by the GA are as follows:

- A1: 0.053
- A2: 156
- A3: 516
- B1: 0.008
- B2: 235
- B3: 187
- B4: 0
- r: 0.26

#### 2.9 AI-assisted ecDNA counting of FISH images

##### 2.9.1 Overview

The automated pipeline for detecting and quantifying extrachromosomal DNA (ecDNA) in Fluorescence in situ Hybridization (FISH) images was developed to process paired RGB FISH images (labeled with probes such as HER2 or MYC) and DAPI grayscale images. The pipeline consists of image preprocessing, enhancement, object detection, classification, hyperparameter optimization, validation, batch processing with external comparison, and data management.

Each step was implemented in Python (version 3.10.9) on the Longleaf cluster. The pipeline utilizes standard libraries including OpenCV (4.11.0.86), NumPy (1.26.4), pandas (2.1.4), Matplotlib (3.8.4), Seaborn (0.13.2), SciPy (1.13.1), and bayesian-optimization (2.0.3). All dependencies were version-controlled, a dedicated virtual environment (ecDNA_env) was employed, and reproducibility was ensured by setting a fixed NumPy random seed (10).

The following subsections detail each component of the pipeline, including its purpose, implementation, parameters, and outputs.

##### 2.9.2 Image Processing

The initial preprocessing step isolates nuclear regions within RGB FISH images to ensure that subsequent analyses focus on one cell and to reduce background noise. The DAPI image, which highlights nuclear material, was used to create a mask: pixels with zero intensity in the DAPI image (indicating non-nuclear regions) were set to zero in the RGB image, excluding irrelevant areas effectively. This process was performed using OpenCV with NumPy libraries. Following masking, the RGB image was converted to grayscale using OpenCV's cv2.cvtColor function with the cv2.COLOR_BGR2GRAY flag, to simplify the image for further processing.

*Image Enhancement*

The image enhancement step amplifies the visibility of small, bright ecDNA signals while suppressing larger chromosomal structures that could interfere with detection. The masked grayscale image undergoes a four-stage enhancement process, with each stage controlled by parameters optimized through Bayesian Optimization.

First, a morphological top-hat transformation was applied using OpenCV's cv2.morphologyEx function with an elliptical structuring element defined by the kernel_size parameter. This step enhances small, bright objects like ecDNA by subtracting the background, while a larger kernel (chrom_kernel_size) estimates chromosomal regions, which are then dampened by a dampening_factor to reduce their intensity.

Second, a sharpening filter was applied using a high-pass kernel scaled by the strength parameter, enhancing the edges of potential ecDNA objects.

Third, Contrast Limited Adaptive Histogram Equalization (CLAHE) was performed with OpenCV's cv2.createCLAHE, using clip_limit and tile_grid_size to improve local contrast and highlight ecDNA signals.

Finally, a sigmoid transformation adjusted pixel intensities with cutoff and gain parameters, further boosting the visibility of ecDNA.

*Object Detection*

The object detection step detects potential ecDNA and chromosome objects within the enhanced grayscale image. The process begins with binarization using Otsu's thresholding method in OpenCV, which separates objects from the background by automatically determining an optimal threshold.

To refine the binary image, morphological cleaning was applied: opening and closing operations with a "2x2" elliptical kernel were used to remove noise and connect fragmented objects, ensuring that ecDNA signals are not split into multiple parts. Connected components analysis was then performed using OpenCV's cv2.connectedComponentsWithStats, which labels and extracts objects along with their bounding boxes, centroids, and areas.

Objects were filtered based on their area, keeping only those between min_area and max_area to exclude noise and overly large artifacts. To address potential over-segmentation, objects closer than a merge_distance (measured by centroid distance using Python's math.dist function) were merged into a single object by combining their bounding boxes and areas.

*Object Classification*

Object classification categorizes the detected objects as either ecDNA or chromosomes based on their color properties in the original RGB image. For each detected object, a region of interest (ROI) was extracted from the RGB image using the object's bounding box coordinates. The ROI was converted to the HSV color space with OpenCV's cv2.cvtColor function and the cv2.COLOR_BGR2HSV flag, since HSV better separates brightness and color information.

The mean value (V) and saturation (S) of the HSV ROI were computed using OpenCV's cv2.mean function. An object was classified as a "chromosome" if its mean value exceeded the white_value_threshold and its mean saturation fell below the white_saturation_threshold, reflecting the typically white appearance of chromosomes in FISH images; otherwise, it was classified as "ecDNA," which appears as colored spots due to FISH probes.

##### 2.9.3. Hyperparameter Optimization

Hyperparameter optimization was conducted to tune the pipeline's parameters for optimal ecDNA detection accuracy. Bayesian Optimization was employed using the bayesian-optimization library, to minimize the Median Absolute Percentage Error (MdAPE) between predicted and ground truth ecDNA counts. The optimized parameters include kernel_size, chrom_kernel_size, and dampening_factor for top-hat filtering; clip_limit, tile_grid_size, strength, cutoff, and gain for enhancement; merge_distance, min_area, and max_area for detection; and white_value_threshold and white_saturation_threshold for classification. The optimization process involved fifteen initial random evaluations followed by eighty-five iterations, totaling 100 evaluations, to explore the parameter space efficiently. The resulting optimized parameters were saved in a JSON file (best_params.json) and used for all future processing.

*Validation*

Validation and comparison were performed in a two-stage process to ensure robust assessment of the pipeline's performance against manual counting and predictions from the MIA method. The dataset of 388 FISH images with ground truth ecDNA counts was initially split into a training set (300 images) and a test set (88 images) to evaluate the consistency of optimization results across different image subsets.

In the first validation stage, hyperparameter optimization was conducted using only the training set, minimizing the Median Absolute Percentage Error (MdAPE) between predicted and ground truth ecDNA counts. The process involved 15 initial random evaluations followed by 85 directed iterations (100 total evaluations), efficiently exploring the parameter space. When the resulting parameters were applied to the test set, consistent performance was observed, confirming the robustness of the optimization approach.

Following this confirmation of consistency, a second optimization was performed on the full dataset of 388 images to determine parameters that would provide optimal performance across all available data. Notably, the hyperparameters resulting from both optimization processes (training set only versus full dataset) were remarkably similar, further validating the stability of the approach. The final optimized parameters (**Supplementary Table.3**) were saved and used for all subsequent processing.

**Supplementary Table 3. Optimized Hyperparameters for the ecDNA Detection Pipeline**

| Parameter | Value | Parameter | Value |
|---|---|---|---|
| kernel_size | 10.83 | chrom_kernel_size | 168.9 |
| dampening_factor | 0.3346 | clip_limit | 3.95 |
| tile_grid_size | 21.71 | strength | 4.666 |
| cutoff | 68.44 | gain | 23.14 |
| merge_distance | 8.013 | min_area | 4.133 |
| max_area | 619.6 | white_value_threshold | 179.5 |
| white_saturation_threshold | 63.32 | | |

Comparison with MIA[22]

Using these optimized parameters, the pipeline's predictions were compared to both ground truth manual counts and MIA predictions, calculating several metrics: MdAPE (median of per-image absolute percentage errors), Mean Absolute Error (MAE), trimmed MAE (after removing the top and bottom 5% of errors due to presence of outliers), and Pearson correlation between predicted and true counts. A Wilcoxon signed-rank test, implemented with SciPy, was used to statistically compare the pipeline's performance to MIA.

Results were visualized using scatter plots and box plots (on a logarithmic scale) with Seaborn and Matplotlib, showing predicted versus ground truth counts and error distributions. The pipeline achieved an MdAPE of 13.31%, MAE of 37.60, trimmed MAE of 31.62, and a correlation of 0.94, significantly outperforming MIA (MdAPE: 47.88%, MAE: 112.88, correlation: 0.72; p < 10^-29), demonstrating its superior accuracy and consistency in ecDNA detection.

#### 2.9.4. Batch Processing

Batch processing was implemented to apply the pipeline to larger datasets and compare its predictions with MIA outputs, enabling a comprehensive assessment of its performance. In one replication a total of 1141 FISH images were processed in parallel using Python's multiprocessing.Pool with eight workers to optimize computational efficiency. Each image underwent the full pipeline—preprocessing, enhancement, detection, and classification—and results were saved as intermediate debug images (e.g., masked RGB, enhanced grayscale, annotated overlays) for manual validation.

MIA-predicted masks were segmented using connected components analysis with OpenCV, and objects were matched to the pipeline's detections using Intersection over Union (IoU) thresholds (iou_full=0.01, iou_partial=0.001) and a centroid distance_threshold of 30.0 pixels. Matching metrics—True Positives (TP), Partial TP, False Negatives (FN), and False Positives (FP) were computed, with the pipeline detecting an average of 211.84 objects per image compared to MIA's 82.94 (average TP: 37.53, TP_partial: 29.94, FN: 144.37, FP: 15.47).

Data Management and Reproducibility

Data management and reproducibility were prioritized to ensure organized output and accessibility for future analyses. Output files, including debug images, were automatically organized into subfolders based on image metadata (e.g., G0/High_HER2, G1/Low_HER2, G2/Total_HER2), with summary files (metrics_summary.csv, updated_ecDNA_counts.csv, ref_objects.json) moved to a Summary folder using Python's shutil module.

All dependencies were pinned in a requirements.txt file for reproducibility. A virtual environment (ecDNA_env) was used to manage these dependencies, ensuring a consistent computational environment. A random seed of "10" was set for NumPy to ensure consistent results across runs. The pipeline's outputs are structured to facilitate validation, sharing, and reuse by other researchers, supporting transparency and reproducibility in ecDNA studies.
